# Metabolite profiling reveals slow and uncoordinated adjustment of C_4_ photosynthesis to sudden changes in irradiance

**DOI:** 10.1101/2025.04.13.648559

**Authors:** Stéphanie Arrivault, David Barbosa Medeiros, Cristina Rodrigues Gabriel Sales, Manuela Guenther, Johannes Kromdijk, Alisdair R. Fernie, Mark Stitt

## Abstract

In the field plants continually experience changes in irradiance. Research in C_3_ species has revealed that whilst the Carbon-Benson cycle (CBC) adjusts rapidly to changing irradiance, there is substantial loss of photosynthesis efficiency due to slow adjustment of energy dissipation and stomatal conductance. Less is known about the impact of changing irradiance on photosynthetic efficiency in C_4_ species. We have subjected maize to a sudden increase or decrease of irradiance in the non-saturating range and performed time-resolved measurement of photosynthetic rate and profiling of metabolites from the CBC, the CO_2_-concentrating mechanism (CCM), photorespiration and end-product biosynthesis. After a decrease in irradiance, photosynthesis is transiently buffered by energy delivered by transformations in the large pools of metabolites that are involved in intercellular shuttles. During the subsequent decline in photosynthesis, metabolism transitions to a sub-optimal state for photosynthesis in low irradiance, from which it takes several minutes to recover. One reason is that the large pools of metabolites that facilitate intercellular shuttles run down too far and it takes time to build them up again. After an increase in irradiance, there is a long delay until a new steady rate of photosynthesis is achieved. The delay is partly due to regulation of enzymes in the CBC and CCM, but also to the need to build up large pools of metabolites to drive intercellular shuttles. In addition, in both transitions, transient imbalances between the pumping and utilization of CO_2_ lead to increased back-leakage of CO_2_ or to photorespiration, further decreasing photosynthetic efficiency.

## Introduction

Plants in the field experience daily changes of irradiance due to the elevation of the sun and shadowing by fixed objects as well as irregular changes due to changing cloud cover and wind-movement in canopies (Townsend et al. 2018). The response to fluctuating irradiance is an important factor for photosynthetic efficiency in the field, especially in dense crop canopies (Tayler and Long 2017; Vialet-Chabrand et al. 2017; Townsend et al. 2018; Tanaka et al. 2019; Wu et al. 2019).

The response of photosynthetic carbon (C) metabolism in C_3_ species to fluctuating irradiance was investigated in the last century (Pearcy 1990; Peary et al. 2005; Mott and Woodrow 2000). Wild C_3_ species that live in the understorey of woods and forests where occasional sun flecks provide a large part of total intercepted irradiance show an extreme adaptation. In low light their Calvin-Benson cycle (CBC) is poised to maintain a very large pool of 3-phosphoglycerate (3PGA), allowing full use of the NADPH and ATP that is produced during a brief sun fleck without this requiring an increase in flux around the CBC (Pearcy 1990; Pearcy et al. 2005). In most C_3_ species, it typically takes 1-2 min to establish steady state CBC flux after a switch from darkness or low light to high light (Woodrow and Walker 1980; Laing et al. 1981; Wirtz et al. 1982; Woodrow et al. 1983, 1985; Sage et al. 1987; Mott and Woodrow 2000). During this short lag, CBC metabolite levels rise and CBC enzymes are post-translationally activated. In the case of Rubisco, the speed of activation depends on the abundance of Rubisco activase (Mott et al. 1997; Woodrow et al. 1996; Hammond et al. 1998), with a trade-off between the abundance of activase, which determines how quickly Rubisco activation responds to a change in irradiance, and of Rubisco which impacts on the rate of photosynthesis under stable irradiance (Woodrow and Mott 1989; Yamori et al. 2012; Carmo-Silva and Salvucci 2013; Kaiser et al. 2016). Post-translational activation of plastidic fructose 1,6-bisphosphatase (FBPase) and sedoheptulose 1,7-bisphosphatase (SBPase) is promoted by rising substrate levels, which promote their reduction by thioredoxin (Woodrow and Walker 1980; Laing et al. 1981; Woodrow et al. 1985; Faske et al. 1995; Scheibe 1991; Stitt et al. 2010; Michelet et al. 2013; Knuesting and Scheibe 2018). In addition to regulation within the CBC, regulation of end-product synthesis poises CBC metabolites at levels that allow a rapid rise in CBC flux when irradiance suddenly increases (Stitt et al. 2021). After a decrease in irradiance the response of the CBC is dominated by the extremely rapid turnover of the small pools of NADPH, ATP (about 0.1 s) and CBC intermediates (0.1-1 s) (Stitt et al. 1980; Arrivault et al. 2009). An almost instantaneous decline of NADPH and ATP arrests 3PGA reduction, and within 1-5 s falling levels of triose-phosphate (triose-P) and other CBC intermediates restrict ribulose-1,5-bisphosphate (RuBP) regeneration and CO_2_ fixation (Prinsley et al. 1986a; 1986b; Stitt et al. 1989). CBC enzymes are inactivated over the next minutes (Woodrow and Walker 1980; Laing et al. 1981; Wirtz et al. 1982; Woodrow et al. 1983; 1985; Sage et al. 1987; Woodrow and Berry 1988; Mott and Woodrow 2000). Enzyme inactivation in low irradiance is important to avoid energy wastage in futile cycles (Stitt et al. 2021). Overall, in C_3_ plants CBC flux adjusts rather quickly to changes in irradiance. The research focus has shifted to processes that adjust more slowly and have a larger impact on photosynthetic performance in fluctuating light such as energy dissipation in the light reactions (Kromdijk et al. 2016; Taylor and Long 2017; Tanaka et al. 2019; Wu et al. 2019) and stomatal conductance (Lawson et al. 2012; Vialet-Chabrand et al. 2017; Deans et al. 2019).

Less is known about the response of C_4_ photosynthesis to fluctuating irradiance. C_4_ photosynthesis evolved multiple times in response to selective pressure imposed by a change in the climate including a drop in the atmospheric CO_2_ concentration from about 1000 ppm to 250 ppm or below about 30 million years ago (Christin et al. 2008, 2011; Edwards et al. 2010). This decline exerted strong selective pressure on the CBC because Rubisco catalyses a competing side reaction with O_2_ (Lorimer 1981; Tcherkez et al. 2006) leading to formation of 2-phosphoglycolate (2PG) that has to be removed by the energetically wasteful process of photorespiration (Osmond 1981; Foyer et al. 2009; Walker et al. 2016; Betti et al. 2016). In C_4_ photosynthesis this process is partly suppressed by a biochemical CO_2_-concentrating mechanism (CCM) (Osmond and Harris 1971; Hatch, 2002; von Caemmerer and Furbank 2003; Sage et al. 2011; Schlüter and Weber, 2020). The CCM starts in the mesophyll cells (MC) with incorporation of bicarbonate by phospho*enol*pyruvate carboxylase (PEPC) to form 4-carbon (4C) metabolites. These move to the bundle sheath cells (BSC) where they are decarboxylated to generate a high concentration of CO_2_ (Hatch and Osmond 1976; Edwards and Walker 1983; von Caemmerer and Furbank, 2003; Weber and von Caemmerer 2010; Schlüter and Weber 2020). This CO_2_ is utilised by Rubisco which together with the rest of the CBC is located in the BSC. The 3-carbon (3C) products of decarboxylation move back to the MC, where they are used to regenerate phospho*enol*pyruvate (PEP). Intercellular shuttling of 4C and 3C metabolites is thought to occur largely by diffusion (Hatch and Osmond 1976), requiring large pools to generate the necessary intercellular concentration gradients (Leegood 1985; Stitt and Heldt, 1985a; Arrivault et al. 2017).

Decarboxylation can occur via NADP-malic enzyme (NADP-ME) in the chloroplast, NAD-malic enzyme (NAD-ME) in the mitochondria as well as PEP carboxykinase (PEPK) in the cytosol. Which of these decarboxylases controls the major part of C_4_ acid decarboxylation flux differs between species and affects which 4C and 3C metabolites move between the two cell types as well as energy requirements in the MC and BSC (Hatch and Osmond 1976; Edwards and Walker 1983; Hatch 2002; Bräutigam et al. 2018). Decarboxylation by NADP-ME is linked to a malate/pyruvate shuttle, whereas decarboxylation by NAD-ME or PEPCK requires exchange of other metabolites including aspartate, alanine and possibly PEP. Furthermore, decarboxylation via NADP-ME delivers NADPH to BSC chloroplasts at a rate that covers about half the NADPH required by the CBC. Many NADP-ME species including maize have dimorphic BSC chloroplasts with strongly decreased levels of photosystem II and NADPH production (Woo et al. 1970; Laetsch 1974; Munekage 2016). The resulting shortfall in NADPH is compensated by a second intercellular ‘energy’ shuttle in which 3PGA moves to the MC, where it is reduced to triose-P that returns to the BSC (Leegood 1985; Stitt and Heldt 1985; Arrivault et al. 2017). Intercellular movement again occurs by diffusion and requires large pools of 3PGA and triose-P to generate intercellular concentration gradients (Leegood 1985; Stitt and Heldt 1985a; Arrivault et al. 2017).

Efficient operation of C_4_ photosynthesis requires coordinated flux in the CCM and the CBC (Furbank et al. 1990; von Caemmerer 2000, Kromdijk et al. 2014). If CO_2_ transfer is too slow, the CO_2_ concentration in the BSC (C_BSC_) will fall and photorespiration will increase. C_4_ plants have substantial activities of enzymes for photorespiration (Osmond and Harris 1971; Ohnishi and Kanai 1983; Ueno et al. 2005) and still carry out photorespiration, although at 5-to 10-fold lower rates than those typical for C_3_ photosynthesis (Volk and Jackson 1972; de Veau and Burris 1989; Laisk and Edwards 1998; Carmo-Silva et al. 2008; Mallmann et al. 2014, Weissmann et al. 2016; Arrivault et al. 2017; Medeiros et al. 2022). On the other hand, excessive CO_2_ transfer will drive C_BSC_ up, leading to faster back-leakage of CO_2_ to the MC. It has been estimated that 13-30% of the CO_2_ released in the BSC leaks back to the MC (Farquhar 1983; Hatch et al. 1995; Bellasio and Griffiths 2014b; Kromdijk et al. 2014, Wang et al. 2022) with a corresponding wastage of energy.

Even in stable conditions, C_4_ photosynthesis is less efficient in low irradiance than high irradiance. For example, quantum yield of C_4_ photosynthesis has been reported to decrease at low irradiance (Pengelly et al. 2010; Pignon et al. 2017; Tazoe et al. 2008; Ubierna et al. 2013; Sales et al. 2023). Recently, it was shown by analysis of ^13^CO_2_ labeling kinetics (Medeiros et al. 2022) that photosynthetic efficiency is impaired at low irradiance due to a combination of factors including excess flux at PEPC and back-leakage of CO_2_ from the BSC to the MC (see also Ubierna et al. 2013; Kromdijk et al. 2014), restriction of decarboxylation by NADP-malic enzyme and a switch towards other decarboxylation routes, less effective use of metabolite pools to drive intercellular shuttles, and higher rates of photorespiration.

In addition to this inherent inefficiency in low irradiance, its complex topology may render C_4_ photosynthesis vulnerable to sudden changes in irradiance Firstly, NADPH and ATP are consumed by numerous reactions in two pathways, the CCM and CBC, which are distributed between two cell types, each of which has its own photosynthetic electron transport systems. Sudden changes in irradiance may lead to local imbalances in redox and energy status that, in turn, unbalance fluxes in the CCM and CBC. Secondly, the temporal dynamics of metabolite pools differ greatly between the CBC and the CCM. CBC metabolite pools are small and have half-times of the order of <0.1 to 1 s (Stitt and Zhu 2014; Arrivault et al. 2017) resembling those in C_3_ photosynthesis (Stitt et al. 1980; Arrivault et al. 2009). CBC flux *per se* in C_4_ plants is therefore likely to respond rapidly to a change in irradiance, as in C_3_ plants (see above). In contrast, metabolite pools in the CCM are large with a relatively slow turnover time (∼10 s, Stitt and Zhu 2014; Arrivault et *al*. 2017). The same holds for metabolites in the energy shuttle in NADP-ME species (3PGA, triose-P). The steady state size of these pools is larger in high irradiance than in low irradiance (Usuda 1987; Leegood and von Caemmerer 1988, 1989; Doncaster et al. 1989; Ubierna et al. 2013) presumably to generate the larger concentration gradients required to drive faster intercellular diffusion in high irradiance. There are interesting implications for the response of C_4_ photosynthesis to a sudden change in irradiance. On the one hand, these large pools might buffer against a sudden decrease in irradiance. Due to the very short half-lives of ATP and NADPH, in C_3_ plants there is an almost immediate inhibition of CO_2_ fixation after a decrease in irradiance (see above). As pointed out by Stitt and Zhu (2014, see also Slattery et al. 2018), the large pools of triose-P and malate in C_4_ photosynthesis might provide a buffer, with their conversion to 3PGA and oxaloacetate allowing continued generation of NADPH and ATP to support CO_2_ fixation for several seconds after a sudden decrease in irradiance. In C_3_ species, assimilation stops abruptly after darkening and decreases abruptly after a decrease in irradiance even when low O_2_ is used to suppress the post-illumination photorespiratory burst of CO_2_ (Lee et al. 2022; Arce-Cubas et al. 2023b). This contrasts with C_4_ species, where the decline in C assimilation is much slower (Laisk and Edwards 1998; Lee et al. 2022, Arce-Cubas et al. 2023b). On the other hand, slow build-up of these pools might delay the response of CO_2_ assimilation to an increase in irradiance. In agreement, pool sizes change slowly during transients from darkness to light (Leegood and Furbank1984; Usuda 1985) or high to low irradiance (Doncaster et al. 1989).

Slow changes in metabolite pool size may interact with or be superimposed on post-translational activation of CCM cycle enzymes. After a dark-light transition, activation of PEPC requires 10-15 min (Tirumala Devi and Raghavendra 1993), activation of pyruvate, phosphate dikinase (PPDK) 2-6 min (Roeske and Cholet 1989; Om et al. 2022) and NADP-MDH up to 10 min (Ashton et al. 1990) and modifies their kinetic properties and response to changes in metabolite levels (Ashton et al. 1990; Doncaster and Leegood 1987; Vidal et al. 2002). These enzymes are also post-translationally regulated in response to light intensity (Chen et al. 2014).

The multiplicity of decarboxylation pathways (see above) introduces a further complication. Although C_4_ species are classified into subtypes depending on the main route of decarboxylation, pathways often operate concomitantly (Furbank 2011; Bräutigam et al. 2014; Wang et al. 2014a). Recently, ^13^CO_2_ labelling studies with the NADP-ME species maize estimated that around 30% and 14% of the flux of 4C metabolites from the MC to the BSC is carried by aspartate rather than malate (Weissmann et al. 2016; Arrivault et al. 2017), with the implication that a similar proportion of the decarboxylation occurs by a route other than NADP-ME. It has been suggested that simultaneous operation of decarboxylases may provide added flexibility (Furbank 2011; Wang et al. 2014a; Bellasio and Griffiths 2014a). Analyses of pool sizes (Usuda 1987; Leegood and von Caemmerer 1989), enzyme activities (Sales et al. 2018) and ^13^CO_2_ labelling kinetics (Medeiros et al. 2022) in steady state conditions indicate that the contribution of NADP-ME decreases and that of other decarboxylation routes increases under low irradiance. It has also been reported that during a dark-light transition, aspartate and alanine levels decline in the first 5 min after illumination (Furbank and Leegood 1984) or that aspartate transiently declines and recovers and alanine transiently rises and falls (Usuda 1985). These observations indicate that the contribution of aspartate-dependent decarboxylation may change during induction of photosynthesis and, potentially, after sudden changes in light intensity.

There have been relatively few experimental studies of C_4_ photosynthesis during light transitions, and even fewer under fluctuating light. These are differing scenarios, especially when light is fluctuating rapidly (Arce-Cubas et al., 2023b), as is typically the case in a canopy. Studies with two C_3_ and two C_4_ species. Kubásek et al. (2013) revealed a similar or larger decrease in photosynthesis efficiency in C_4_ than C_3_ species in a regime with short periods of high light interspersed by ten minutes of low light. A similar conclusion was reached in comparisons of larger panels of C_4_ and C_3_ species under fluctuating light with shorter phases of low light (Li et al. 2021; Lee et al. 2023). However, except for paired Flaveria species in Li et al. (2021), these studies used phylogenetically unrelated species.

In agreement with the ideas of Stitt and Zhu (2014) in fluctuating regimes after switching to low light *A_n_* declined more slowly in C_4_ than C_3_ plants (Li et al. 2021; Lee et al, 2023; Arce-Cubas et al. 2023b). As this should even increase photosynthetic efficiency, any loss of efficiency in fluctuating regimes is presumably due to *A_n_* increasing slowly after switching to high light. *A_n_* was reported to rise more slowly in C_4_ species than C_3_ species after a sudden increase in light intensity (Kubásek et al. 2013) and under fluctuating light (Li et al. 2021; Lee et al. 2023). However, when three phylogenetically-related species pairs were compared in fluctuating light (Arce-Cubas et al. 2023b), *A_n_*increased at similar rates in C_4_ species and paired C_3_ species. This contrasted with a sudden dark-light transition (Arce-Cubas et al. 2023a) where *A_n_* increased more slowly in C_4_ species than paired C_3_ species. Li et al. (2021) suggested that the slow increase of *A_n_* in C_4_ plants after a low-high light transition might reflect the time needed to build-up metabolite pools. On the other hand, a theoretical study that extended a steady state model of NADP-ME type photosynthesis C_4_ photosynthesis (Wang et al. 2014b) to address dynamic changes predicted that the major restrictions were not slow build-up of metabolite pools but, instead, slow activation of Rubisco and PPDK and slow stomatal opening (Wang et al. (2021).

Thus, it remains an open question what underlying mechanisms impact C_4_ photosynthesis during changes in irradiance, and whether the impact is stronger than for C_3_ photosynthesis. e report experiments in which we subjected maize plants to a sudden decrease or increase in irradiance, monitored the rate of CO_2_ fixation and harvested material for detailed time-series metabolite profiling. The applicability of observations from single transitions to responses in fluctuating light may depend on the frequency of fluctuations, with the response of *A_n_* becoming increasingly similar when the fluctuations are slow (Arce-Cubas et al. 2023b). The overall goals were to provide a systems-level overview of the temporal kinetics of the response of C_4_ metabolism to a sudden decrease or increase in irradiance, and to compare these with responses under fluctuating light. More specifically, we wanted to interrogate the detailed metabolite time-course to (i) test if large pools of shuttle metabolites transiently buffer photosynthesis against a sudden drop in irradiance, (ii) test if build-up of shuttle metabolite pools delays the response to a sudden increase in irradiance, (iii) ask if the contribution of different decarboxylation routes changes during such transitions, (iv) ask if and how a sudden increase or decrease of irradiance perturbs the balance between the CCM and the CBC.

## Results

### Response of CO_2_ assimilation and stomatal conductance

We investigated the response to a relatively small change of irradiance in the limiting range, in order to focus on changes in C metabolism and minimize superimposed responses linked with activation and relaxation of energy dissipation mechanisms that arise in shifts between over-saturating and very low irradiance (see Stitt et al. 1989). The plants were grown at a moderate irradiance of 550 µmol photons m^-2^ s^-1^. In stable irradiation, net CO_2_ assimilation (*A_n_*) rose steeply up to about 800 µmol photons m^-2^ s^-1^ and more gradually as irradiance was increased further (Supplementary Fig. S1A).

We investigated transitions between moderate irradiance (550 µmol photons m^-2^ s^-1^, ML) and low irradiance (160 µmol photons m^-2^ s^-1^, LL). Steady state *A_n_* at ML and LL was about 25 and 10 µmol CO_2_ m^-2^ s^-1^, respectively (about 135 and 53 nmol CO_2_ g^-1^ FW s^-1^, estimated using a specific leaf area of 0.0055 m^-2^ g^-1^(Medeiros et al. 2022), corresponding to about 74 and 29%, respectively, of the maximum rate of photosynthesis (Supplementary Fig. S1A), Plants were illuminated at growth irradiance (= ML) for at least four hours and then transferred to 160 µmol photons m^-2^ s^-1^ (ML-LL) or were illuminated from the start of the light period at 160 µmol photons m^-2^ s^-1^ for at least four hours before transfer to 550 µmol photons m^-2^ s^-1^ (LL-ML). Gas exchange parameters were logged every second, starting 15 min before and continuing until 30 min after the light transition. As gas exchange measurements shortly after the light transitions strongly violate the steady state assumption underlying default rate equations, dynamic equations were implemented (Saathoff and Welles 2021). Figs. 1A-B show the temporal response of *A_n_* during transitions with time on a log scale (see Supplementary Figs. S1B and S1F for plots with a linear time scale and Supplementary Dataset S1 for the underlying data).

**Figure 1.**
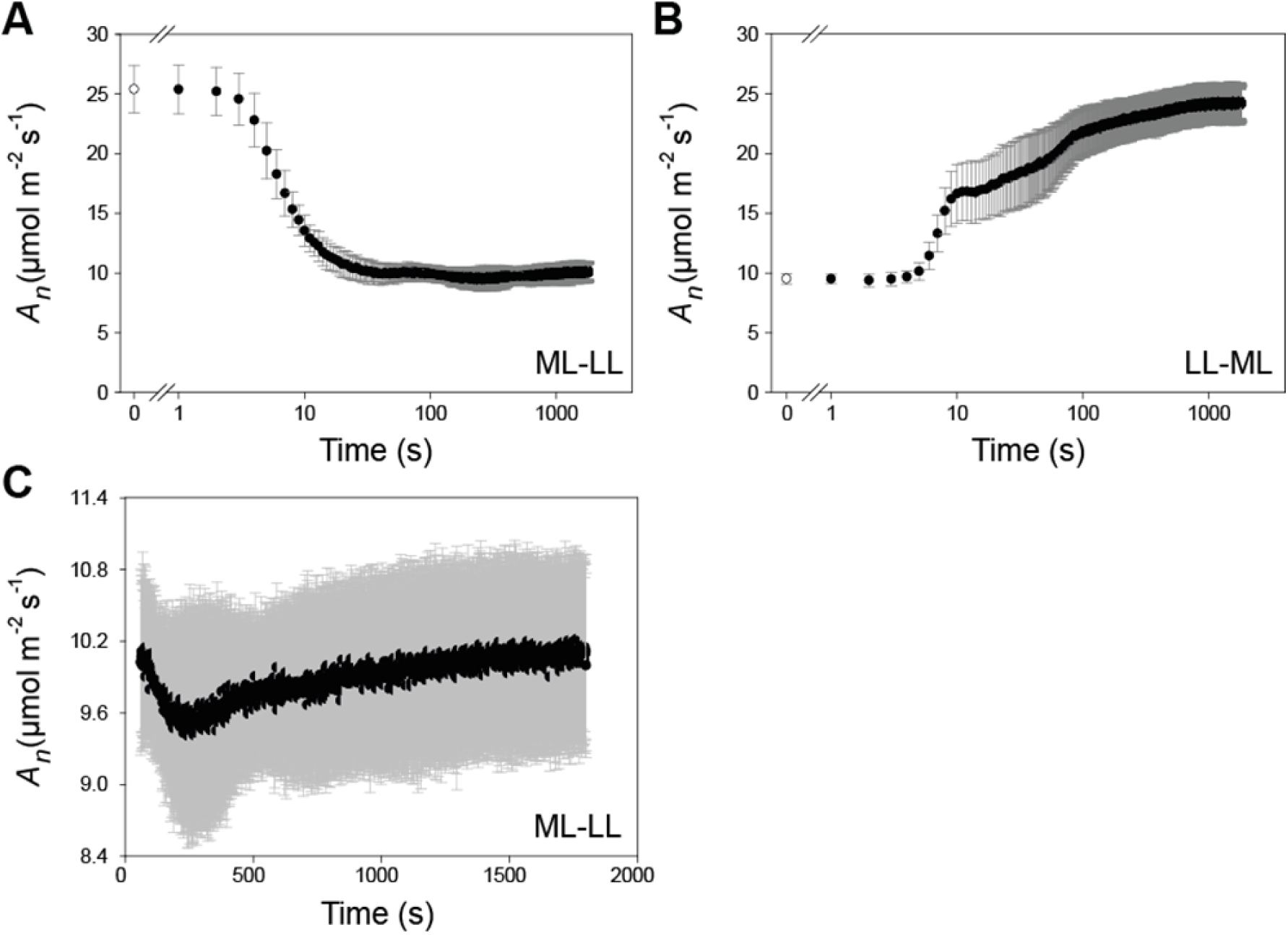
CO_2_ assimilation. (**A**) Responses during a moderate light-low light (ML-LL) transition and (**B**) a low light-moderate light (LL-ML) transition. X-axes correspond to time on a log scale. As gas exchange measurements shortly after the light transitions violate the steady state assumption underlying default rate equations, dynamic equations were implemented (see Methods). Corrected values are shown for each second. Data shown in (**A**) and (**B**) are mean ± SD (*n*=5 for ML-LL, *n*=6 for LL-ML). The time points in white correspond to time 0, before the light transition. The gap in the x-axes is for a better visualization of the first two time points. (**C**) Expansion of the ML-LL response from 60 s onwards to show the slight recovery of *A_n_*. The display shows mean ± SD (*n*=5). X-axis corresponds to time on a linear scale. The relatively high SD is due to differences in *A_n_* between the five independent replicates (see Supplemental Dataset S1). Significance was analyzed using a paired t-test to compare *A_n_* at different times in a given replicate, and separate this from differences in *A_n_*between replicates. The increase between the trough and the end of the transition was significant (p = 0.002, *n*=5, paired t-test comparing, for each replicate, the average *A_n_* between 250-259 s with average *A_n_* between 1791-1800 s). See Supplementary Figs S1B and S1BF for plots of (**A**) and (**B**) on a linear time scale, Supplementary Fig. S1C and Supplementary Fig. S1G for curve-fitting to different time spans of the ML-LL and LL-ML responses and Supplementary Fig. S1D-E and H-I for stomatal conductance and *Ci.* The original data are provided in Supplementary Dataset S1A.

In the ML-LL transition (Fig. 1A), *A_n_* remained high for 3 s, started to fall at 4 s, was significantly lower at 5 s (*p*= 0.006, Student’s t-Test), continued to decline until ∼250 s, and then recovered slightly until the end of the measurements at 1800 s (Fig. 1C). The initial decrease was quasi-exponential with a t_0.5_ of about 11 s (Supplementary Fig. S1C). After 5, 10 and 15 s in LL, *A_n_* was 80, 53 and 45%, respectively, of that in ML, compared to a value of 40% at the end of the transitionAt the trough, *A_n_* was 38% of that in ML and 94% of the steady state rate in LL. The subsequent recovery was significant (*p* = 0.002, paired t-test, legend of Fig. 1C), represented a 6% increase in *A_n_* and was equivalent to 7.6% of Δ*A_n_* (the difference between *A_n_* in ML and *A_n_* after 1800 s in LL). Stomatal conductance (*g*) responded more slowly, with the decline starting after about 30 s and continuing until about 720 s, when *g* was about 50% of that in ML (Supplementary Fig. S1D). Internal CO_2_ concentration (*Ci*) rose by 190% to a peak at about 30 s, before declining until about 12 min, when *Ci* was about 35% higher than in ML (Supplementary Fig. S1E).

In the LL-ML treatment (Fig. 1B; Supplementary Fig. S1F), *A_n_*showed no significant change in the first 5 s and then rose, starting at 6 s (20% above that in LL, p= 0.003, Student’s t-Test). The rise can be divided into four phases, which were curve fitted (Supplementary Fig. S1G). An initial fast phase slowed down by 9-10s and accounted for 49% of the overall rise in the rate of photosynthesis. *A_n_* then plateaued until about 15 s and rose again, first until 90 s (31% of the rise) then slowly until the end of measurements (20% of the rise). *g* increased gradually between 30 s and 6 min (Supplementary Fig. S1H). *Ci* declined about 3-fold to a minimum at 50 s before rising about 1.7-fold in the next 4 min (Supplementary Fig. S1I).

### Global analysis of response of metabolism

The underlying changes in metabolism were investigated in a separately-grown batch of plants. Gas exchange was also measured in these plants. For technical reasons, the correction for time lag could not be performed, but the essential features of response of *A_n_*were reproduced (see Supplementary Text Section 1 and Supplementary Fig. S1). Replicate samples were collected before (0 s) and 5, 10, 15, 30, 60, 120, 300, 600, 1200 and 1800 s after transfer to the new irradiance (*n*=4, except for time 0 of LL-ML, *n*=10). Samples were collected in a randomised manner between 4-6 hour into the light period. The ML-LL and LL-ML experiments were performed one week apart on separately-grown batches of plants. Analysis by LC-MS/MS, GC-MS and enzymatic assays allowed detection and quantification of 36 metabolites (Supplementary Dataset S2). To provide a global overview of the response of metabolism, we first performed principal component (PC) analysis with all metabolites and *A_n_* (Fig. 2, Supplementary Fig. S2).

**Figure 2.**
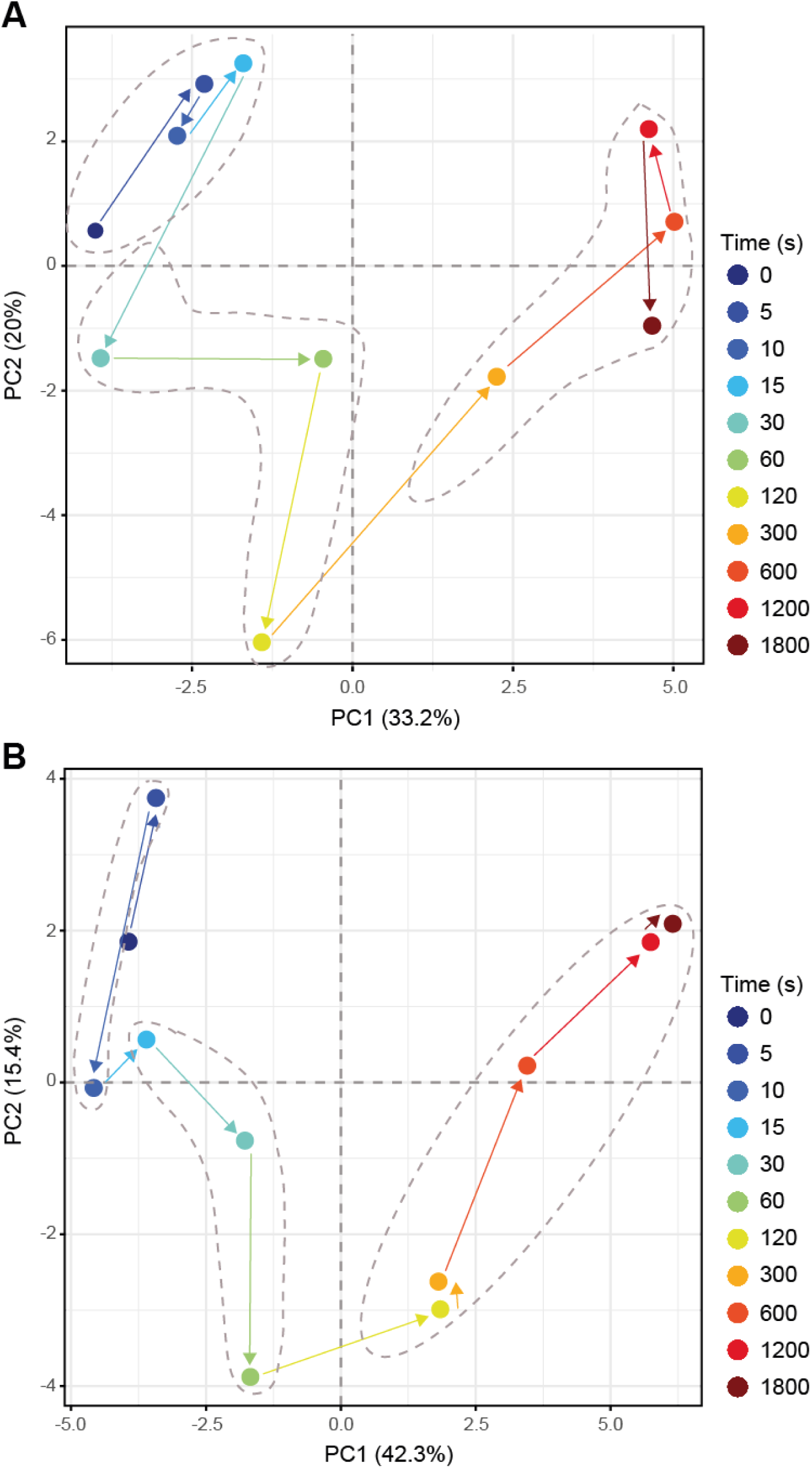
Principal Components Analysis. (**A**) Analysis for a moderate light-low light (ML-LL) transition and (**B**) analysis for a low light-moderate light (LL-ML) transition. Means for each time point were used to perform the analyses (*n*=4 to 5 for ML-LL and *n*=4, except for time 0 s where *n*=10 for LL-ML). The time points are identified by color (see insert) and the arrows show the time sequence in the transition. Different phases are indicated by dotted grey ellipses. Data are provided in Supplementary Dataset S2. Plots showing loadings of metabolites in PC1 and PC2 are shown in Supplementary Fig. S2.

In the ML-LL transition (Fig. 2A), PC1 and PC2 accounted for 33 and 20% of total variation, respectively. Visual inspection pointed to three sequential responses; an initial (0 to 15 s) shift to a more positive value in PC2 (time points 5, 10 and 15 s are close to each other), followed (15-120 s) by a large shift to negative values in PC2, followed (120 s onwards) by a large shift to positive values in PC1 and reversal of the changes in PC2. The first phase corresponds to the initial somewhat delayed decrease of *A_n_*. the second phase mainly to the continuing slow decline of *A_n_*that was completed by 250 s and the third phase to the subsequent small recovery of *A_n_* (see Fig. 1C; Supplementary Fig. S1L). PC1 differentiated the steady state metabolite profiles in ML and LL and captured gradual changes during adjustment to LL, whilst PC2 captured transient changes. Inspection of loadings (Supplementary Fig. S2A) reveals that PC1 was driven by intermediates in the CBC (mainly ribulose 5-phosphate + xylulose 5-phosphate [Ru5P+Xu5P], ribose 5-phosphate [R5P], ribulose-1,5-bisphosphate [RuBP]), in starch synthesis (ADP-glucose [ADPG]), CCM (pyruvate, alanine) and in photorespiration (glycine, glycerate) as well as 2-oxoglutarate (2OG), all of which align with *A_n_*, and by contrasting responses of aspartate, glutamate and fumarate. There was no large contribution from fructose 1,6-bisphosphate (FBP) or sedoheptulose 1,7-bisphosphate (SBP). PC2 was mainly driven by 3PGA, sedoheptulose 7-phosphate (S7P), fructose 6-phosphate (F6P), Ru5P+Xu5P and ADPG, which align with *A_n_*, and reciprocal changes of FBP, SBP, RuBP, 2OG and alanine. It should be noted that the response of malate is not informative, because maize leaves contain a large pool of malate that is not involved in C_4_ photosynthesis (Stitt and Heldt 1985a; Weismann et al. 2016; Arrivault et al. 2017) and masks any changes in the malate pool that is directly involved in the CCM.

In the LL-ML shift (Fig. 2B) PC1 and PC2 accounted for 42 and 15% of total variation, respectively. Three sequential responses could be distinguished. The first (between 0 and 5 s (less well-defined as it depends on one time point only) was associated with a slight shift to more positive values in PC2, the second (5-60 s) by a large shift to negative values in PC2, and the third (60 s onwards) by a strong shift to positive values in PC1 and a reversal of the changes in PC2. The first phase (0-5 s) corresponds with the time where *A_n_*did not change. The second phase corresponds to the rapid rise in *A_n_*between 5 and 11 s, the plateau until 15 s (time points 10 and 15 s are close to each other) and the subsequent rise until about 90 s. The third phase corresponds to the subsequent slower rise in *A_n_* (see Fig. 1B and Supplementary Fig. S1G). PC1 differentiated steady state metabolite profiles in LL and ML and captured gradual changes to the adjustment to ML, whilst PC2 captured transient changes. Inspection of loadings (Supplementary Fig. S2B) revealed that PC1 is driven by changes of most CBC intermediates (3PGA, DHAP, SBP, S7P, R5P, Ru5P+Xu5P, RuBP), some CCM intermediates (pyruvate, alanine, PEP), glycine and 2OG, which all aligned with *A_n_*, and contrasting responses of aspartate and to a lesser extent FBP. PC2 was mainly driven by glucose 6-phosphate (G6P), sugars and serine, and contrasting changes of RuBP, ADPG, PEP, glycerate and, to a lesser extent, FBP and SBP.

### Time-dependent changes of individual metabolites, metabolite ratios and sets of metabolites

We next inspected the temporal responses of individual metabolites. Temporal responses in the detailed time course are shown for ML-LL in Fig. 3A and Supplementary Fig. S3A, and for LL-ML in Fig. 4 and Supplementary Fig. S4A. Supplementary Figs. S3B and S4B provide a summary of statistically significant changes. We tested for changes compared to time zero, and for changes within later time intervals. We did this because testing against zero might not capture important changes in later phases of the multi-phasic responses. The PC analysis in Fig. 2 was used to define the time intervals. We also investigated changes of metabolite ratios (Figs. 3B and 4B). These provide information about poising; the extent a particular reaction is restricting flux in an entire pathway. We estimated metabolite ratios for 3PGA/DHAP (indicative of the supply of NADPH and ATP, Dietz and Heber 1984), FBP/F6P, SBP/S7P and pentose-P (sum of R5P, Ru5P+Xu5P)/RuBP (indicative of resistance to flux at FBPase, SBPase, and phosphoribulokinase, [PRK], respectively), and RuBP/3PGA and RuBP/2PG (indicative of resistance to flux at the Rubisco carboxylase and oxygenase reactions, respectively). We estimated pyruvate/alanine ratios, which provide indirect evidence about the contribution of NADP-ME and other decarboxylation routes, and the 2OG/glutamate ratio which provides information about the availability of amino groups and organic acid acceptors. In addition, we summed the amount of C in sets of metabolites that are involved in the CBC and the CCM, as well as the amount of P in CBC intermediates (Figs. 3C and 4C). The CBC pools include 3PGA and DHAP, which are involved in the energy shuttle. Malate was excluded from the metabolites used to estimate the CCM pool because much of the malate in a maize leaf is in a pool not directly involved in the CCM (see above).

**Figure 3.**
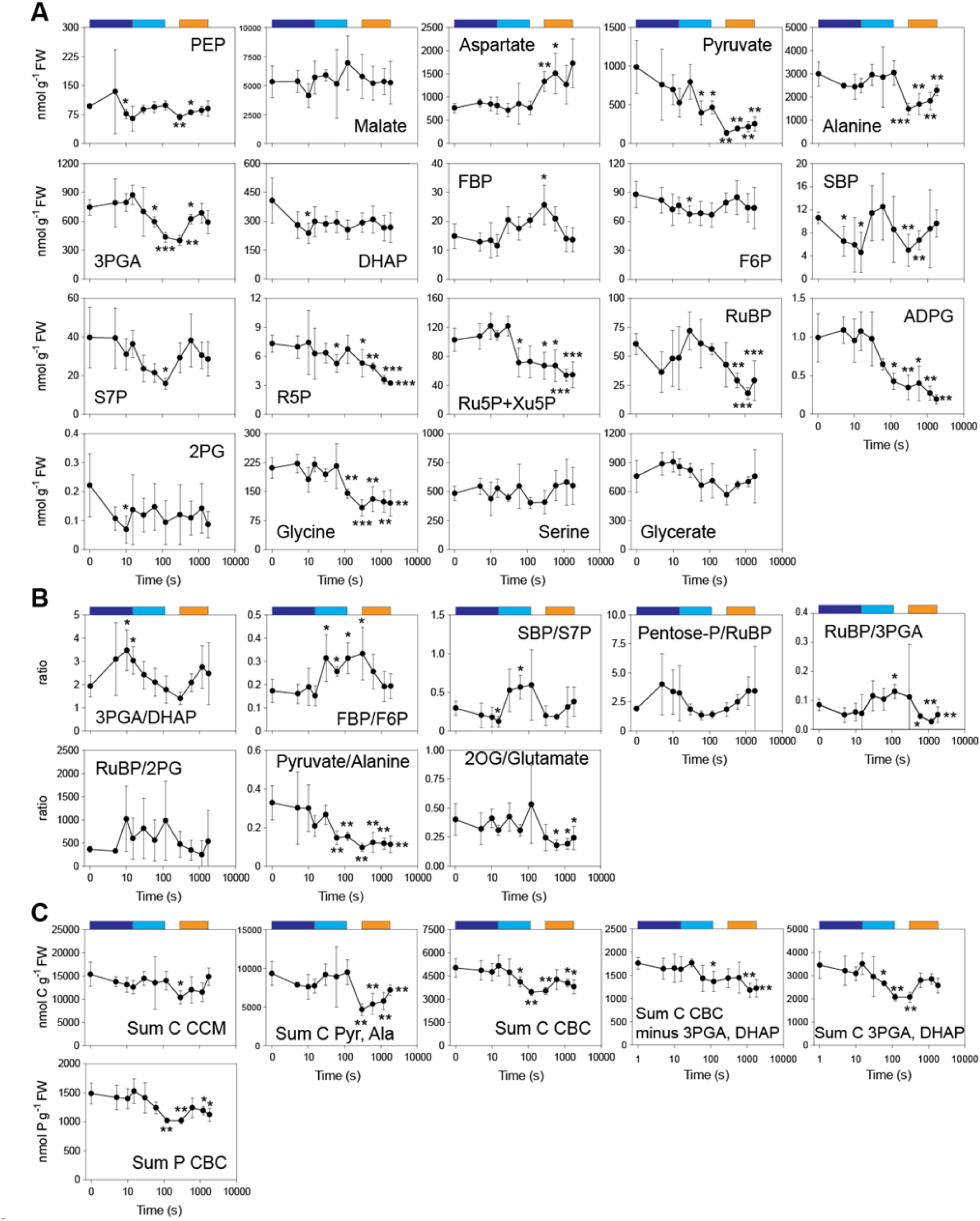
**Changes of individual metabolites, metabolite ratios and sets of metabolites in a moderate to low light transition**. (**A**) Amounts of metabolites (nmol g^-1^ FW) involved in CCM, CBC (including ADPG), and photorespiratory pathway, (**B**) ratios of metabolite amounts, and (**C**) sum of C in CCM and sums of C and P in CBC (nmol C g^-1^ FW and nmol P g^-1^ FW). In (**C**) metabolites summed for CCM were PEP, aspartate, pyruvate and alanine (malate was not included because the active pool could not be defined). Data is shown as mean ± SD (*n*=4 to 5). Significant differences (t-test) to time zero (i.e., ML) are indicated by stars (* p < 0.05; ** p < 0.01: *** p < 0.001). Time is shown on a log scale. The colored boxes above the graphs indicate the segments used to perform additional t-tests (Supplementary Fig. S3B). The original data are provided in Supplementary Dataset S2. Further metabolite levels are shown in Supplementary Fig. S3A.

**Figure 4.**
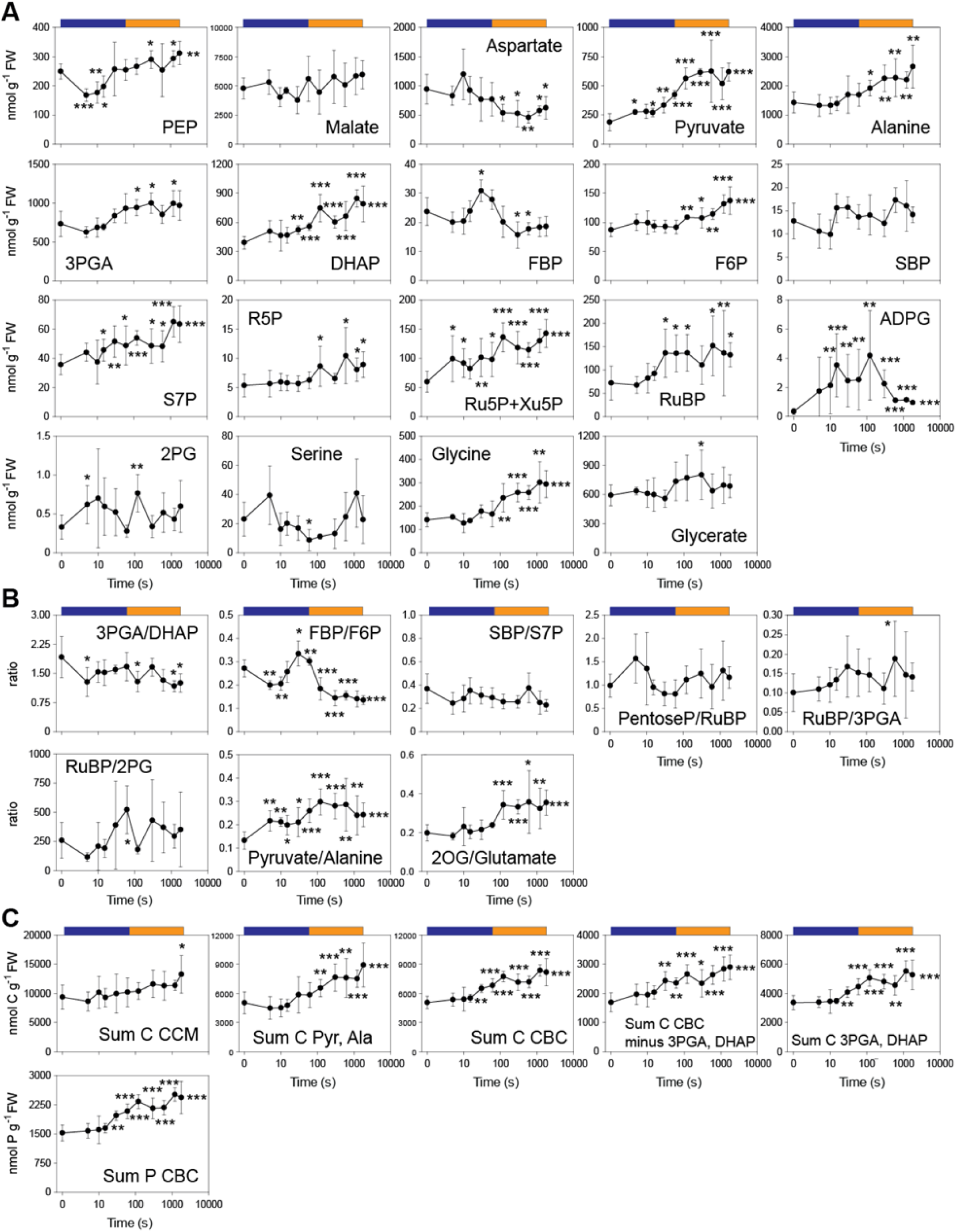
**Changes of the levels of individual metabolite ratios and sets of metabolites in a low to moderate light transition**. (**A**) Amounts of metabolites (nmol g^-1^ FW) involved in CCM, CBC (including ADPG), and photorespiratory pathway, (**B**) ratios of metabolite amounts, and (**C**) sum of C in CCM and sums of C and P in CBC (nmol C g^-1^ FW and nmol P g^-1^ FW). In (**C**) metabolites summed for CCM were PEP, aspartate, pyruvate and alanine (malate was not included because the active pool could not be defined). Data shown are means ± SD (*n*=4, except for time 0 s where *n*=10). Significant differences (t-test) to time zero (i.e., ML) are indicated by stars (* p < 0.05; ** p < 0.01: *** p < 0.001). Time is shown on a log scale. The colored boxes above the graphs indicate the segments used to perform additional t-tests (Supplementary Fig. S4B). The original data are provided in Supplementary Dataset S2. Further metabolite levels are shown in Supplementary Fig. S4A.

In the next two sections, the temporal response of metabolite levels, metabolite ratios and summed C in pathways are presented in the context of the sequential phases identified in gas exchange and the PC analyses. Metabolite ratios and summed C in pathways are collectively referred to as ‘metabolic traits’.

There is scatter in the levels of several metabolites in leaves of C_4_ species including maize (see also Leegood and von Caemmerer 1988, 1989). We therefore performed further experiments in which we sampled larger numbers of replicates at 0, an early (10 s) and a late /1200 s) timepoint after a ML-LL or a LL-ML transition (Supplementary Figs. S5A-B, Supplementary Dataset S3). Most metabolites showed qualitatively consistent responses to those in the detailed time course, but a small number differed (Supplementary Figs. S3C-D; in ML-LL, the RuBP/2PG ratio at 10s and the RuBP/3PGA and RuBP/2PG ratios at 20 min; in LL-ML, ADPG at 10 s, FBP and SBP at 10 and 1200 s, and fumarate, fructose and raffinose at 1200 s).

### Time-dependent responses in the ML-LL transition

The first phase of the ML-LL response as defined by PCA (Fig. 2) was from 5-15 s, in which time *A_n_*, after a 3 s delay declined at 5, 10 and 15 s to 80, 53 and 45%, respectively, of that in ML (Fig. 1A). This somewhat delayed decline of *A_n_* was associated with a slight increase in 3PGA, a large decrease (∼108 nmol g^-1^ FW) in DHAP (Fig. 3A) and a large increase of the 3PGA/DHAP ratio (Figs. 3A-B, significant for DHAP and the 3PGA/DHAP ratio (Supplementary Fig. 3SB).. There was a small significant decrease of SBP (Fig. 3A) and the SBP/S7P ratio (Figs. 3A-B) but not of FBP or the FBP/F6P ratio or of RuBP. Total C and P in CBC intermediates remained high (Fig. 3C). ADPG remained high (Fig. 3A) consistent with CBC operation continuing for a short time after switching to LL. There was a rapid significant drop in 2PG (Fig 3A; Supplementary Fig. 3SB), the RuBP/2PG ratio remained unaltered or rose and the RuBP/3PGA ratio showed a non-significant decrease (Fig. 3B), consistent with slower oxygenation relative to carboxylation of RuBP.

The next phase of the ML-LL transition (until ∼120 s) corresponded to a further slow decline in *A_n_*. During this time there was a significant decrease of 3PGA (Fig. 3A; Supplementary Fig. 3SB) and the 3PGA/DHAP ratio (Fig. 3B; Supplementary Fig. S3B, see analysis for the 15-120 s time segment), significant increases of the FBP/F6P and SBP/S7P ratios (Fig. 3B; Supplementary Fig. S3B), a significant decline of S7P and pentose-P, a non-significant decline of the pentose-P/RuBP ratio, a significant decline in ADPG (Fig. 3A; Supplementary Fig. S3B) and a significant decline in total C and P in CBC intermediates (Fig. 3C; Supplementary Fig. 3SB) RuBP levels showed a non-significant decline and recovery to a similar level to that in ML, whilst the RuBP/3PGA ratio rose (significant at 120 s Supplementary Fig. S3B. Pyruvate started to decline, but alanine, and the summed CCM metabolite pool (PEP, aspartate, pyruvate and alanine) remained high (Figs. 3A, 3C). In photorespiration, 2PG remained low, the RuBP/2PG ratio high and glycine started to decline (Fig 3A; Supplementary Fig. S3B. Overall, these observations indicate that CBC flux is falling due to a shortfall of energy, that C is draining from the CBC, that FBPase, SBPase and Rubisco are being inactivated, and that CCM pools initially remain relatively high.

The third phase corresponded to the small but highly significant recovery of *A_n_* from ∼250 s onwards. During this time there was an increase of 3PGA and the 3PGA/DHAP ratio (Figs. 3A-B, significant when tested in this phase Supplementary Fig. S3B), a decrease of the FBP/F6P (significant when tested in this phase), a significant decrease of pentose-P, RuBP and the RuBP/3PGA ratio (Fig. 3B; Supplementary Fig. 3SB), whilst total C and P in CBC intermediates rose slightly (Fig. 3C, significant when tested in this time segment, Supplementary Fig. S3B). Pyruvate and alanine decreased between 120 and 300 s, and then rose. Aspartate, which did not change in the first part of the transient, increased markedly from 300 s onwards (Fig. 3A, significant when tested in this phase, Supplementary Fig. S3B). Fumarate also increased (Supplementary Fig. S3A, significant from 300 s onwards). As malate and fumarate are interconverted in a reversible reaction catalysed by fumarase, this increase of fumarate provides indirect evidence for an increase of a metabolically-active malate pool. There was a marked and significant decrease in the 2OG/glutamate ratio (Fig. 3B; Supplementary Fig. S3B). This occurred even though aspartate was increasing and the summed level of serine and glycine did not change. The most obvious source of amino groups was the large decrease in alanine between 120 and 300 s. Overall, these changes point to adjustments in poising of the CBC that allow more effective use of light energy and a slight increase in *A_n_*. This is linked possibly with activation of plastidic FBPase, SBPase and changes favoring RuBP utilization. They also point to adjustment in poising of the CCM, including a rise in the pools of aspartate, possibly malate, pyruvate and alanine.

### Time dependent response of individual metabolite traits in the LL-ML transition

After an increase in irradiance, rapid metabolic adjustments occurred in the first 5 s, before there was a significant change in *A_n_*(Fig. 1B). This included a trend to lower 3PGA, a significant rise in DHAP and a significant decrease in the 3PGA/DHAP ratio (Figs. 4A-B; Supplementary Fig. S4B), presumably because the increase in irradiance leads almost immediately to increased availability of ATP and NADPH for 3PGA reduction. It might be noted the light reactions immediately generate more ATP in both the MC and BSC, but NADPH mainly in the MC. There was a significant decrease in the FBP/F6P ratio (Fig. 4B) due to small non-significant reciprocal changes of FBP and F6P (Fig 4A). The reciprocal changes of DHAP and FBP immediately after an increase in irradiance may point to flux restriction at aldolase, or to complexities introduced by intercellular compartmentation (see Discussion). In the CCM there was a significant ∼40% decrease in PEP and a significant increase in pyruvate (Fig. 4A: Supplementary Fig. S4B, pointing to increased PEPC activity, faster decarboxylation by NADP-ME and temporary restriction of flux at PPDK.

The second phase, lasting until 90 s, was associated with a marked increase in *A_n_* that accounted for about 80% of the overall gain over the entire transition from LL to ML (see Fig. 1B). This phase consisted of three parts; a rapid increase of *A_n_* that started at 5 s and slowed down by 9-10 s (49% of the total rise), a plateau until about 15 s, and a rapid increase of *A_n_* (31% of the rise) until 90 s, albeit not as rapid as between 6 and 9 s (Fig. 1B).

The rise of *A_n_* after 5 s, slowdown by 9 s and plateau from 11-16 s were not accompanied by marked changes in metabolite levels. There was a significant increase of ADPG at 10 s (but this was not seen in the second experiment). By 15 s the FBP/F6P ratio started to rise indicating a restriction on flux at plastidic FBPase. The initial decline of the 3PGA/DHAP ratio started to revert, reflecting increased use of light energy as *A_n_* rose. There was a marked and significant decline of PEP at 5-15 s, reflecting either decreased synthesis from 3PGA and/or insufficient flux at PPDK relative to it consumption by PEPC points to a temporary restriction of PEPC activity by PEP (see Supplementary Text Section 2.2 for more details.

Between 15 and 90 s, *A_n_* rose at an accelerating pace (Fig.1B). Time point 90 s was not collected for metabolite analyses but by 60 s, when the rise in *A_n_* represented about 70% of the gain in the entire LL to ML transition, there was a coordinated increase in the levels of many CBC metabolites including 3PGA, DHAP, FBP, F6P, S7P, pentose-P, and RuBP (Fig. 4A, all significant, see Supplementary Fig. S4B). The FBP/F6P ratio, which rose at 30-60 s indicating a restriction on flux at FBPase, subsequently decreased (Fig. 4B; Supplementary Fig. S4B). There was a significant ∼2-fold increase in the sum of C and of P in CBC intermediates (Fig. 4C; Supplementary Fig. S4B). Together, these observations point to the increase in CBC flux being linked both to activation of CBC enzymes and to a general increase in the pool sizes of CBC intermediates.

There was a highly significant 10-fold increase in ADPG (Fig. 4A; Supplementary Fig. S4B). The increase at 10 s was not seen in the replicate experiment (Supplementary Fig. S5D) but the considerable noise in these early time points might reflect variation in the speed of the response (see Supplementary Text Section 2.2). The increase in ADPG was accompanied by a decrease in G6P (Supplementary Fig. S5, significant at 30 and 60 s; Supplementary Fig. S4B). In C_3_ species such as spinach and barley (Stitt et al. 1980, 1983; Gerhardt et al. 1987), changes in total G6P reflect the behavior of the cytosolic G6P pool because the majority of the G6P is located in the cytosol, whilst F6P is more evenly distributed between chloroplast and cytosol. Assuming a similar distribution in maize, the decline in total G6P is consistent with a decline of G6P in the cytosol in the MC where sucrose synthesis occurs (Furbank et al. 1985). This is consistent with a delay in activation of flux at cytosolic FBPase. This might also explain the strong rise in ADPG. Any delay in activating end-product synthesis will support build-up of the CBC and CCM metabolite pools (see Discussion).

In the CCM, as already mentioned, PEP declined by 5 s and recovered from 15 s onwards (Fig. 4A, highly significant, see Supplementary Fig. S4B. The response of PEP qualitatively resembled but was larger than that of 3PGA. There was a significant increase in pyruvate from 15 s onwards, and an upward trend for alanine (but only significant from 120 s onwards) (Fig. 4A; Supplementary Fig. S4B. This points to increased movement of C from the CBC via PEP into the CCM intermediate pools. The 2OG:glutamate ratio started to rise, mainly due to an increase in 2OG (Fig. 4B; Supplementary Fig, S4, the changes until 60 s were non-significant but the trend is confirmed by significant changes at 120s).

From 90 s onwards, *A_n_* continued to rise although more slowly, with this phase contributing ∼20-35% of the total increase in *A_n_*. Within the CBC, there was a further rise of DHAP, F6P, S7P but not of 3PGA or RuBP, whilst FBP declined (Fig. 4A; Supplementary Fig. S4B). Together, this resulted in a further small increase of total C or P in CBC intermediates (Fig. 4C; Supplementary Fig. S4B). There was a progressive and significant rise in G6P and UDPG (Supplementary Fig. S5, significant for both when tested in this time segment, Supplementary Fig. S4B), which is indicative of increased flux over the cytosolic FBPase to sucrose (see above). There was a 5-fold decrease in ADPG, indicative of slower starch synthesis (Fig. 4A). The most marked changes after 120 s were further significant increases in PEP, pyruvate and alanine (Fig. 4A) and summed CCM metabolites (Fig. 4C; Supplementary Fig. S4B. *Ci* fell to a minimum (∼80 ppm) after 60-100 s before slowly rising to about 140 ppm (Fig. 1B; Supplementary Fig. S1B). There was no consistent change in 2PG levels or the RuBP/2PG ratio, but there was a progressive rise in glycine (Fig. 4A; 4B).

### Correlation analysis of A_n_ with metabolite levels and metabolic traits

The above analysis pointed to the metabolic response in the ML-LL and LL-ML transitions involving slow and progressive changes in the levels of some metabolites, and transient changes of other metabolites. We determined Pearsońs correlation coefficients and hierarchical clustering across each transition to search for sets of metabolites that change in a coordinated way and, in particular, to detect metabolites that correlate with *A_n_*. In this analysis we excluded the time zero (i.e., the value before changing the irradiance) in order to focus on the response as metabolism adjusts to the new irradiance. Fig. 5 provides an overview of the metabolic connectivity (see Supplementary Figs. S6A-B for expanded plots). Further correlation analyses were performed between *A_n_* (again excluding the time zero point) and metabolic traits (Fig. 6A). Plots of the relationship between *A_n_* and selected metabolites or metabolic traits are shown in Figs. 5B-C and 6B-C. and plots for all metabolites and metabolic traits are provided in Supplementary Figs. 7B-C and 8B-C. We also compared selected metabolic traits with each other, again excluding the time zero point (Fig. 7).

**Figure 5.**
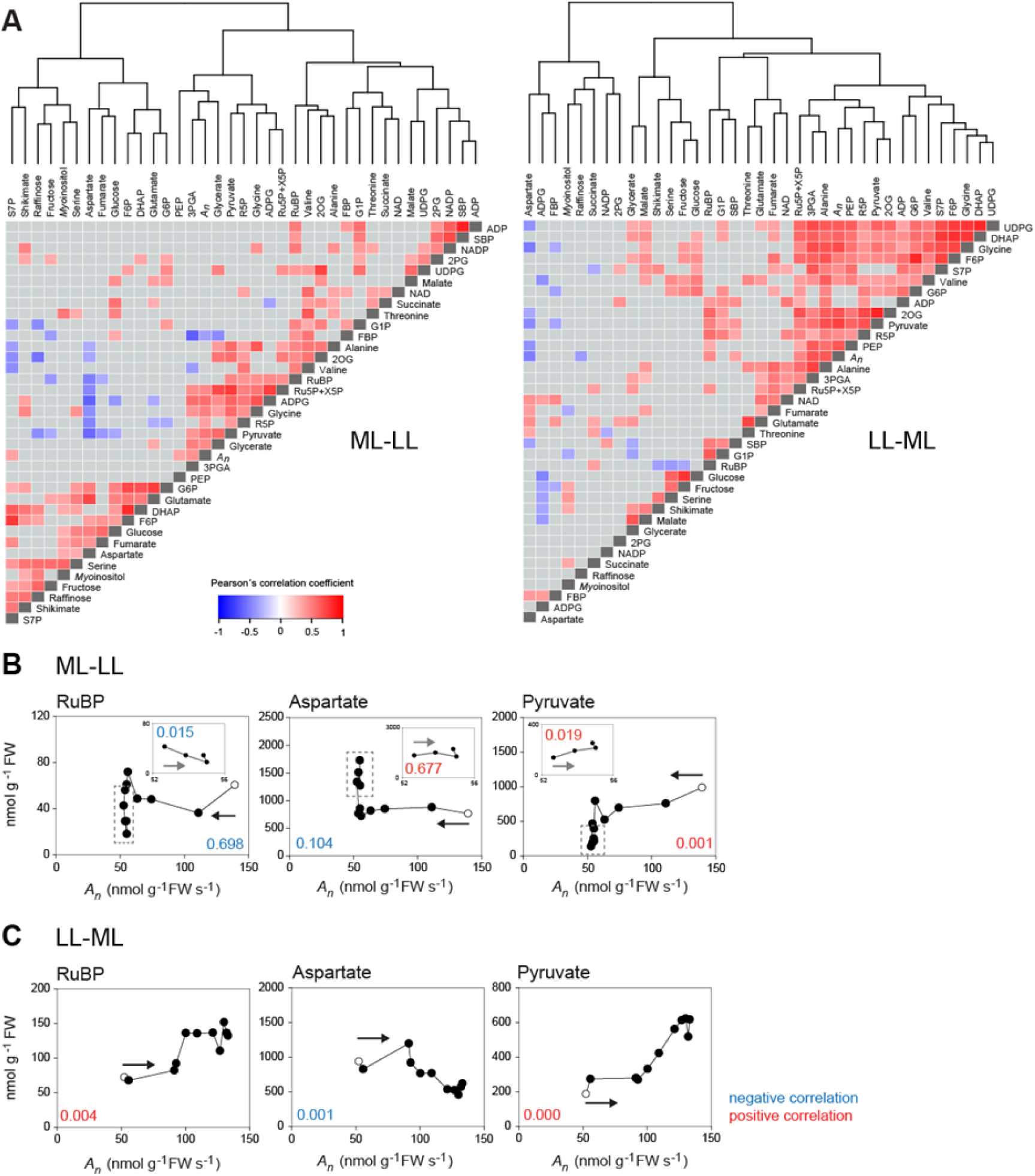
**Correlations between metabolites and *A_n_.*** (**A**) Heatmaps showing Pearsońs correlation coefficients for metabolites and CO_2_ assimilation (*A_n_*) in the ML-LL transition (left hand side) and LL-ML transition (right hand side). Pearson’s correlation coefficients were calculated using individual samples at all time points after the transition to LL or the transition to ML (time zero excluded). *A_n_*was taken from Supplementary Dataset S1. Significant correlations (*p* < 0.05) are colored red (positive) and blue (negative) (see insert), while correlations that were not significant are shown in grey. Self-correlations are identified in dark grey. The hierarchical clustering is shown. Larger displays to aid identification of individual metabolites are provided in Supplementary Fig. S6A-B and a summary of R^2^ and *p*-values in Supplementary Fig. S6C. (**B**) Response of selected metabolites (y-axis) in the ML-LL transition versus *A_n_*(x-axis). (**C**) Response of selected metabolites (y-axis) in the LL-ML transition versus *A_n_* (x-axis). For clearer visualisation, the means are shown (*n*= 3 to 10). In panel **B**, the open symbol is the value at ML, and the filled symbols are values after transition to LL, with the time sequence running from left to right (as indicated by a black arrow). The insert in each display shows time points between 300-1800 s (dotted grey box in main panel), with an expanded scale for *A_n_*(52 – 56 µmol CO_2_ g^-1^ FW s^-1^) to visualize correlations during the slight recovery of *A_n_* after 250 s (see Supplementary Fig. S1C). In the insert the time sequence runs from right to left (indicated by a red arrow). In panel **C**, the open symbol is the value in LL, and the filled symbols are values after different times in ML, the time sequence runs from right to left (rising rate of photosynthesis with time). In both **A** and **B**, R^2^, slope direction and *p* values were calculated by linear regressions using individual samples at all time points after the transition (time zero excluded) and in the case of panel **A**, also from 300 to 1800 s (in insert). The *p* value is colored according to the slope direction; red and blue for positive and negative correlations, respectively). Additional responses of metabolites versus *A_n_* are provided in Supplementary Figs S7A and S8A. The original data are provided in Supplementary Dataset S2.

**Figure 6.**
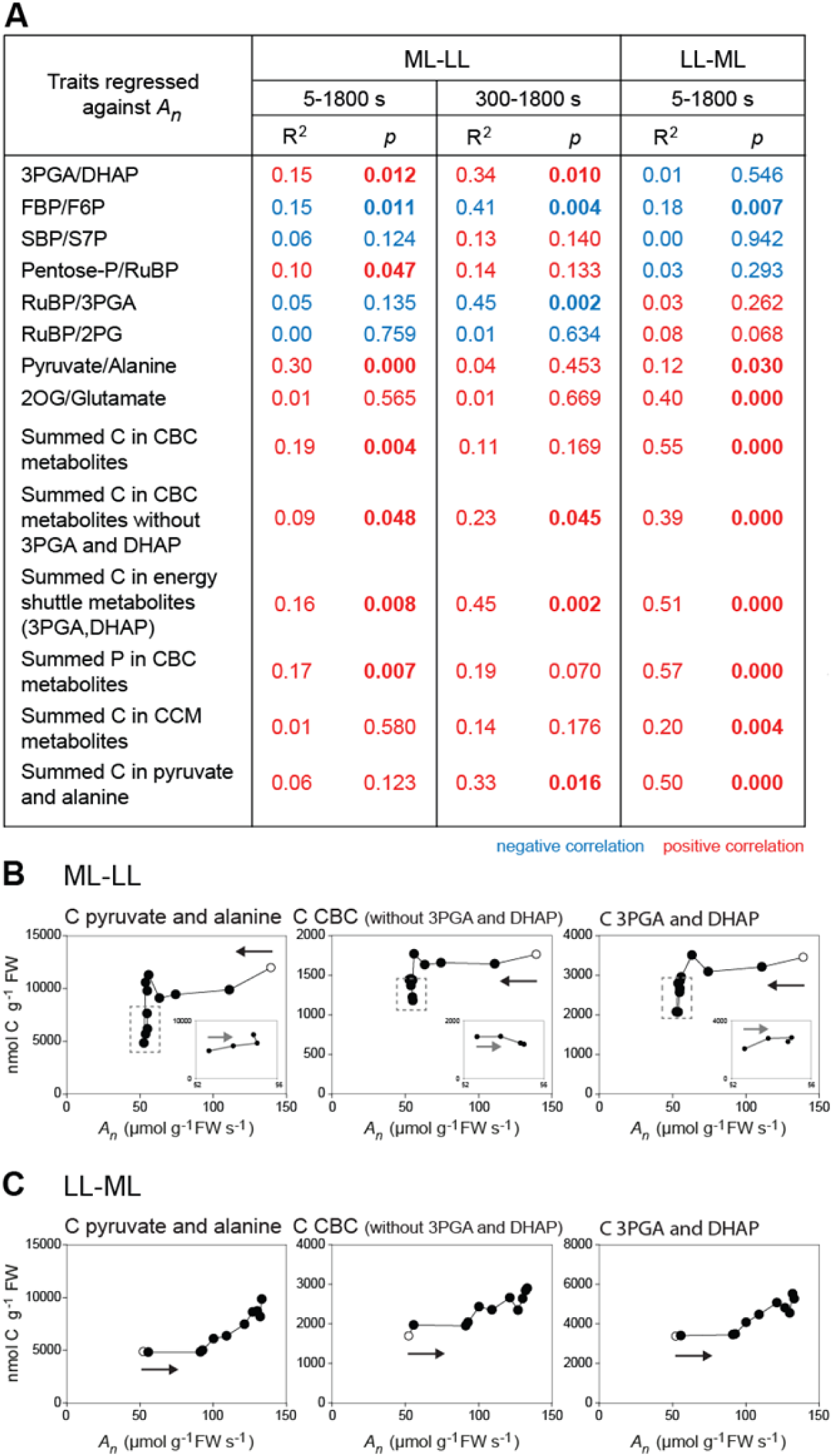
Correlations between metabolic traits and *A_n_*. (**A**) Summary of R^2^, slope directions (slope d.) and *p* values. These were calculated by linear regressions using individual samples (*n*=3 to 5) at all points after the transition (time zero excluded). Additional values were calculated from 300 to 1800 s for the ML-LL transition. Positive and negative correlations are indicated by color (red and blue, respectively). Those with significant *p* values (*p* < 0.05) are in bold. (**B**) Response of summed C pools (y-axis) in the ML-LL transition versus *A_n_* (x-axis). (**C**) Response of summed C pools (y-axis) in the LL-ML transition versus *A_n_* (x-axis). In panels (**B**) and (**C**), C or P in the CBC is the sum of C or P in 3PGA, DHAP, FBP, F6P, SBP, S7P, R5P, Xu5P+Ru5P and RuBP, summed C in CBC (without 3PGA and DHAP) is for all the above metabolites excluding 3PGA and DHAP, summed C in the CCM is the sum of C in aspartate, pyruvate, alanine and PEP (malate was excluded because the active pool could not be defined). Also shown is summed C in energy shuttle metabolites (3PGA and DHAP), and summed C in pyruvate and alanine as representative for the C in the CCM pool. These plots were made and presented as described in legend of Figure 5. Additional responses of metabolic traits versus *A_n_* are provided in Supplementary Figs. S7B-C and S8B-C. The original data are provided in Supplementary Dataset S2.

**Figure 7.**
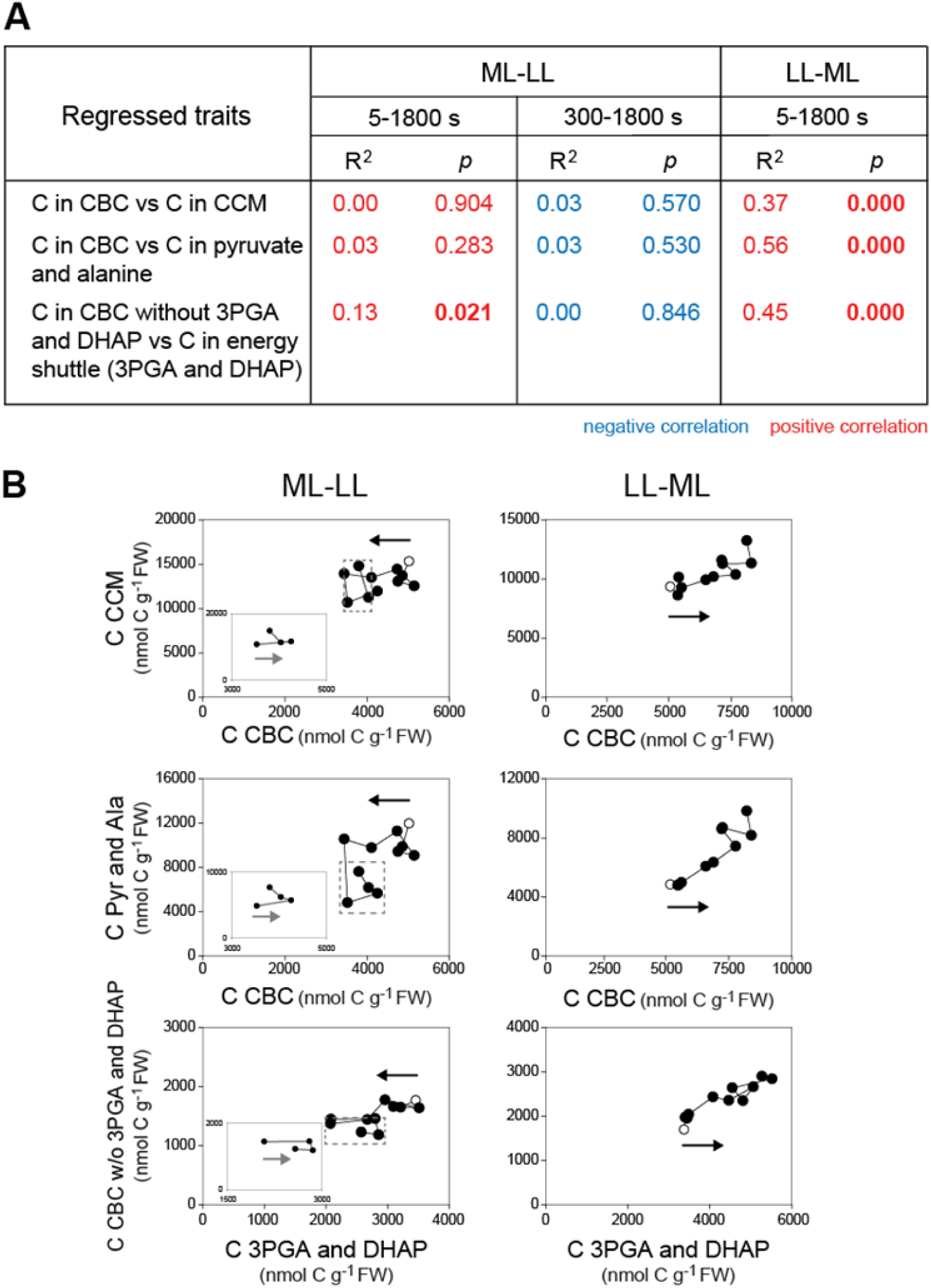
**Correlations between C pools in the CBC and in the CCM**. (**A**) Summary of R^2^, slope directions (slope d.) and *p* values. These were calculated by linear regressions using individual samples (*n*=3 to 5) at all points after the transition (time zero excluded). Additional values were calculated from 300 to 1800 s for the ML-LL transition. Positive and negative correlations are indicated by color (red and blue, respectively). Those with significant *p* values (*p* < 0.05) are in bold. (**B**) Individual regression plots in the ML-LL and LL-ML transitions. These plots were done and presented as described in legend of Figure 5. The original data are provided in Supplementary Dataset S2.

In the ML-LL transition (Fig. 5A; Supplementary Figs. S6A, S6C, S7B-C), a small subset of CBC intermediates, (3PGA, R5P, pentose-P), the starch synthesis intermediate ADPG, two photorespiratory intermediates (glycine, glycerate) and two CCM intermediates (PEP, pyruvate) correlated significantly with each other and clustered together, and also correlated significantly with and clustered with *A_n_*. Most of the metabolites in this small cluster had strong loadings in PC1 of the PC analysis (see Supplementary Fig. S2A). RuBP was only loosely associated with this group, and DHAP, FBP and SBP clustered elsewhere. Aspartate showed negative correlations with *A_n_*, as well as with many metabolites (Fig. 5A; Supplementary Fig. S5A). *A_n_* was significantly and positively correlated with the 3PGA/DHAP ratio (Fig. 6A), pointing to the balance between the supply and consumption of NADPH and/or ATP changing in favour of energy consumption during the adjustment to low irradiance. *A_n_* was weakly but significantly negatively correlated with the FBP/F6P ratio. *A_n_*was positively and significantly correlated with summed C and summed P in CBC intermediates. This included correlations both with the summed C in metabolites in the energy shuttle (3PGA, DHAP) as well as with summed C in other CBC metabolites (Fig. 6A). However, *A_n_* was unrelated to summed C in CCM intermediates, or summed C in pyruvate plus alanine. Furthermore, summed C in CBC intermediates and summed C in CCM intermediates or summed C in pyruvate plus alanine changed independently of each other (Fig. 7).

Plots of *A_n_* against individual metabolites or metabolite traits (Fig. 5B; Supplementary Figs. S6C and S7A) revealed rather diverse responses in the ML-LL transition, especially in the later phase after 250 s when *A_n_* was low but showed a slight recovery (inserts in Fig. 5B and Supplementary Fig. S7). The spread of values in the later part of the transition will distort some of the correlations performed over the entire time series. For this reason, additional analyses were performed on the time segment 300-1800 s (Supplementary Fig. S6C; these responses are considered in more detail below. Overall, the response of metabolism to a decrease in irradiance was complex and rather uncoordinated.

A different picture emerged for the LL-ML transition, where a much larger set of CBC (3PGA, DHAP, F6P, S7P, pentose-P), CCM (PEP, pyruvate, alanine) and photorespiratory (glycine) intermediates as well as 2OG clustered together, correlated significantly with each other, and also clustered with and correlated significantly with *A_n_* (Fig. 5A; Supplementary Figs. S6B-C). These metabolites had strong loadings in PC1 of the PC analysis (Supplementary Fig. S2B). RuBP also correlated with *A_n_* and most of the above metabolites. The slight separation of RuBP may reflect its rather constant level from about 60 s onwards as *A_n_* rose from 87 to about 130 nmol CO_2_ g^-1^ FW s^-1^ (Fig. 4A). SBP and, especially, FBP clustered elsewhere. Plots of selected metabolites against *A_n_* (Fig. 5C; Supplementary Figs. S6C, S8A) revealed a progressive increase in the level of many metabolites as *A_n_* rose, except that RuBP plateaued in the last half of the rise in *A_n_* (Fig. 5C; Supplementary Fig. S7A). Notable exceptions included 2PG that showed a rapid initial rise and subsequent fluctuations (Fig. 5C), and FBP and SBP that changed independently of *A_n_* (Supplementary Fig. S8A). Aspartate was negatively correlated with many metabolites (Fig. 5A; Supplementary Fig. S6B-C) and showed a progressive decrease as *A_n_* rose (Fig. 5C). *A_n_* was unrelated to the 3PGA/DHAP ratio but, as a consequence of the coordinated increase of many metabolites, was significantly and positively correlated with summed C and summed P in CBC intermediates, summed C in metabolites in the energy shuttle (3PGA, DHAP), summed C in all other CBC metabolites, summed C in pyruvate plus alanine, as well as summed C in all accessible CCM metabolites (malate was excluded for reason explained above) (Fig. 6). Furthermore, summed CBC metabolites correlated strongly and positively with summed CCM metabolites and with summed pyruvate plus alanine. Summed CBC metabolites excluding 3PGA and DHAP also correlated much more strongly with summed 3PGA plus DHAP in the LL-ML transition than in the ML-LL transition (Fig. 7). These analyses point to a coordinated increase in CBC, energy shuttle and CCM pool sizes in the LL-ML transition, in contrast to the less coordinated response in the ML-LL transition.

Some trends were common to both transitions (Fig. 6). This included a positive correlation of the summed CBC metabolite pool and its components with *A_n_* in both transitions, although the overall correlation was stronger in LL-ML than ML-LL. There was a strong correlation of the pyruvate/alanine ratio with *A_n_* in both transitions, consistent with the contribution of NADP-ME declining relative to other decarboxylases in low irradiance, and with this shift occurring gradually over the entire transition and being linked with the progressive changes in the rate of photosynthesis. The 2OG/glutamate ratio was strongly positively correlated with *A_n_* in the LL-ML transition and showed a weak positive trend with *A_n_* in the ML-LL transition, pointing to the balance between availability of amino and carboxylic groups being involved in adjustment to changed irradiance. This includes a gradual increase in glycine and aspartate during the ML-LL and decrease during the LL-ML transition (see Figs. 3A, 4A).

### Relationship between A_n_ and metabolite levels and metabolic traits in different phases of the transitions

As already noted, correlation analyses performed on an entire time series may not detect relationships in a part of the response, especially in ML-LL where *A_n_* falls and then recovers slightly. Visual inspection of Figs. 5B-C, Figs. 6B-C and Supplementary Fig. S7. revealed some relationships that were masked when regressions were performed across the entire transition (see also Figs. 3-4; Supplementary Figs. S3B and S4B).

In the ML-LL transition, the gradual decline of *A_n_* in the first 10-15 s after decreasing the light intensity was associated with a decline of DHAP (Supplementary Fig. S7A) and a high 3PGA/DHAP ratio (presumably reflecting consumption of DHAP to provide energy), a high pentose-P /RuBP ratio and a low RuBP/3PGA ratio (Supplementary Fig. S7B). The subsequent rapid drop in *A_n_* until about 120-250 s was accompanied by a decline in 3PGA, and a decline of the 3PGA/DHAP ratio (Supplementary Fig. S7B), reflecting the decline in energy consumption as *A_n_* fell. An increase of FBP and SBP levels (Supplementary Fig. S7A) and the FBP/F6P and SBP/S7P ratios Supplementary Fig. S7B) pointed to rapid inactivation of FBPase and SBPase. There was also a large decline of pyruvate (Supplementary Fig. S7A) pointing to loss of C from CCM metabolites.

As already mentioned, an especially striking feature of the MM-LL transition was the wide spread of values for metabolite levels and metabolic traits between ∼250 and 1800 s (Figs. 5B-C, 6B-C; Supplementary Figs. S7A-C). The slight recovery of *A_n_* in this time span (see Fig. 1C, Supplementary Fig. S1C) was accompanied by a decline in RuBP (Fig. 5B), a rise in 3PGA and PEP, a decline in FBP, a rise in SBP (from 300 s onwards) (Supplementary Fig. S7A), a decrease of the RuBP/3PGA ratio, an increase of the 3PGA/DHAP ratio, a decrease of the FBP/F6P ratio and an increase of the pentose-P/RuBP ratio (Supplementary Fig. S7B). There was a decline of summed C in CBC metabolites when 3PGA and DHAP were excluded (Fig 6B) mainly due to the decrease in RuBP. There was an increase of summed C in energy shuttle metabolites (3PGA plus DHAP) (Fig 6B). After a strong decrease until 300 sec, pyruvate levels showed a slight recovery (insert plot in Fig. 5B) that was accompanied by an increase in alanine (Supplementary Fig. S7A), in summed C in pyruvate plus alanine (Fig. 6B) and in summed C in all CCM intermediates (Supplementary Fig. S7C). Many of these changes were significant when regression analysis was performed on the 300-1800 s time segment (Supplementary Fig. S6C, also when the responses at 600, 1200 and 1800 s were tested for significance against 300s, Supplementary Fig. S3B). These observations point to the slight recovery of *A_n_* from ∼250 s on being associated, in the CBC, with a decrease of RuBP, increased activation of FBPase and, increased activity of PRK and Rubisco, whilst delivery of energy for 3PGA reduction becomes more restrictive. The latter presumably reflects increased energy consumption due to the slight recovery of *A_n_*. The slight recovery of *A_n_* was also associated with an overall rise in the levels of metabolites involved in intercellular energy and CCM shuttles. Overall, as *A_n_* rose from 39.7 to 41.8 nmol CO_2_ g^-1^ FW s^-1^ between 300 s and 1800s (Supplementary Fig. S1C), the combined pool of C in the CBC and CCM increased by 4764 nmol C g^-1^ FW s^-1^. This is equivalent to about 114 s of photosynthesis in LL, about 45 s of the difference in photosynthesis between ML and LL (Δ*A_n_*), and about 10% of the CO_2_ fixed during the entire recovery phase. or even more in the first part of the recovery, which was completed by 900-1000 s. Thus, there may be a trade-off between short-term carbon gain immediately after the switch at the expense of medium-term carbon gain, slightly later after the switch, which is restored by re-establishing shuttles at the timepoints furthest away from the switch. To do this, approximately 10% of the fixed C needs to be invested in building up CCM and CBC pools.

In the LL-ML transition, plots of *A_n_* against metabolites and metabolic traits reveal an immediate and significant decrease of the 3PGA/DHAP ratio (Supplementary Fig. S8B, for significance see Supplementary Fig. S3B). This reflects the rapid increase in provision of ATP and NADPH after an increase in irradiance. Between 5 and 120 s, by when *A_n_*had risen to about 116 nmol CO_2_ g^-1^ FW s^-1^, the 3PGA/DHAP ratio rose slightly (Supplementary Fig. S8B). This reveals that increasing energy consumption associated with rising *A_n_* leads to 3PGA reduction exerting a progressively larger restriction on CBC flux. The FBP/F6P ratio rose to a maximum at an intermediate *A_n_*of 75-89 nmol CO_2_ g^-1^ FW s^-1^ and then declined as *A_n_* rose further (Supplementary Fig. S8B, for significance see Supplementary Fig. S4B). The SBP/S7P ratio showed a similar and the pentose-P/Rubisco ratio an opposite trend (Supplementary Fig. S8B). This points to FBPase and, possibly, SBPase and PRK initially restricting CBC flux and this restriction being subsequently relaxed, presumably by activation of the enzymes. There was already a small increase in summed CBC as *A_n_* rose to 87 nmol CO_2_ g^-1^ FW s^-1^ (Supplementary Fig. S8B), mainly due to an increase in metabolites involved in RuBP regeneration and RuBP itself (Fig. 6C). Metabolites involved in the energy shuttle (3PGA, DHAP) showed little change until *A_n_* rose above 87 nmol CO_2_ g^-1^ FW s^-1^, after which the energy shuttle pool increased markedly (Fig. 6C; Supplementary Figs. 8A, 8C). The combined pyruvate and alanine pool increased slightly as *A_n_* rose to 87 nmol CO_2_ g^-1^ FW s^-1^and strongly as *A_n_* increased to the final value of about 130 nmol CO_2_ g^-1^ FW s^-1^ (Fig. 6C). Overall, the initial part of the rise of *A_n_* is associated with rapid build-up of CBC pools involved in RuBP regeneration and, probably, activation of CBC enzymes, and the second part is associated with a slow build-up of energy shuttle pools and CCM pools. Notably, this second part, which was equivalent to half of the total increase in *A_n_*, was not accompanied by a further increase in RuBP (Figs. 4A, 5C). Overall, the rise in *A_n_* from about 50 to 130 nmol CO_2_ g^-1^ FW s^-1^ was accompanied by an increase of summed C in the CBC, energy shuttle and CCM of about 7000 nmol C g^-1^ FW, which is equivalent to about 55 s of *A_n_* in ML, and considerably more of the lower *A_n_* that prevailed for the first part of the transition.

There was a weak positive correlation between the RuBP/2PG ratio and *A_n_* in the total transient, but somewhat above the significance cut-off (Fig. 6A, p = 0.068). This was mainly due to a preponderance of low ratios early after transfer to LL, and higher ratios later in the transition (Supplementary Fig. S8B).

## Discussion

Evidence is emerging for a larger loss of photosynthetic efficiency in C_4_ photosynthesis than C_3_ photosynthesis after sudden changes in irradiance (Kubásek et al. 2013; Li et al. 2021; Arce-Cubas et al. 2023a) with a more nuanced response under fluctuating light (Arce-Cubas et al. 2023b). These responses may reflect the complex topology of C_4_ metabolism. In C_3_ photosynthesis, CBC metabolite pools are small and change quickly and enzyme activation is quite fast, allowing a rapid increase in CBC flux after an increase in irradiance, whilst an almost instantaneous shortfall of ATP and NADPH rapidly inhibits CBC flux after a decrease in irradiance. Further factors could potentially come into play in C_4_ photosynthesis, including the challenge of coordinating fluxes in the CCM and the CBC, the need to balance energy provision and consumption in two cell types, and the time required to adjust the size of large metabolite pools that drive intercellular shuttles. Their impact of any given factor will depend on when and how quickly it becomes imbalanced and is corrected, and on how quickly the light intensity is changing. To provide information about the response of C_4_ photosynthetic metabolism to changes in irradiance, we carried out time-resolved measurements of *A_n_* and metabolite levels after a sudden increase or decrease in irradiance in maize. To reduce complications due to slow relaxation of energy dissipation, we worked in an irradiance range that was limiting for photosynthesis.

Our results reveal that C_4_ photosynthesis responds to sudden changes in irradiance in a complex manner, including delayed and in part uncoordinated responses in the CCM and the CBC. PC analysis of the global metabolic response pointed to the adjustment involving to several phases, with slow progressive changes of some metabolites and transient fluctuations of other metabolites. A similar picture emerged from scrutiny of the responses of individual metabolites, metabolite ratios and summed pools in the energy shuttle and the CCM (Figs, 3-4; Supplementary Figs. S3-S4), as well as from analysis of the relationship between *A_n_* and metabolite levels or metabolic traits (Figs, 5-6; Supplementary Figs. S6-S8). The responses to a decrease and increase of irradiance are schematically summarized in Fig. 8. The response after a decrease in irradiance was especially complex (Fig. 8A). Briefly, relatively high rates of photosynthesis were maintained for a short time, followed by a drop to a minimum after ∼250 s and a small recovery over the next ∼1400 s. After an increase in irradiance (Fig. 8B) the increase in *A_n_* took several minutes. Analysis of the temporal kinetics of *A_n_* as well as PC analysis of the metabolite profile indicated that this rise included an initial short delay, followed by a rapid rise until about 90 s that accounted for about 80% of the gain in *A_n_*, and a slower rise that accounted for about 20% of the gain in *A_n_*.

**Figure 8.**
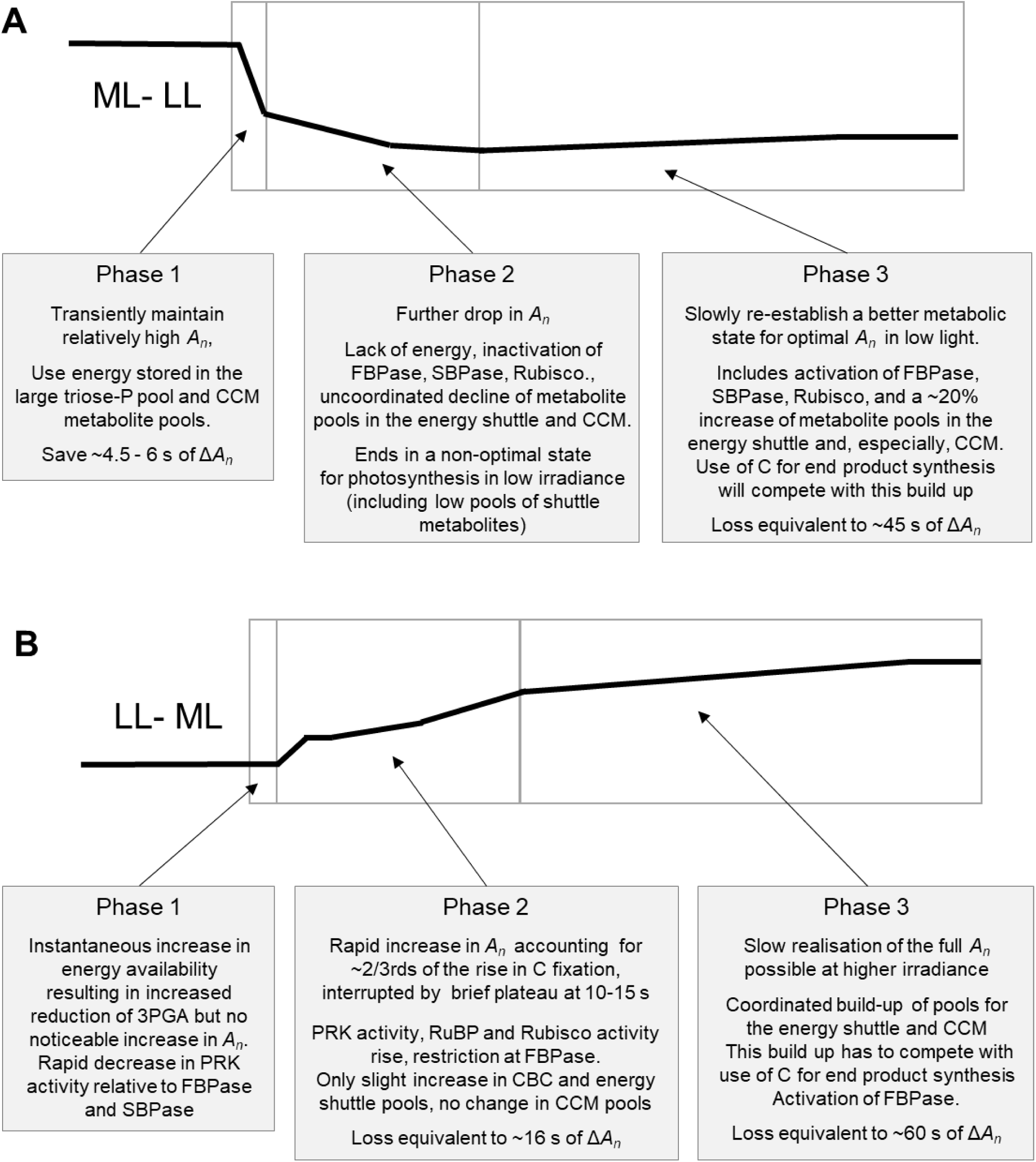
**Schematic summary of the adjustment of C_4_ photosynthesis in maize to a change in irradiance in the non-saturating range**. (**A**) Adjustment to a decrease in irradiance, (**B**) adjustment to an increase in irradiance: The displays show schematically the change of *A_n_*in each phase and the boxes summarise the main accompanying changes in metabolism. The loss of *A_n_* in a given phase was estimated as follows. In the ML-LL transition, *A_n_* above (phase 1) or below (phase 3) steady state *A_n_* in LL was integrated over the phase, and subtracted from the Δ*A_n_* (the difference between *A_n_* in ML and LL). In the LL-ML transition *A_n_* below steady state *A_n_* in ML was integrated over phase 2 or phase 3 and subtracted from Δ*A_n_*.

The slow and complex response of *A_n_* in single transitions are reminiscent of those reported in earlier studies of the response of C_4_ photosynthesis to sudden transitions or to fluctuating light, whereby in the latter case the starting point depends on the frequency of the fluctuation and differs from that after moving from one steady irradiance to another. In studies where irradiance was decreased, C assimilation continued for a time after darkening (Laisk and Edwards 1998), after imposing and then removing a short period of high light (Krall and Pearcy 1993) or after switching to low light in a fluctuating regime (Lee et al. 2022; Arce-Cubas et al. 2023b; Sales et al. 2025). This contrasts with C_3_ photosynthesis, where in 21% O_2_ there is a burst of CO_2_ is released from photorespiration intermediates after darkening or a decrease in illumination, and in 2% O_2_ a very a rapid decline of *A_n_* (Arce-Cubas et al. 2023b).A transient trough of A_n_ followed by a slight recovery has been reported for a sudden high to low light transition with maize (Doncaster et al. 1989), and in some but not other studies with fluctuating light (for three C_4_ NADP-ME species but not maize (Lee et al. 2022), for one of three C_4_ species (Arce-Cubas et al. (2023b) but not maize and was absent for several species (Li et al. 2021). This may reflect differences in poising, or whether the time in low light was long enough to capture the trough. For treatments where irradiance was increased, earlier studies have reported a slow rise of *A_n_* over minutes after sudden illumination (Furbank and Leegood 1984; Usuda 1985; Arce-Cubas et al. 2023a) or a sudden switch from low to high light (Kubasek et al. 2013). Under fluctuating light, however, the recovery of *A_n_* in three C_4_ species was not consistently slower than in phylogenetically-paired C_3_ species (Arce-Cubas et al. 2023b). Earlier studies also reported complex transients immediately after a switch from darkness to light (Laisk and Edwards 1998) or from low light to high light (Krall and Pearcy 1993). Some studies of C_4_ species under fluctuating light have reported that the rise in *A_n_* after switching to high light includes a plateau after about 60 s (Lee et al. 2022; Arce-Cubas et al. 2023b).

The metabolic events following the imposition of a sudden increase or a decrease in irradiance in maize are summarized in Fig. 8. The next two sections and the Supplementary Text Sections 2-6 use this dataset to explore possible reasons for the complex response of C_4_ photosynthesis to changes in irradiance,

### Response of the CBC and CCM to a sudden decrease in irradiance

Our results confirm the prediction of Stitt and Zhu (2014) that the large pools of metabolites in the energy shuttle and the CCM temporarily buffer C_4_ photosynthesis against a sudden shortfall of ATP and NADPH from the light reactions. Crucially, the observed changes in metabolite levels in our study, especially the decline of DHAP, were quantitatively large enough to supply the energy required to support the temporarily elevated rate of CO_2_ fixation over that in steady state low light (see Supplementary Text Section 2.1).

The subsequent fall in *A_n_* was primarily due to a shortfall in energy and was accompanied by a decline of the pools of CBC intermediates, decreased activation of FBPase, SBPase and Rubisco, as well as a decline in the levels of metabolites in the energy shuttle (Supplementary Text Section 2.1) as well as of pyruvate due to exchange of C between the CCM and CBC pools (Supplementary Text Section 3. It ended after ∼200-250 s with a transient trough in *A_n_*. This was followed by a slow partial recovery of *A_n_*.

Several factors might in principle contribute to the transient trough and subsequent slight recovery of *A_n_*. One possibility is that the rapid decline in CO_2_ fixation leads to sub-optimal poising of CBC metabolite levels and enzyme activities. However, inspection of changes in metabolite levels and metabolite ratios indicates that, except for 3PGA and DHAP, CBC metabolites do not rise during the partial recovery of *A_n_*(Figs. 3, 6A-B; Supplementary Figs. S3B, S6C, S7). There may be some activation of FBPase and Rubisco (Supplementary Fig. 3B) and the latter may partly explain why RuBP levels are unaltered or fall slightly during the partial recovery of *A_n_* (Fig. 3A; Supplementary Fig. S7A, but see below for further discussion). However, adjustment in the CBC is unlikely to make a major contribution to the slow recovery of *A_n_*. As in C_3_ photosynthesis, CBC metabolite pools are small and turn over rapidly (Arrivault et al. 2017) and activation of CBC enzymes adjusts in a relatively short time as in C_3_ species (Doncaster et al. 1989). Another possibility would be that the trough and gradual recovery is due to slow relaxation of energy dissipation. However, this seems unlikely because the experiments were carried out in the limiting light range where large changes in energy dissipation are unlikely and the 3PGA/DHAP ratio rose during the recovery (Fig. 3B; Supplementary Figs. S3B, S6C) rather than falling as would be expected if delivery of ATP and NADPH from the light reactions was increasing. The recovery also cannot be due to slow adjustment of stomatal conductance, which together with *C_i_* decreased in this time (Supplementary Fig. S1D).

Based on these prior considerations, the major explanation for the trough of *A_n_* seems to be that, during the rapid decline of *A_n_*, metabolite pools in the 3PGA-DHAP energy shuttle and, especially, the CCM decline to below those required for optimal operation at the new low irradiance (see above and Supplementary Text Section 2.1). Low irradiance leads via a reduced rate of CO_2_ fixation to a decline in CBC metabolite levels (see Introduction). As the CBC and CCM are connected via the near-equilibrium reactions of phosphoglycerate kinase and enolase, a decrease in 3PGA will divert PEP away from PEPC, draining C from the CCM (for details see Supplementary Text Section 3). This explanation is supported by the decrease of the 3PGA/PEP ratio at 60-120 s (Figure 3A, Supplementary Text Sections 2.1 and 3.2). Furthermore, metabolite pools drive intercellular shuttles less efficiently in low irradiance than high irradiance (Medeiros et al. 2022). This will further exacerbate the impact of the depletion of these pools on *A_n_*after a sudden decrease in irradiance.

The subsequent gradual recovery of *A_n_* was accompanied by a rise in the level of energy shuttle intermediates (Figs. 3A, 6B) and a somewhat delayed rise in the levels of pyruvate, alanine, aspartate (Fig 3A, Supplementary Fig S7A) and total C in CCM metabolites (Fig. 3C; Supplementary Fig. S7C) (see Fig. 6A and for significance Supplementary Figs. 3B, 6C, also Supplementary Text Section 2.1). Overall, the C that accumulated in the CBC, energy and CCM pools was equivalent to about 10% of all the CO_2_ fixed during the recovery phase, and more in the first part of the recovery, with most of this being due to build-up of CCM pools (see Supplementary Text Section 2.1). It might be noted that a gradual improvement in operation of the CCM in low irradiance provides an explanation for why *A_n_* rises independently of changes in RuBP levels and whilst stomatal conductance and *C_i_* are decreasing.

Further factors may also contribute to the delay until the CCM operates with full efficiency at low irradiance. One is the somewhat uncoordinated build-up of the CBC and CCM pools (Fig. 7, for more discussion see Supplementary Text Section 3). Another is that low irradiance restricts decarboxylation by NADP-ME (Medeiros et al. 2022), leading to a shift of flux towards other decarboxylases and an associated change in which 4C and 3C metabolites move between the BSC and the MC, in particular, an increased contribution of aspartate and alanine. The slow and complex responses of amino acid pools to a decrease in irradiance (see Figs. 3A-C; Supplementary Fig. S7A; Supplementary Text Section 2.1) may contribute to the initial inhibition and slow recovery of *A_n_*.

The over-depletion and slow recovery of the energy shuttle and CCM pools in the ML-LL transition is not due to large pool size *per se*. It is a consequence of how the large pools impact poising of C gain with end product synthesis. Due to the topology of the CBC, use of fixed C for end product synthesis must be tightly regulated to maintain CBC intermediates in a concentration range that allows rapid regeneration of RuBP (Edwards and Walker 1983; Heldt et al. 2021; Stitt et al. 2010, 2021). This can be illustrated by considering synthesis of sucrose, the main end product of photosynthesis. In C_3_ plants, triose-P are exported from the plastids to the cytosol, where they are converted to sucrose. Fructose 2,6-bisphosphate (Fru2,6BP) interacts with other metabolites in a highly cooperative network to strongly restrict cytosolic FBPase activity in response to a small decline in the levels of CBC metabolites (Herzog et al. 1984; Stitt, 1990; Stitt et al. 2010, 2021). This network allows rapid inhibition of triose-P consumption for sucrose synthesis after a decrease in irradiance, whilst CBC metabolites are kept at relatively high levels (Stitt et al. 1983; Stitt 1990; Stitt et al. 2021). In maize, sucrose is synthesized in the cytosol of the MC (Furbank et al. 1985) and this process must be coordinated with the maintenance of high triose-P concentrations in the MC to drive their diffusion back to the BSC (Leegood 1985; Stitt and Heldt 1985a; Arrivault et al. 2017). High triose-P concentrations can be maintained in the MC because, compared to the C_3_ cytosolic F1,6BPase, the maize enzyme has a circa 10-fold higher K_m_ for FBP (Stitt and Heldt 1985b). Thus, whilst a decrease in C fixation and the associated partial inhibition of sucrose synthesis is associated with a decrease in triose-P levels, this occurs in a much higher concentration range in maize than in C_3_ species (Stitt and Heldt 1985a, 1985b), compare also steady state DHAP levels in C_4_ and C_3_ plants in Arrivault et al. (2019). Crucially, the larger absolute size of the triose-P pool and associated pools like 3PGA means that a given relative change in triose-P concentration is associated with a much larger change of absolute pool size in maize than in a C_3_ plant. Further, as 3PGA is linked via the reversible phosphoglycerate mutase and enolase reactions to PEP (Arrivault et al. 2017, see also Supplementary Text Section 3), a decline in triose-P and 3PGA levels leads to drainage of C from the large pools of CCM intermediates. Thus, when photosynthetic rate falls, any delay in restricting flux at cytosolic F1,6BPase will lead to a much larger absolute drainage of C from photosynthetic metabolism in a C_4_ plant like maize than in a C_3_ plant. Consequently, it takes longer to reverse this process and build up the pools again.

In experiments where maize was transferred from stable illumination at 1700 to 144 µmol m^-2^ s^-1^ Doncaster et al. (1989), a slight recovery of *A_n_* from about 2 min onwards was accompanied by a slight decline of the 3PGA/DHAP ratio, a decline of the FBP/F6P ratio and an increase of RuBP indicating improved poising in the CBC including activation of plastidic FBPase. Some specific responses differed between our study and that of Doncaster et al. (1989) for example, in their study pyruvate levels rose for the first 2 min after the reduction in irradiance and then declined, whereas in our study pyruvate declined immediately rather than showing an initial peak. This may reflect differences in the regulation of NADP-ME or PPDK when irradiance is reduced from a very high level or a moderate level.

Taken together, immediately after transferring maize from stable high to low irradiance, substantially higher *A_n_* is maintained than in steady state low light. However, by 200-250 s the metabolic network transitions into a state that is not optimally poised for effective use of the low irradiance, and it takes several minutes for a better poise to be established. The gain of C fixation due to high *A_n_* immediately after reducing irradiance was equivalent to about 6 s of Δ*A_n_*, (the difference between *A_n_*in ML and LL) (Fig. 8A). This is more than cancelled out by the loss of C fixation due to the trough and slow recovery, which was equivalent to about 45 s of Δ*A_n._* (Fig. 8A). Hence, whilst our experimental study confirms the prediction of Stitt and Zhu (2014) that large metabolite pools may temporarily buffer C_4_ photosynthesis against a sudden reduction in irradiance, it also reveals that there can be a larger loss of photosynthetic efficiency due to sub-optimal responses during the subsequent adjustment to low irradiance. This might in part result from metabolic transformations that deliver energy to support high *A_n_*immediately after the decrease in irradiance. However, these will mainly move C between different metabolites in the CBC and the CCM. The trough is due to depletion of the overall pool of C in the CBC and CCM pools to below the level in steady state low light, due to slow inactivation of end-product synthesis. That said, under rapid light fluctuations that often predominate in dense C_4_ grass canopies, gains due to short-term buffering of *A_n_* will probably outweigh the losses that occur after a switch from high light to prolonged low light.

### Response of the CBC and CCM to an increase in irradiance

Immediately after a sudden increase in irradiance the 3PGA/DHAP ratio decreased, as expected from the increased availability of NADPH and ATP (Fig. 4A, Supplementary Fig. 4B,). Starting at about 6 s, there was a rapid rise in *A_n_*, a transient plateau at around 11-15 s and a second rapid rise until 90. The transient plateau and is probably linked to delayed activation of the CCM and CBC, including uncoordinated responses that may lead to transient back-leakage of CO_2_ (see Supplementary Text Sections 2.2, and 5). Overall, between 6 and 90 s, the rise in *A_n_* was associated with an overall increase in CBC metabolite pools (Figs. 4A, 6C; Supplementary Figs. S4B, S6B) and transient changes indicative of enzyme regulation (see Supplementary Text Section 2.2). From 90 s onwards, *A_n_*continued to rise but more slowly. This slow rise represented about one quarter of the total gain in A*_n_*. It was accompanied by a rise of the levels of 3PGA and DHAP, PEP, pyruvate and alanine, Importantly, the gradual rise in *A_n_* correlated strongly with the increase in the pool size of the energy shuttle and CCM (Figs. 6A, 6C; Supplementary Figs. S4B, S8C).

Overall over 1800 s, summed C in the CBC, energy shuttle and CCM increased by about 7000 nmol C g^-1^ FW, which is equivalent to about 60 s of *A_n_*in ML, and considerably more of the lower *A_n_* that prevailed in the first part of the transition (for more details see Supplementary Text Section 2.2). The increase in energy shuttle metabolite and CCM pool size was rather coordinated, with the pools being strongly correlated throughout the adjustment (Figs. 4C, 6, 7). This coordinated rise reveals that neither the energy shuttle nor the CCM is prioritized when maize is adjusting to a sudden increase in irradiance (see also Supplementary Text Sections 2.2 and 3). This contrasts markedly with the rather uncoordinated response during the ML-LL transition. That said, the rise of metabolite pools after an increase in light intensity can be divided into two phases; an increase in CBC metabolites (excluding 3PGA and DHAP) that predominated as *A_n_* rose about 50 to 90 nmol CO_2_ g^-1^ FW s^-1^ in the first 90 s, and an increase of energy shuttle and CCM metabolites that predominated during the subsequent gradual increase of *A_n_* from 90 to 130 nmol CO_2_ g^-1^ FW s^-1^. The latter provides strong correlative evidence that the second and slower part of the rise in *A_n_* is closely linked with the gradual buildup of the large metabolites pools that drive intercellular shuttles.

The initial increase in CBC pool size and the subsequent rise in energy shuttle and CCM pool size requires tight regulation of end-product synthesis. G6P and UDPG fell until about 60 s, but then rose (Supplementary Figs. S4A-B, pointing to an initial restriction at cytosolic FBPase that is subsequently relaxed, after which sucrose synthesis competes for newly fixed C with the build-up of pools in the energy shuttle and CCM. A somewhat delayed response of the cytosolic FBPase is also seen in C_3_ species, and is at least partly due to a short delay until the Fru2,6BP levels fall (Stitt 1990; Stitt et al. 2010; 2021). Incidentally, the initial increase and subsequent decrease of ADPG (Fig. 4A; Supplementary Fig. S4B) is also consistent with delayed stimulation of flux at cytosolic FBPase.

Interestingly, in a study under fluctuating light Li et al. (2021) observed that, after switching from low to high light, C_3_ species showed a rapid rise of *A_n_* within 1 min to a value similar to or only slightly lower than that attained in steady state high light. In contrast, in C_4_ species, the response was biphasic with a rapid initial rise, followed by a much slower gradual increase. This pattern has been seen in many NADP-ME and NAD-ME species (see also Lee et al. 2023), although not in two PEPCK species (Lee et al. 2023; Arce-Cubas et al. 2023) (see Supplementary Text Section 2.2) In many cases the steady-state rate in high light was not reached within 2 min and, in the maize LL-ML transition, full recovery required more than 8 min. Thus, the rapid initial rise and slower gradual rise seen in maize in a LL-ML transition (Fig. 1B; Supplementary Fig. S1E) is a general feature of NADP-ME and NAD-ME subtypes, although not necessarily PECK subtypes.

Wang et al. (2021) modelled the response of C_4_ photosynthesis to an increase in irradiance, in this case from darkness. Their model predicted that the delay until maximum *A_n_* was achieved was relatively short, and mainly due to limitation by Rubisco activase, the PPDK regulatory protein and stomatal conductance. Although their model correctly predicted slow build-up of these pools, this build-up continued long after modelled *A_n_* had reached a new maximum value, leading them to conclude that the delay is not due to slow build-up of energy shuttle and CCM metabolite pools. In earlier experimental studies of dark-light transitions Leegood and Furbank (1984) and Usuda et al. (1985) observed a closer match between the increase of *A_n_* and the increase in 3PGA, DHAP and metabolites like pyruvate that are involved in the CCM. This resembles our study of a transition to higher light. As noted by Wang et al. (2014b), parameterization of the resistance to intercellular movement of metabolites is challenging. It is possible that the parameterization in Wang et al. (2021) underestimated the resistance to intercellular movement of metabolites and, hence, underestimated the contribution of a slow build-up of metabolite pools to the response of *A_n_*after an increase in irradiance.

As already mentioned, the model of Wang et al. (2021) predicted that activation of Rubisco activase and the PPDK regulatory protein are major factors in the loss of photosynthetic efficiency during dark to high light transitions. Our analyses of metabolites in a transition between low and moderate light provide experimental support for the idea that slow Rubisco activation restricts *A_n_* in the first phase of the response after a switch from low to moderate irradiance (see above, Figs. 4A, 5C; Supplementary Fig. S8A). In earlier studies of a darkness to high light transition, RuBP rose to a peak at 2 min and then declined as *A_n_*rose further (Usuda 1985), again pointing to delayed activation of Rubisco. Our data also support the idea that the rise of *A_n_* is initially restricted by slow activation of PPDK. The marked decrease of PEP and slight but significant increase of pyruvate at 5-15 s after a sudden increase in irradiance (Figs. 4A, 5B; Supplementary Fig. S4B), points to consumption of PEP by PEPC outstripping delivery of PEP by PPDK. In earlier studies of a darkness to high light transition, pyruvate initially declined, before starting to rise from about 2 min on (Furbank and Leegood 1984; Usuda 1985). This initial drop may reflect the difference between a transition starting from darkness and from low light, when PEPC and NADP-ME may already be slightly activated. The subsequent rise of pyruvate in these earlier studies started at about the same time as in our LL-ML transition.

The model of Wang et al. (2021) also predicted that slow adjustment of *g* restricts photosynthesis during a dark to high light transition. Slow adjustment of *g* leading to a minimum of *C_i_* was seen ∼60 s after a sudden increase in light intensity (Fig. 1B). The resulting gradual rise in *C_i_* might contribute to the gradual recovery of *A_n_*. *Ci* fell to a minimum (∼80 ppm) after 60-100 s before slowly rising to about 150 ppm (Fig. 1B; Supplementary Fig. S1B). As the former may slightly restrict photosynthesis in maize (Carmo-Silva et al. 2008; Bellasio et al. 2016), part of the subsequent slow gain of *A_n_* may be related to gradual stomatal opening which supports the CCM by allowing fuller saturation of PEPC with bicarbonate. It might be noted that measurement of *g* is technically challenging in transients, and the changes be underestimated due to dampening effects of water vapor interactions with cuvette walls.

### Contribution of different decarboxylation routes

In addition to NADP-ME, maize BSC contains PEPCK (Walker et al. 1997; Supplementary Text Section 4). Analyses of metabolite levels in high and low irradiance (Usuda 1987; Leegood and von Caemmerer 1989) and after a switch from high to low irradiance (Doncaster et al. 1990) as well as ^13^CO_2_ labelling kinetics in steady state low and high light (Medeiros et al. 2022) point to the contribution of the aspartate-based PEPCK route increasing in low irradiance. In our study, aspartate showed a strikingly different pattern to most other metabolites (Fig. 5; Supplementary Fig. S6), rising during the ML-LL transition (Fig. 3A; Supplementary Fig S3B) and decreasing during the LL-ML transition (Fig. 4A; Supplementary Fig. S4B). This occurred from ∼120 s onwards, pointing to a gradual increase in the contribution of PEPCK during adjustment to lower light, and a gradual decrease during adjustment to higher light (see also Supplementary Test Section 4).

Movement of aspartate from the MC to the BSC needs to be coupled to movement of an amino acid back to the MC. In the most parsimonious pathway, this would be alanine (Weissman et al. 2016; Bräutigam et al. 2018). However, in the ML-LL and LL-ML transitions aspartate and alanine levels were often negatively correlated (Figs. 3A, 4A, 5A; Supplementary Fig. SA-B6, for details see Supplementary Test Section 4). This argues against operation of a tight shuttle between aspartate and alanine, which would require parallel changes of the two shuttle partners. Alternatively, nitrogen stoichiometry could be maintained by a secondary intercellular shuttle, for example, between glutamate and 2OG. This idea is supported by the direction and timing of changes of 2OG and glutamate (Supplementary Text Section 4). Interestingly, the metabolic traits that changed during optimisation of C_4_ photosynthesis between C_4_-like and true C_4_ species in the *Flaveria* genus included an increase in the levels of 2OG and glutamate (Borghi et al. 2019). A secondary shuttle would add flexibility to C_4_ photosynthesis.

The increased contribution of PEPCK in low irradiance may reflect multiple factors. On the one hand, any shortfall of NADPH in the MC would restrict conversion of OAA to malate, both due to mass action and because low NADPH/NADP ratios inhibit thioredoxin-activation of NADP-MDH (Ashton and Hatch 1983; Rebeille and Hatch, 1986; Ashton et al. 1990). In the BSC, NADP-ME activity may be restricted by lack of demand for the reaction products (Bräutigam et al. 2018) and incomplete post-translational activation (Bovdilova et al. 2018). On the other hand, maize PEPCK is not subject to post-translational regulation (Wingler et al. 2019; Walker et al. 2002) and is presumably as active in low as in high irradiance. Operation of PEPC requires movement of PEP from the BSC back to the MC. In principle, this could occur as diffusion of PEP or, alternatively, as diffusion of PGA with enolase and phosphoglycerate mutase converting PEP to 3PGA in the BSC and 3PGA to PEP in the MC (see Supplementary Text, Section 3). PEP levels and/or enolase and phosphoglycerate mutase activity may suffice to support the low rate of movement required in low light but become restrictive under high light (see also Supplementary Text, Section 4).

### Metabolite analysis point to rapid perturbation and slow adjustment of the CO_2_ concentration in the bundle sheath

Changes in irradiance might lead to a temporary imbalance between the concentration of CO_2_ by the CCM and its utilization by Rubisco (Furbank et al. 1990; von Caemmerer 2000; Kromdijk et al. 2014; Wang et al. 2024). An excess of CO_2_ consumption over CO_2_ influx will lead to a decrease in C_BSC_ and an increase in the rate of RuBP oxygenation relative to RuBP carboxylation, whilst an excess of CO_2_ influx over CO_2_ consumption will lead to an increase in C_BSC_ and increased back-leakage of CO_2_. Our measurements of 2PG provide a qualitative proxy for the rate of RuBP oxygenation, and the relationship between the RuBP/2PG ratio and the RuBP/3PGA ratio provides a qualitative proxy for the relative rates of RuBP oxygenation and RuBP carboxylation (for more details see Supplementary Text Section 4).

Immediately following a decrease in irradiance, there was a significant >2-fold decrease in the level of 2PG (Fig. 3A; Supplementary Fig. S3B), a decline of the RuBP/3PGA ratio and an increase of the RuBP/2PG ratio. Conversely, following an increase in irradiance, there was a significant increase of 2PG (Fig. 4A; Supplementary Fig 8A), an increase of the RuBP/3PGA ratio and a decrease of the RuBP/2PG ratio (Fig. 4B; Supplementary Fig. S8B). These observations point to a transient increase in carboxylation relative to oxygenation after a decrease in irradiance, and a transient decrease in carboxylation relative to oxygenation after an increase in irradiance. They are consistent with C_BSC_ increasing after a decrease in light intensity, and C_BSC_ decreasing after an increase in irradiance, with accompanying changes in back-leakage of CO_2_.

We also asked if our metabolite analyses provide information about how quickly these changes are reversed (see also Supplementary Text Section 5). The 2PG level and RuBP/3PGA and RuBP/2PG ratios did not shown consistent changes later in the transients. Another approach to obtain indirect information about C_BSC_ is to compare *A_n_* with RuBP levels, under the assumption that they should correlate unless further factors are influencing Rubisco activity, of which one is C_BSC_. In the ML-LL transient, the 7% recovery of *A_n_* from ∼250 s onwards was accompanied by a large significant decline in RuBP levels (Fig. 3A; Supplementary Figs. S3B, S6C, S7A), whilst other CCM metabolites rose (see above). These observations indicate that C_BSC_ may continue to fall until ∼250 s and subsequently recover as CCM operation is optimized to low irradiance. (Figs. 4A, 4C, 6C; Supplementary Figs. S8A, S8C). In the LL-ML, from about 120 s onwards, *A_n_* rose (from about 86 to 140 nmol CO_2_ g^-1^FW s^-1^) without any further increase in RuBP levels (Figs. 4A, 5C), consistent with a gradual increase in CO_2_ pumping and C_BSC_.

Photorespiration contributes to photosynthetic response transients in C_3_ species by serving as a source of C to buffer pools or by sequestering C from the CBC cycle (Fu et al. 2023). Glycine fell in the ML-LL transition and rose in the LL-ML transition, but the magnitude of the changes was too small to make more than a minor contribution to the dynamics of CBC and CCM metabolite pools, or *A_n_* (see Supplementary Text Section 6).

In conclusion, several sub-processes in metabolism modify photosynthetic efficiency of maize during sudden transitions in light intensity. After a decrease in irradiance, photosynthesis is transiently buffered by energy stored in large pools of metabolites that are involved in intercellular shuttles. However, photosynthesis subsequently declines to a trough and it takes several minutes to recover to the steady rate in low light. One reason for the trough and slow recovery is that the large pools of metabolites involved in intercellular shuttles run down too far and it takes time to build them up again. After an increase in irradiance, it takes several minutes to establish a steady and higher rate of photosynthesis. Whilst the first part of this delay is associated with building-up pools of CBC metabolites and activating enzymes, the second part is mainly due to the need to build-up the large pools of metabolites required to drive intercellular energy and CCM shuttles. In addition, transient imbalances between the concentration and utilization of CO_2_ can lead to higher photorespiration or back-leakage of CO_2_ after an increase and decrease in irradiance, respectively. The impact of these responses and trade-offs between them under fluctuating light will depend, among other things, on the frequency of the fluctuations. When rapid fluctuations are experienced, as may often be the case in a canopy in the field, the gains immediately after the drop in irradiance may outweigh losses after longer exposure to low light. Furthermore, the supra-steady state rates maintained immediately after the decrease in light intensity may be partly linked to the suboptimal state underlying the later trough of photosynthesis. That said, the metabolic transformations that generate energy and support *A_n_* immediately after a drop in light intensity and the slower depletion of the overall pools of CBC and CCM metabolites are separate processes. Photosynthetic efficiency in fluctuating regimes might be enhanced by enhancing the ability to rapidly regulate end-product synthesis and maintain higher metabolite levels at low irradiance, both to avoid a trough of *A_n_* after a decrease in irradiance and to allow faster recovery of *A_n_* when irradiance increases.

## MATERIALS AND METHODS

### Chemicals

Chemicals were from Sigma-Aldrich (Darmstadt, Germany; www.sigmaaldrich.com), Roche Applied Science (Mannheim, Germany; lifescience.roche.com) or Merck (Darmstadt, Germany; www.merckmillipore.com).

### Plant growth

For metabolic sampling and gas exchange parameters recorded at 6 s intervals, maize (*Zea mays* L. cv. B73) seeds were germinated in darkness in petri dishes on moistened filter paper (3 days, 28°C), transferred to soil (Einheitserde Typ Topf [Naturton, white peat, wood fibers] mixed 2:1 with quartz sand; www.einheitserde.de) in 10-cm diameter pots, grown for 5 days under 16/8 h day/night cycles (irradiance 105 µmol photons m^-2^ s^-1^, 22/18°C, 70% relative humidity) and then under 14/10 h day/night cycles (irradiance 550 µmol photons m^-2^ s^-1^, 29/22°C, 65% relative humidity).

For gas exchange parameters recorded every second, maize (*Zea mays* L. cv. B73) seeds were sown in Levington advance F2 compost (Scotts, Ipswich, UK) in seed trays, and germinated under 14/10 h day/night cycles (irradiance 550 µmol photons m^-2^ s^-1^, 28/20°C, 65% relative humidity). After one week, seedlings were transplanted to 1.3 L pots, containing a mixture of 2:2:1 of Levington M3 compost (Scotts, Ipswich, UK): Top Soil (Westland, Dungannon, UK): Perlite 2.0-5.0 mm (Sinclair, Ellesmere Port, UK). Each pot was supplemented with 3.5 g L^-1^ of magnesium salts (Scotts Miracle-Gro, Marysville, OH, USA), and 7 g L^-1^ of garden lime (Needham Chalks, Suffolk, UK) and moved to a cabinet set to temperature conditions of 29°C/22°C day/night. After a week, a 5 g osmocote fertilizer tablet (14N-8P:11K+2MgO+trace elements; Osmocote Exact, ICL, Saltburn, UK) was added per pot.

For both growth conditions, plants were well-watered. The fourth fully elongated leaves of three-week plants were used.

### Gas exchange

For gas exchange parameters recorded every second, plants were transferred to a Percival E-41HO controlled environment chamber (Perry, IA, USA) set at 22°C on the evening before measurements. In the morning, lights were turned on and irradiance inside the chamber was kept at 550 μmol m^-2^ s^-1^ (ML) or 160 μmol m^-2^ s^-1^ (LL), to perform the ML-LL or LL-ML transition experiments, respectively. Each irradiance transition was performed in consecutive days. Gas exchange parameters were measured using a Li-6800 portable infrared gas analyser (IRGA) system (software version 1.4.22, LI-COR, Lincoln, Ne, USA) with a 6 cm^2^ chamber. Air CO_2_ concentration in the reference analyzer was controlled at 400 μmol mol^-1^, heat exchanger temperature at 25°C, and relative humidity at 65%, and the relative proportion of blue light was set to 10% PPFD. Flow rate was set to 600 µmol s^-1^. Gas exchange parameters were logged every second. Measurements started about 4h into the light period. For the ML-LL transition, leaves were measured for 15 min at 550 μmol m^-2^ s^-1^, then light inside the chamber was switched to 160 μmol m^-2^ s^-1^, and gas exchange parameters were logged for another 30 min. For the LL-ML transition, leaves were measured for 15 min at 160 μmol m^-2^ s^-1^, then light inside the chamber was switched to 550 μmol m^2^ s^1^, and parameters logged for another 30 min. As gas exchange measurements during and shortly after the light switch strongly violate the steady state assumption underlying default rate equations, dynamic equations were implemented (Saathoff and Welles, 2021). The dynamics of water vapour fluxes are less accurately measured than CO_2,_ mainly due to absorption/desorption on cuvette walls which dampen the responses and their derivatives (i.e., g_s_ and Ci).

Gas exchange parameters recorded at 6 s intervals were performed with plants in the growth chamber without shade (550 µmol m^-2^ s^-1^, ML) or with plants under a large polystyrene plate to reduce incident irradiance (160 µmol m^-2^ s^-1^, LL). Measurements were performed on two days for ML-LL and one day for LL-ML. Gas exchange parameters were measured using an open-flow infrared gas exchange analyser system (LI-6400XT, LI-COR Inc. Lincoln, NE, USA) equipped with an integrated fluorescence chamber head (LI-6400-40, 2 cm^2^ leaf chamber; LI-COR Inc. Lincoln, NE). The photosynthetically active photon flux density (PPFD) inside the chamber was kept at 550 µmol m^-2^ s^-1^ (ML) or 160 µmol m^-2^ s^-1^ (LL). Blue light was set to 10% PPFD, CO_2_ was kept at 400 µmol mol^-1^, leaf temperature at 29°C and relative humidity at 65%. Gas exchange parameters were recorded at 6 s intervals. Measurements started about 4h into the light period. In the ML-LL and LL-ML transitions, leaves were measured for 15 min at the initial irradiance, then light inside the chamber was switched and parameters logged for 45 and 55 min in the ML-LL and LL-ML transition, respectively. In a separate experiment with the same batch of plants, a light saturation curve was performed over a range of irradiance up to 2000 µmol m^-2^ s^-1^, with preincubation at 1000 µmol m^-2^ s^-1^, before transfer to 2000 µmol m^-2^ s^-1^, and measurement at successively lower irradiances for at least 3 min.

### Sampling for metabolite measurements

Light switch experiments were performed with separately grown sets of plants for the ML to LL and the LL to ML using growth irradiance (550 µmol photons m^-2^ s^-1^, ML) and low irradiance (160 µmol photons m^-2^ s^-1^, LL, obtained by placing a polystyrene plate below the light source). For the LL to ML switch, maize plants were grown at 550 µmol photons m^-2^ s^-1^, and then-shaded with a polystyrene plate, starting just before dusk on the day prior to the experiment. LL to ML and ML to LL switches were performed by moving maize plants individually from one light condition to the other light condition and harvesting 5, 10, 15, 30, 60, 120, 300, 600, 1200 or 1800 s after transfer. Control samples (t=0 s; plant not transferred) were also harvested (*n*=4 and 10 for ML to LL and LL to ML switch, respectively). The ML-LL and LL-ML treatments were performed at a two-week interval on separately-grown batches of plants. Samples for a given light switch treatment were collected on a single day. On each day, the first light switch treatment started at about 4 h after light on, and the last was timed such that the treatment and harvest was completed by 6.5 h after light on. This narrow time window was used to minimize sample-to-sample variation due to gradual accumulation of metabolites during the light period. Samples for different time points were randomized across time-of-day. Material was harvested by cutting the leaf (seven cm long section, about 10 cm below the leaf tip) and quenching it in a bath of liquid N_2_ under prevalent irradiance, avoiding shading. After quenching, frozen plant material was stored at −80 °C until further use.

A second ML-LL and LL-ML light switch experiment was performed with separately-grown plants, with less time points (0, 10 s and 20 min, to capture changes immediately after the change in irradiance and at the end of the transition), but more replicates (*n*=10), and samples for one light switch being collected on one day and the other light switch on the following day,

### Metabolite analyses

Frozen samples were homogenised using a ball mill (Retsch, Haan, Germany; https://www.retsch.com) at liquid N_2_ temperature. Metabolites were extracted and quantified by LC-MS/MS and GC-MS as described in Arrivault et al. (2019) and in Lisec et al. (2006), respectively. PEP, pyruvate and 3PGA were determined enzymatically in freshly prepared trichloroacetic acid extracts (Merlo et al. 1993) using a spectrophotometer (Shimadzu, Kyoto, Japan; www.shimadzu.de).

### Statistical analyses

Statistical analysis was performed in Excel and in R Studio Version 1.1.463 (www.rstudio.com) with R version 3.5.1 (https://cran.r-project.org/). Additional information is in the figure legends.

## Data Availability

All data obtained for this study are presented within the Supplementary Datasets.

## Supplementary Material

**Supplementary Figure S1**. Additional gas exchange data.

**Supplementary Figure S2.** Principal Components Analysis: loadings of metabolites.

**Supplementary Figure S3.** Further metabolites and statistical tests in the moderate to low light transition, or the low to moderate light transition.

**Supplementary Figure S4**. Further metabolites and statistical tests in the low to moderate light transition.

**Supplementary Figure S5.** Changes of individual metabolites, metabolite ratios and sets of metabolites in additional light transitions.

**Supplementary Figure S6**. Relationship between *A_n_* and metabolite levels.

**Supplementary Figure S7.** Additional plots of metabolite levels and metabolic traits against *A_n_* in a transition from moderate to low light.

**Supplementary Figure S8.** Additional plots of metabolite levels and metabolic traits against *A_n_* in a transition from low to moderate light

## Supplementary Text

**Supplementary Dataset S1.** Gas exchange data.

**Supplementary Dataset S2.** Metabolite data in the time resolved ML-LL and LL-ML transitions.

**Supplementary Dataset S3**. Metabolite data at 10 s and 1200 s in an independent ML-LL and LL to ML transition.

## Abbreviations

2OG: 2-oxoglutarate
2PG: 2-phosphoglycolate
3PGA: 3-phosphoglycerate
ADPG: ADP- glucose
BSC: bundle sheath
CBC: Calvin Benson Cycle
*A_n_*: CO_2_ assimilation
*Ci*: CO_2_ concentration
CCM: CO_2_- concentrating mechanism
DHAP: dihydroxyacetone phosphate
FBP: fructose 1,6-bisphosphate
FBPase: fructose 1,6-bisphosphatase
Fru2,6BP: fructose 2,6-bisphosphate
F6P: fructose 6-phosphate
G6P: glucose 6- phosphate
MC: mesophyll cell
NADP-MDH: NADP-malate dehydrogenase
NAD-ME: NAD-dependent malic enzyme
NADP-ME: NADP-dependent malic enzyme
PEPCK: phospho*enol*pyruvate carboxykinase
PC: principal component
PEP: phospho*enol*pyruvate
PEPC: phospho*eno*lpyruvate carboxylase
PPDK: phosphate dikinase
PRK: phosphoribulokinase
R5P: ribose 5-phosphate
Ru5P: ribulose 5-phosphate
Ru5P: ribulose-1,5- bisphosphate
Rubisco: ribulose-1,5-bisphosphate carboxylase/oxygenase
SBP: sedoheptulose 1,7- bisphosphate,sedoheptulose 1,7-bisphosphatase
S7P: sedoheptulose 7-phosphate
SBPase: fructose-1,6- seduheptulosebisphosphatase
*g*: stomatal conductance
triose-P: triose phosphate
UDPG: UDPG-glucose
Xu5P: xylulose 5-phosphate.

## Acknowledgments

We would like to thank Hirofumi Ishihara and Eva-Theresa Pyl and for their help during the harvests. This work was financially supported by the Max Planck Society (MG), the German Federal Ministry of Education and Research (BMBF grant 031B0205B to SA, DBM, ARF, MS), the UK Biological and Biotechnological Scientific Research Council (via grant BB/T007583/1 awarded to JK) and UK Research and Innovation - Future Leaders Fellowships scheme (via grant MR/T042737/1 awarded to JK).

## Author contributions

SA, DBM, ARF and MS conceived and planned the experiments. SA grew and harvested the plants. SA, DBM and MG extracted the samples for downstream analyses. MG and SA processed and analyzed LC-MS/MS and enzymatically assayed samples. DBM processed and analyzed GC-MS samples. CS and JK performed gas exchange measurements using dynamic equations. DBM performed initial gas exchange measurements and analysis. SA and DBM performed statistical analyses. SA, DBM, ARF and MS wrote the manuscript. ARF and MS supervised the project. All authors approved the final version.

## Conflict of interest

The Authors declare that there is no conflict of interest.

## Notes

### Competing Interest Statement

The authors have declared no competing interest.

## References

Arce Cubas, L., Vath, R.L., Bernardo, E.L., Sales, C.R.G., Burnett, A.C., Kromdijk, J. (2023a) Activation of CO_2_ assimilation during photosynthetic induction is slower in C_4_ than in C_3_ photosynthesis in three phylogenetically controlled experiments. Front Plant Sci 2023:13:1091115. 10.3389/fpls.2022.1091115

Arce Cubas, L., Sales, C.R.G., Vath, R.L., Bernardo, E.L., Burnett, A.C., Kromdijk, J. (2023b) Lessons from relatives: C_4_ photosynthesis enhances CO_2_ assimilation during the low-light phase of fluctuations. Plant Physiol 193, 1073–1090. 10.1093/plphys/kiad355

Arrivault, S., Guenther, M., Ivakov, A., Feil, R., Vosloh, D., van Dongen, J.T., Sulpice, R. and Stitt, M. (2009) Use of reverse-phase liquid chromatography, linked to tandem mass spectrometry, to profile the Calvin cycle and other metabolic intermediates in Arabidopsis rosettes at different carbon dioxide concentrations. Plant J. 59, 824–839. 10.1111/j.1365-313X.2009.03902.x

Arrivault, S., Obata, T., Szecowka, M., Mengin, V., Guenther, M., Hoehne, M., Fernie, A.R. and Stitt, M. (2017) Metabolite pools and carbon flow during C_4_ photosynthesis in maize: ^13^CO_2_ labeling kinetics and cell-type fractionation. J Exp Bot 68, 283–298. 10.1093/jxb/erw414

Arrivault, S., Moraes, T.A., Obata, T., Medeiros, D.B., Fernie, A.R., Boulouis, A., Ludwig, M., Lunn, J.E., Borghi, G.L., Schlereth, A., Guenther, M. and Stitt, M. (2019) Metabolite Profiles reveal interspecific variation in operation of the Calvin-Benson cycle in both C_4_ and C_3_ plants. J Exp Bot 6870, 1843–1858. 10.1093/jxb/erz051

Ashton, A.R., Hatch, M.D. (1983). Regulation of C4 photosynthesis: regulation of activation and inactivation of NADP-malate dehydrogenase by NADP and NADPH. Arch Biochem Biophys. 227, 416–24. doi: 10.1016/0003-9861(83)90471-x.

Ashton, A.R., Burnell, J.N., Furbank, R.T., Jenkins, C.L.D., Hatch, M.D. (1990) Enzymes of C4 photosynthesis. In: Lea PJ, ed. Methods in plant biochemistry, Volume 3, Enzymes of primary metabolism. London, 39-72.

Bellasio, C., Griffiths, H. (2014a) The operation of two decarboxylases, transamination, and partitioning of C_4_ metabolic processes between mesophyll and bundle sheath cells allows light capture to be balanced for the maize C_4_ pathway. Plant Physiol 164, 466–480. 10.1104/pp.113.228221

Bellasio, C., Griffiths, H. (2014b). Acclimation to low light by C_4_ maize: implications for bundle sheath leakiness. Plant Cell Environ 37, 1046–1058. 10.1111/pce.12194

Bellasio, C., Beerling, D.J., Griffiths, H. (2016) Deriving C_4_ photosynthetic parameters from combined gas exchange and chlorophyll fluorescence using an Excel tool: theory and practice. Plant Cell Environ 39, 1164–1179. 10.1111/pce.12626

Betti, M., Bauwe, H., Busch, F.A., Fernie, A.R., Keech, O., Levey, M., Ort, D.R., Parry, M.A., Sage, R., Timm, S., Walker, B., and Weber, A.P. (2016) Manipulating photorespiration to increase plant productivity: recent advances and perspectives for crop improvement. J Exp Bot 67: 2977–2988. 10.1093/jxb/erw076

Bovdilova, A., Alexandre, B.M., Hoppner A., Matias Luıs, I., Alvarez, C.E., Bickel, D., Gohlke, H., Decker, C., Nagel-Steger L., Alseekh, S., Fernie, A.R., Drincovich, M.F., Abreu, I.A., Maurino, V.G. (2019) Posttranslational modification of the NADP-malic enzyme involved in C_4_ photosynthesis fine-tunes the enzymatic activity during the day. Plant Cell 31, 2525–2539. doi: 10.1093/plcell/koab262

Bräutigam, A., Schliesky, S., Külahoglu, C., Osboune, C.P., Weber, A.P.M. (2014) Towards an integrative model of C_4_ photosynthetic subtypes: insights from comparative transcriptome analysis of NAD-ME, NADP-ME, and PEP-CK C_4_ species. J Exp Bot 65, 3579-2593. 10.1093/jxb/eru100

Bräutigam, A., Schlüter, U., Lundgren, M.R., Flachbart, S., Ebenhöh, O., Schönknecht, G., Christin, P.A., Bleuler, S., Droz, J.M., Osborne, C.P., Weber, A.P.M., Gowik, U. (2018) Biochemical mechanisms driving rapid fluxes in C_4_ photosynthesis. bioRxiv preprint doi.org/10.1101/387431.

Borghi, G.-L., Arrivault, S., Günther, M., Medeiros, D. B., Dell’Aversana, E., Fusco, G.M., Carillo, P. Ludwig. M., Fernie, A.R., Lunn, J.E. and Stitt, M. (2022) Metabolic Profiles in C_3_, C_3_-C_4_ intermediate, C_4_-like and C_4_ species in the genus *Flaveria*. J Exp Bot 73, 1581-1681 10.1093/jxb/erab540

Carmo-Silva, A.E., Powers, S.J., Keys, A.J., Arrabaça, M.C., Parry, M.A. (2008) Photorespiration in C_4_ grasses remains slow under drought conditions. Plant Cell Environ. 31, 925–940. 10.1111/j.1365-3040.2008.01805.x.

Carmo-Silva, A.E., Salvucci, M.E. (2013) The regulatory properties of Rubisco activase differ among species and affect photosynthetic induction during light transitions. Plant Physiol 161, 1645–1655. 10.1104/pp.112.213348

Chen, Y.B., Lu, T.C., Wang, H.-X., Shen, J., Bu, T.-T., Chao, Q., Gao, Z.-F., Zhu, X.-G., Wang, Y.-F., Wang, B.-C. (2014) Posttranslational Modification of Maize Chloroplast Pyruvate Orthophosphate Dikinase Reveals the Precise Regulatory Mechanism of Its Enzymatic Activity. Plant Physiol 165, 534–549. 10.1104/pp.113.231993

Christin P.-A., Besnard G., Samaritani E., Duvall M.R., Hodkinson T.R., Savolainen V. & Salamin N. (2008) Oligocene CO_2_ decline promoted C_4_ photosynthesis in grasses. Curr Biol 18, 37–43. 10.1016/j.cub.2007.11.058

de Veau, E.J., Burris, J.E. (1989) Photorespiratory rates in wheat and maize as determined by o-labeling. Plant Physiol 90, 500–511. 10.1104/pp.90.2.500

Dietz, K.J., Heber, U. (1984) Rate-limiting factors in leaf photosynthesis. I. Carbon fluxes in the Calvin cycle, Biochim. Biophys. Acta 767, 432–443, 10.1016/0005-2728(84)90041-0

Deans, R.M., Farquhar, G.D., Busch, F.A. (2019) Estimating stomatal and biochemical limitations during photosynthetic induction. Plant Cell Environ 42, 3227–3240.

Doncaster, H.D., Leegood, R.C. (1987) Regulation of Phosphoenolpyruvate Carboxylase Activity in Maize Leaves. Plant Physiol 84, 82–87. 10.1104/pp.84.1.82

Doncaster, H.D., Adcock, M.D., Leegood, R.C. (1989) Regulation of photosynthesis in leaves of C_4_ plants following a transition from high to low light. BBA 973, 176–184, 10.1016/S0005-2728(89)80419-0

Edwards, G.E. & Walker, D. (1983) C_3_, C_4_: Mechanisms, Cellular and Environmental Regulation of Photosynthesis. University of California Press, Berkeley, CA, USA. ISBN-10: 0520050185.

Edwards, E.J., Osborne, C.P., Stromberg, C.A.E., Smith, S.A., Bond, W.J., Christin, P.A., et al. (2010) C_4_ grasses consortium. The origins of C_4_ grasslands: integrating evolutionary and ecosystem science. Science 328, 587-591. 10.1126/science.117721

Faske, M., Holtgrefe, S., Ocheretina, O., Meiste, M., Backhausen, J.E., Scheibe, R. (1995) Redox equilibria between the regulatory thiols of light/dark-modulated chloroplast enzymes and dithiothreitol: fine-tuning by metabolites. BBA 1247, 135–142. 10.1016/0167-4838(94)00203-s

Foyer, C.H., Bloom, A.J., Queva, l.G., Noctor, G. (2009) Photorespiratory metabolism: genes, mutants, energetics, and redox signaling. Annu Rev Plant Biol 60, 455–484. 0.1146/annurev.arplant.043008.091948.

Fu, X., Gregory, L.M., Weise, S.E., Walker, B.J. (2023) Integrated flux and pool size analysis in plant central metabolism reveals unique roles of glycine and serine during photorespiration. Nature Plants 9, 169–178. 10.1038/s41477-022-01294-9

Farquhar, G. (1983) On the Nature of Carbon Isotope Discrimination in C_4_ Species. Funct Plant Biol 10, 205–226. 10.1071/PP9830205

Furbank, R.T., Jenkins, C.L.D., Hatch, M.D. (1990) C_4_ photosynthesis: quantum requirement, C_4_ and overcycling and Q-cycle Involvement. Aust J Plant Physiol 17, 1-7. 10.1071/PP9900001

Furbank, R.T. (2011). Evolution of the C_4_ photosynthetic mechanism: are there really three C_4_ acid decarboxylation types? J Exp Bot 62, 3103–3108. 10.1093/jxb/err080

Furbank, R.T., Foyer, C., Stitt, M. (1985) The localization of sucrose synthesis in maize leaves. Planta 164, 172–178. 10.1007/BF00396079

Furbank, R.T., Leegood, R.C. (1984) Carbon metabolism and gas exchange in leaves of *Zea mays* L.: Interaction between the C_3_ and C_4_ pathways during photosynthetic induction. Planta 162, 457-462, 10.1007/BF00393459

Gerhardt, R., Stitt, M., Heldt, H.W. (1987) Subcellular metabolite levels in spinach leaves. Regulation of sucrose synthesis during diurnal alterations in photosynthesis. Plant Physiol 83, 399–407. 10.1104/pp.83.2.399

Hatch, M.D., Agostino, A., Jenkins, C. (1995) Measurement of the leakage of CO_2_ from bundle-sheath cells of leaves during C_4_ photosynthesis. Plant Physiol 108, 173–181. 10.1104/pp.108.1.173

Hatch, M.D., Osmond, C.B. (1976) Compartmentation and transport in C4 photosynthesis. In Encyclopedia of Plant Physiology, S. C.R. and H. U., eds (Springer-Verlag. Berlin), pp. 144–184.

Hatch, M.D. (2002) C_4_ photosynthesis: Discovery and resolution. Photosynth Res 73: 251–256. 10.1007/1-4020-3324-9_78

Hammond, E.T., Andrews, T.J., Mott, K.A., Woodrow, I.E. (1998) Regulation of Rubisco activation in antisense plants of tobacco containing reduced levels of Rubisco activase. Plant J 1 14, 101–110. 10.1046/j.1365-313X.1998.00103.x

Heldt, H.W., Piechulla, B. (2021) Plant biochemistry. 5^th^ Edition, Cambridge, MA: Academic Press.

Herzog, B., Stitt, M., Heldt, H.W. (1984) Control of photosynthetic sucrose synthesis by fructose 2,6-bisphosphate. III. Properties of the cytosolic fructose-1,6-bisphosphatase. Plant Physiol 75, 561-565. 10.1104/pp.75.3.554

Kaiser, E., Morales, A., Harbinson, J., Heuvelink, E., Prinzenberg, A.E., Marcelis, L.F.M. (2016) Metabolic and diffusional limitations of photosynthesis in fluctuating irradiance in *Arabidopsis thaliana*. Scientific Reports 6, 31252. 10.1038/srep31252.

Knuesting, J., Scheibe, R. (2018) Small Molecules Govern Thiol Redox Switches. Trends in Plant Sci. 23, 769–782. 10.1016/j.tplants.2018.06.007

Krall, J. P., Pearcy, R. W. (1993) Concurrent measurements of oxygen and carbon dioxide exchange during light flecks in maize (*Zea mays* L.). Plant Physiol. 103, 823–828. 10.1104/pp.103.3.823

Kromdijk, J., Ubierna, N., Cousins, A.B., and Griffiths, H. (2014) Bundle-sheath leakiness in C_4_ photosynthesis: a careful balancing act between CO_2_ concentration and assimilation. J Exp Bot 65, 3443–3457. 10.1093/jxb/eru157

Kromdijk, J., Głowacka, K., Leonelli, L., Gabilly, S.T., Iwai, M., Niyogi, K.K., Long, S.P. (2016) Improving photosynthesis and crop productivity by accelerating recovery from photoprotection. Science 354, 857–861. 10.1126/science.aai887

Kubasek, J., Urban, O., Santrucek, J. (2013) C_4_ plants use fluctuating light less efficiently than do C_3_ plants: a study of growth, photosynthesis and carbon isotope discrimination. Physiol. Plant. 149, 528–539. 10.1111/ppl.12057

Laetsch, W.M. (1974) The C_4_ syndrome: a structural analysis. Ann Rev Plant Physiol 25, 27–52. 10.1146/annurev.pp.25.060174.000331

Laing, W.A., Stitt, M., Heldt, H.W. (1981) Changes in the activity of ribulosephosphate kinase and fructose-and sedoheptulose-bisphosphatase in chloroplasts. BBA 637, 348–359. 10.1016/0005-2728(81)90174-2

Laisk, A., Edwards, G.E. (1998) Oxygen and electron flow in C_4_ photosynthesis: Mehler reaction, photorespiration and CO_2_ concentration in the bundle sheath. Plantae 20, 632–645 10.1007/s004250050366

Lawson, T., Kramer, D.M., Raines, C.A. (2012) Improving yield by exploiting mechanisms underlying natural variation of photosynthesis. COBIOT 23, 215–220. 10.1016/j.copbio.2011.12.012

Lee, M. S., Boyd, R. A., Ort, D. R. (2022) The photosynthetic response of C_3_ and C_4_ bioenergy grass species to fluctuating light. GCB Bioenergy 14, 37–53. doi:10.1111/gcbb.12899

Leegood, R.C. (1985) The intercellular compartmentation of metabolites in leaves of *Zea mays L*. Planta 164, 163–171. 10.1007/BF00396078

Leegood, R., and Furbank, R. (1984) Carbon metabolism and gas exchange in leaves of *Zea mays L*. Planta 162, 450–456. 10.1007/BF00393459

Leegood, R.C., and von Caemmerer, S. (1988) The relationship between contents of photosynthetic metabolites and the rate of photosynthetic carbon assimilation in leaves of *Amaranthus edulis L*. Planta 174, 253–262. 10.1007/BF00394779

Leegood, R., and von Caemmerer, S. (1989) Some relationships between contents of photosynthetic intermediates and the rate of photosynthetic carbon assimilation in leaves of *Zea mays L*. Planta 178, 258–266. 10.1007/BF00393202

Li, Y.T., Luo, J., Liu, P., Zhang, Z.-S. et al. (2021) C_4_ species utilize fluctuating light less efficiently than C_3_ species. Plant Physiol 187, 1288–1291. 10.1093/plphys/kiab411

Lisec, J., Schauer, N., Kopka, J., Willmitzer, L., Fernie, A.R. (2006) Gas chromatography mass spectrometry-based metabolite profiling in plants. Nat Protoc 1, 387–396. 10.1038/nprot.2006.59

Lorimer, G.H. (1981) The carboxylation and oxygenation of ribulose 1,5-bisphosphate: the primary events in photosynthesis and photorespiration. Annu Rev Plant Physiol 32, 349–382. doi: 10.1146/annurev.pp.32.060181.002025

Mallmann, J., Heckmann, D., Bräutigam, A., Lercher, M.J., Weber, A.P., Westhoff P., Gowik, U. (2014) The role of photorespiration during the evolution of C_4_ photosynthesis in the genus Flaveria. eLIFE 3:e02478. 10.7554/eLife.02478

Medeiros, D.B., Ishihara, H., Guenther, M., Rosado de Souza, L., Fernie, A.R., Stitt, M. and Arrivault, S. (2022) ^13^CO_2_ labelling kinetics in maize reveal impaired efficiency of C_4_ photosynthesis in low irradiance. Plant Physiol 190, 280–304. doi.org/10.1093/plphys/kiac306

Merlo, L., Geigenberger, P., Hajirezaei, M., Stitt M. (1993) Changes of carbohydrates, metabolites and enzyme activities in potato tubers during development, and within a single tuber along a stolon–apex gradient. Plant Physiol 142, 392–402. 10.1016/S0176-1617(11)81243-5.

Michelet, L., Zaffagnini, M., Morisse, S., Sparla, F., Pérez-Pérez, M.E., Francia, F., Danon, A., Marchand, C.H., Fermani, S., Trost, P., Lemaire, S.D. (2013) Redox regulation of the Calvin-Benson cycle: something old, something new. Front Plant Sci 4, 470. 10.3389/fpls.2013.00470

Munekage, Y.N. (2016) Light harvesting and chloroplast electron transport in NADP-malic enzyme type C_4_ plants. Curr Opin Plant Biol 31, 9–15. 10.1016/j.pbi.2016.03.001

Mott, K.A., Snyder, G.W., Woodrow, I.E. (1997) Kinetics of Rubisco activation as determined from gas-exchange measurements in antisense plants of *Arabidopsis thaliana* containing reduced levels of Rubisco activase. Aust J Plant Physiol 24, 811–818. 10.1071/PP97071

Mott, K.A., Woodrow, I.E. (2000) Modelling the role of Rubisco activase in limiting non-steady-state photosynthesis. J Exp Bot 51, 399–406. 10.1093/jexbot/51.suppl_1.399

Om, K., Arias, N.N., Jambor C.C., MacGregor A., Rezacheck, A.N., Harfud, C., Kunz, H.-H., Wang, Z., Huang, P., Zhang, Q., Rosnow, J., Brutnell T.P., Cousins, A.B., Chastain, C.J. (2022) Pyruvate, phosphate dikinase regulatory protein impacts light response of C_4_ photosynthesis in *Setaria viridis*. Plant Physiol 190, 1117–1133. 10.1093/plphys/kiac333

Onishi, J.-I., Kanai, R. (1983) Differentiation of photorespiratory activity between mesophyll and bundle sheath cells of C_4_ plants I. Glycine oxidation by mitochondria. PCP 24, 1411–1420. 10.1093/oxfordjournals.pcp.a076662

Osmond, C. (1981) Photorespiration and photoinhibition: some implications for the energetics of photosynthesis. BBA 639, 77–98. 10.1016/0304-4173(81)90006-9.

Osmond, C.B., and Harris, B. (1971) Photorespiration during C_4_ photosynthesis. BBA 234, 270–282. 10.1016/0005-2728(71)90082-X.

Pearcy, R.W. (1990) Sunflecks and photosynthesis in plant canopies. “Annu Rev Plant Physiol 41, 421–453. 10.1146/annurev.pp.41.060190.002225

Pearcy, R.W., Muraoka, H., Valladares, F. (2005) Crown architecture in sun and shade environments: assessing function and trade-offs with a three-dimensional simulation model. New Phytol 166, 791–800. 10.1111/j.1469-8137.2005.01328.x

Pengelly, J.J.L., Sirault, X.R.R., Tazoe, Y, Evans, J.R, Furbank, RT, von Caemmerer, S. (2010) Growth of the C_4_ dicot *Flaveria bidentis:* photosynthetic acclimation to low light through shifts in leaf anatomy and biochemistry. J Exp Bot 61, 4109–4122. 10.1093/jxb/erq226

Pignon, C., Jaiswal, D., McGrath, J.M. (2017) Loss of photosynthetic efficiency in the shade. An Achilles heel for the dense modern stands of our most productive C_4_ crops? J Exp Bot 68, 335-345. 10.1093/jxb/erw456

Prinsley, R.T., Dietz, K.-J., Leegood, RC (1986a) Regulation of photosynthetic carbon assimilation in spinach leaves after a decrease in irradiance. The influence of a decrease in irradiance on photosynthetic carbon assimilation in leaves of Spinacia oleracea L. BBA Bioenergetics, 849, 254–263. 0.1016/0005-2728(86)90032-0

Prinsley, R.T., Hunt, S., Smith, A.M., Leegood R.C. (1986b). The influence of a decrease in irradiance on photosynthetic carbon assimilation in leaves of *Spinacia oleracea L*. Planta 167, 414–420. 10.1007/BF00391348

Portis, A.R. Jr., Li, C., Wang, D., Salvucci, M.E. (2008) Regulation of Rubisco activase and its interaction with Rubisco. J Exp Bot 59, 1597–1604. doi: 10.1093/jxb/erm240

Rebeille, F., Hatch, M.D. (1986) Regulation of NADP-malate dehydrogenase in C4 plants: relationship among enzyme activity, NADPH to NADP ratios, and thioredoxin redox states in intact maize mesophyll chloroplasts. Arch Biochem Biophys. 249, 171–9. doi: 10.1016/0003-9861(86)90572-2

Roeske, C.A., Chollet, R. (1989) Role of Metabolites in the Reversible Light Activation of Pyruvate,Orthophosphate Dikinase in *Zea mays* Mesophyll Cells in Vivo. Plant Physiol 90, 330–337. 10.1104/pp.90.1.330

Saathoff, A.J., Welles, J. (2021) Gas exchange measurements in the unsteady state. Plant Cell Environ 44, 3509–3523. doi: 10.1111/pce.14178.

Sage, R. F., McKown, A. D. (2006). Is C_4_ photosynthesis less phenotypically plastic than C_3_ photosynthesis? J Exp Bot 57, 303–317. doi:10.1093/jxb/erj040

Sage, R.F., Christin, P.A., Edwards, E.J. (2011) The C_4_ plant lineages of planet Earth. J Exp Bot 62, 3155–3169. 10.1093/jxb/err048

Sage, R.F, Seemann R, Sharkey TD. (1987) The time course for deactivation and reactivation of ribulose-I,5-bisphosphate carboxylase following changes in CO_2_ and O_2_. In Progress in Photosynthesis Research, ed. 1. Biggins, 3: 285-88, Dordrecht: Martinus-Nijhoff

Sales, C.R.G, Ribeiro, R.V., Hayashi, A.H., Marchiori, P.E.R., Silva, K.I., Martin, M.O. Silveira, J.A.G., Silveira, N.M., Machado, E.C. (2018) Flexibility of C_4_ decarboxylation and photosynthetic plasticity in sugarcane plants under shading, Environ Exp Botany 149, 34–42. doi.org/10.1016/j.envexpbot.2023.105351

Sales, R.G.C, Ribiero, R.F., Marchiori P.E.R., Kromdijk, J., Machado, E.V. (2023) The negative impact of shade on photosynthetic efficiency in sugarcane may reflect a metabolic bottleneck. Environ Exp Botany 105631. doi.org/10.1016/j.envexpbot.2023.105351

Scheibe, R. (1991) Redox-modulation of chloroplast enzymes: a common principle for individual control. Plant Physiol 96, 1–3. 10.1104/pp.96.1.1

Schlüter, U., Weber, A.P.M. (2020) Regulation and Evolution of C_4_ photosynthesis. Annu Rev Plant Biol 71, 183–215. 10.1146/annurev-arplant-042916-040915

Slattery RA, Walker BJ, Weber APM, Ort DR. (2018) The impacts of fluctuating light on crop performance. Plant Physiol 176, 990–1003-doi: 10.1104/pp.17.01234

Stitt, M. (1990) Fructose 2,6-bisphosphate as a regulatory metabolite in plants. Annu Rev Plant Physiol. 41, 153–185. 10.1146/annurev.pp.41.060190.001101

Stitt, M., Wirtz, W., Heldt, H.W. (1980) Metabolite levels during induction in the chloroplast and extrachloroplast compartments of spinach protoplasts. Biochim Biophy Acta 593, 85–102. 10.1016/0005-2728(80)90010-9

Stitt, M., Wirtz, W., Heldt, H.W. (1983) Regulation of sucrose synthesis by cytoplasmic fructose bisphosphatase and sucrose phosphate synthetase during photosynthesis in varying light and carbon dioxide. Plant Physiol 72, 767–774. 10.1104/pp.72.3.767

Stitt. M., Heldt, H.W. (1985a) Generation and maintenance of concentration gradients between the mesophyll and bundle sheath in maize leaves. Biochim Biophys Acta 808, 400–414. 10.1016/0005-2728(85)90148-3

Stitt, M., Heldt, H.W. (1985b) Control of photosynthetic sucrose synthesis by fructose 2,6-bisphosphate. IV. Intercellular metabolite distribution and properties of the cytosolic fructose 1,6-bisphosphatase in maize leaves. Planta 164, 179-188. 10.1007/BF00396080

Stitt, M., Scheibe, R., Feil, R. (1989) Response of photosynthetic electron transport to a sudden decrease of irradiance in the saturating or the limiting range. Biochim Biophys Acta 241–249. 10.1016/S0005-2728(89)80428-1

Stitt, M., Lunn, J., Usadel, B. (2010) Arabidopsis and primary photosynthetic metabolism—more than the icing on the cake. TPJ 61, 1067–1091. 10.1111/j.1365-313X.2010.04142.x

Stitt, M., and Zhu, X.G. (2014) The large pools of metabolites involved in intercellular metabolite shuttles in C4 photosynthesis provide enormous flexibility and robustness in a fluctuating light environment. Plant Cell Environ 37, 1985–1988. 10.1111/pce.12290

Stitt, M., Borghi, G.L., Arrivault, S. (2021) Targeted Metabolite Profiling as a Top-Down Approach to Uncover Inter-Species Diversity and Identify Key Conserved Operational Features in the Calvin Benson cycle. J Exp Bot. 10.1093/jxb/erab291

Tanaka, Y., Adachi, S., Yamori, W. (2019) Natural genetic variation of the photosynthetic induction response to fluctuating light environment. Curr Opin Plant Biol 49, 52–59. 0.1016/j.pbi.2019.04.010

Taylor, S.H., Long, S.P. (2017) Slow induction of photosynthesis on shade to sun transitions in wheat may cost at least 21% of productivity. Philosophical Transactions of the Royal Society B: Biological Sciences 372, 20160543. 10.1098/rstb.2016.0543.

Tazoe, Y., Hanba, Y.T., Furumoto, T., Noguchi, K., Terashima, I. (2008) Relationships between quantum yield for CO_2_ assimilation, activity of key enzymes and CO_2_ leakiness in *Amaranthus cruentus*, a C_4_ dicot, grown in high or low light. Plant Cell Physiol 49, 19–29. 10.1093/pcp/pcm160

Tcherkez, G.G., Farquhar, G.D., and Andrews, T.J. (2006). Despite slow catalysis and confused substrate specificity, all ribulose bisphosphate carboxylases may be nearly perfectly optimized. PNAS 103, 72467251. 10.1073/pnas.0600605103

Tirumala Devi, M., Raghavendra, A.S. (1992) Light Activation of Phosphoenolpyruvate Carboxylase in Maize Mesophyll Protoplasts. J Plant Physiol 139, 431–435. 10.1016/S0176-1617(11)80490-6

Townsend, A.J., Retkute, R., Chinnathambi, K., Randall, J.W.P., Foulkes, J., Silva, E.M., Murchie, E.H. (2018) Suboptimal Acclimation of Photosynthesis to Light in Wheat Canopies. Plant Physiol 176, 1233–1246. 10.1104/pp.17.01213

Ubierna, N., Sun, W., Kramer, D.M., Cousins, A.B. (2013) The efficiency of C_4_ photosynthesis under low light conditions in *Zea mays, Miscanthus × giganteus* and *Flaveria bidentis*. Plant Cell Environ 36, 365–381. 10.1111/j.1365-3040.2012.02579.x

Ueno, O., Yoshimura, Y., Sentoku, N. (2005) Variation in the activity of some enzymes of photorespiratory metabolism in C_4_ grasses. Ann Bot 96, 863–869. 10.1093/aob/mci238

Usuda, H. (1985) Changes in Levels of Intermediates of the C_4_ Cycle and Reductive Pentose Phosphate Pathway during Induction of Photosynthesis in Maize Leaves. Plant Physiol 78, 859–864. 10.1104/pp.78.4.859

Usuda, H. (1987) Changes in Levels of Intermediates of the C_4_ Cycle and Reductive Pentose Phosphate Pathway under Various Light Intensities in Maize Leaves. Plant Physiol 84, 549–554. 10.1104/pp.84.2.549

Vialet-Chabrand, S., Matthews, J.S.A., Simkin, A.J., Raines, C.A., Lawson, T. (2017) Importance of Fluctuations in Light on Plant Photosynthetic Acclimation. Plant Physiol 173, 2163–2179. 10.1104/pp.16.01767

Vidal, J., Bakrim, N., Hodges, M. (2002) The Regulation of Plant Phosphoenolpyruvate Carboxylase by Reversible Phosphorylation. In Foyer CH, Noctor G (eds), Photosynthetic Nitrogen Assimilation and Associated Carbon and Respiratory Metabolism, pp. 135-150. ©Kluwer Academic Publishers. The Netherlands

Volk, R.J., Jackson, W.A. (1972) Photorespiratory phenomena in maize: oxygen uptake, isotope discrimination, and carbon dioxide efflux. Plant Physiol 49, 218–223. 10.1104/pp.49.2.218

von Caemmerer, S. (2000) Biochemical models of leaf photosynthesis. CSIRO Publishing, Collingwood, Australia

von Caemmerer, S., Furbank, R. (2003) The C_4_ pathway: an efficient CO_2_ pump. Photosynth Res 77 191–207. 10.1023/A:1025830019591

Walker, R.P., Acheson, R.M., R. Técsi, L.I, Leegood, R.C. (1997) Phosphoenolpyruvate carboxykinase in C_4_ plants: its role and regulation. Aust J Plant Physiol 24: 459–468). DOI:10.1071/PP97007

Walker RP, Chen ZH, Acheson RM, Leegod RC (2002) Effects of Phosphorylation on Phospho*enol*pyruvate Carboxykinase from the C_4_ Plant Guinea Grass. Plant Physiol 128, 165–172 10.1104/pp.010432

Wingler, A., Walker, R.P., Chen, Z.-H., Leegood, R.C. (1999) Phosphoenolpyruvate Carboxykinase Is Involved in the Decarboxylation of Aspartate in the Bundle Sheath of Maize. Plant Physiol 120, 539–546. DOI: 10.1104/pp.120.2.539

Walker, B.J., VanLoocke, A., Bernacchi, C.J., Ort, D.R. (2016) The Costs of Photorespiration to Food Production Now and in the Future. Annu Rev Plant Biol 67, 107–29. 10.1146/annurev-arplant-043015-111709

Wang, Y., Long, S.P., Zhu, X.-G. (2014a) Elements required for an efficient NADP-malic enzyme type C_4_ photosynthesis. Plant Physiol 164, 2231–2246. 10.1104/pp.113.230284

Wang, Y., Brautigam, A., Weber, A.P., Zhu, X.G. (2014b) Three distinct biochemical subtypes of C_4_ photosynthesis? A modelling analysis. J Exp Bot 65, 3567–3578. 10.1093/jxb/eru058

Wang, Y., Chan, K.Z., Long, S.P. (2021) Towards a dynamic photosynthesis model to guide yield improvement in C_4_ crops. TPJ 107, 343–359. 10.1111/tpj.15408

Wang, Y., Stutz, S. S., Bernacchi, C. J., Boyd, R. A., Ort, D. R., Long, S. P. (2022). Increased bundle-sheath leakiness of CO_2_ during photosynthetic induction shows a lack of coordination between the C_4_ and C_3_ cycles. New Phytol 236, 1661–1675. doi: 10.1111/nph.18485

Weber, A.P., and von Caemmerer, S. (2010) Plastid transport and metabolism of C_3_ and C_4_ plants--comparative analysis and possible biotechnological exploitation. Curr Opin Plant Biol 13, 257–265. 10.1016/j.pbi.2010.01.007

Weissmann, S., Ma, F., Furuyama, K., Gierse, J., Berg, H., Shao, Y., Taniguchi, M., Allen, D.K, Brutnell, T.P. (2016) Interactions of C_4_ subtype metabolic activities and transport in maize are revealed through the characterization of dct2 mutants. The Plant Cell 28, 466–484. 10.1105/tpc.15.00497

Wirtz, W., Stitt, M. and Heldt, H.W. (1982) Light activation of Calvin cycle enzymes as measured in pea leaves. FEBS Letts. 142, 223–226. 10.1016/0014-5793(82)80139-7

Woo, K.C., Anderson, J., Boardman, K., Downton. J, Osmond, C.B., Thorne, S.W. (1970) Deficient photosystem 2 in agranal bundle sheath chloroplasts of C_4_ plants. PNAS 67, 18–25. 10.1073/pnas.67.1.18

Woodrow, I.E., Berry, A. (1988) Enzymatic regulation of photosynthetic CO_2_ fixation in C_3_ plants, Ann. Rev Plant Physiol Plant Mol Biol 39, 533–559. 10.1146/annurev.pp.39.060188.002533

Woodrow, I.E., Mott, K.A. (1989) Rate limitation of non-steady-state photosynthesis by ribulose-1,5-bisphosphate carboxylase in spinach. Funct Plant Biol 16, 487–500. 10.1071/PP9890487

Woodrow, I.E., Walker, D.A. (1980) Light-mediated activation of stromal sedoheptulose bisphosphatase. Biochem J 191, 845–949. 10.1042/bj1910845

Woodrow, I.E., Murphy, D.J., Walker, D.A. (1983) Regulation of photosynthetic carbon metabolism. The effect of inorganic phosphate on stromal sedoheptulose 1,7-bisphosphatase. Eur J Biochem 32, 121-23. 10.1111/j.1432-1033.1983.tb07335.x

Woodrow, I.E., Furbank, R.T., Brooks, A., Murphy, D.I. (1985) The requirements for steady state in the C_3_ reductive pentose phosphate pathway of photosynthesis. BBA 807, 23–71. 10.1016/0005-2728(85)90257-9

Woodrow, I.E., Kelly, M.E., Mott, K.A. (1996) Limitation of the rate of ribulose bisphosphate carboxylase activation by carbamylation and the ribulose bisphosphate carboxylase activase activity; development and test of a mechanistic model. Aust J Plant Physiol 23,141–14. 10.1071/PP9960141

Wu, A., Hammer, G.L., Doherty, A., von Caemmerer, S., Farquhar, G.D. (2019) Quantifying impacts of enhancing photosynthesis on crop yield. Nat Plants 5, 380–388. 10.1038/s41477-019-0398-8

Yamori, W., Masumoto, C., Fukayama, H., Makino, A. (2012) Rubisco activase is a key regulator of non-steady-state photosynthesis at any leaf temperature and, to a lesser extent, of steady-state photosynthesis at high temperature. TPJ 71, 871–880. 10.1111/j.1365-313X.2012.05041.x

Qiao, M. Y., Zhang, Y. J., Liu, L. A., Shi, L., Ma, Q. H., Chow, W. S., & Jiang, C. D. (2020) Do rapid photosynthetic responses protect maize leaves against photoinhibition under fluctuating light? Photosynthesis Research 149, 57–68. 10.1007/s11120-020-00780-5

