## Supplemental Figures and Supplemental Text for "Metabolite profiling reveals slow and uncoordinated adjustment of C_4_ photosynthesis to sudden changes in irradiance"

NEW RESULTS

### SUPPLEMENTAL FIGURES AND TEXT

#### Supplemental Figures (pages 2-20)

**Page 2: Supplementary Figure S1**. Additional gas exchange data.

**Page 6: Supplementary Figure S2.** Principal Components Analysis: loadings of metabolites.

**Page 8: Supplementary Figure S3.** Further metabolites and statistical tests in the moderate to low light transition, or the low to moderate light transition.

**Page 9: Supplementary Figure S4**. Further metabolites and statistical tests in the low to moderate light transition.

**Page 10: Supplementary Figure S5.** Changes of individual metabolites, metabolite ratios and sets of metabolites in additional light transitions.

**Page 14: Supplementary Figure S6**. Relationship between $A_n$ and metabolite levels.

**Page 17 Supplementary Figure S7.** Additional plots of metabolite levels and metabolic traits against $A_n$ in a transition from moderate to low light.

**Page 19: Supplementary Figure S8.** Additional plots of metabolite levels and metabolic traits against $A_n$ in a transition from low to moderate light

#### Supplemental text (page 21 onwards, sections listed on side 21)

**Supplementary Figure S1. Additional gas exchange data.** These figures are Supplementary to Fig. 1 and data are provided in Supplementary Dataset S1A and S1B.
(**A**) Light saturation response (mean ± SD, *n*=5 leaves on separate plants).

(**B**) Time linear plot of $A_n$ for the ML-LL transition (mean ± SD, *n*=5).

(**C**) Curve fitting of the decrease of $A_n$ during the ML-LL transition. The plot shows mean values (*n*=5) and the curve fitting is blue.

(**D**) Time linear and log scaled plots of *g* for the ML-LL transition (mean ± SD, *n*=5). The time point in white corresponds to time 0, before the light transition.

(**E**) Time linear and log scaled plots of *Ci* for the ML-LL transition (mean ± SD, *n*=5). The time point in white corresponds to time 0, before the light transition.

(**F**) Time linear plot of $A_n$ for the LL-ML transition (mean ± SD, *n*=6). (

(**G**) Curve fittings of the increases of $A_n$ during the LL-ML transition. The plots show mean values (*n*=6) and the curve fittings are blue.

(**H**) Time linear and log scaled plots of *g* for the LL-ML transition (mean ± SD, *n*=6). The time point in white corresponds to time 0, before the light transition.

(**I**) Time linear and log scaled plots of *Ci* for the LL-ML transition (mean ± SD, *n*= 6). The time point in white corresponds to time 0, before the light transition.

(**J**) Comparison of *An* data from Figure 1 corrected by dynamic equation ("corrected") and uncorrected ("non-corrected") for the M-LL and LL-ML transitions (mean ± SD, *n*=5). The time point in white corresponds to time 0, before the light transition. The plots are presented with time on a log scale.

(**K**) Comparison of two independent experiments obtained for the ML-LL transition and for the LL-ML transition; "non-corrected" are data from the experiment shown in Figure 1, where data was recorded every second but in this plot (like panel J) are not corrected by the dynamic equation ((i.e., as in panel J), "non-corrected 2" are data from an independent experiment in which *An* was measured at 6 s intervals, and correction was not possible as each time point was a recorded data average. This second set of measurement was performed on the batch of plants that were sampled for metabolite analyses. For a better comparison between the experiments, data are shown as a percentage of *An* at time 0. The plots show ± SD (*n*=5 for non-corrected ML-LL, *n*=6 for non-corrected LL-ML, *n*=3 for non-corrected 2 ML-LL and *n*=5 for non-corrected 2 LL-ML). The time points in white correspond to time 0, before the light transition.

(**L**) Expansion of the ML-LL response from 60 s onwards to show the slight recovery of $A_n$ in the independent uncorrected experiment (data recorded every 6 seconds, not corrected by dynamic equation). The plot shows mean ± SD (*n*=3). The relatively high SD is due to differences in $A_n$ between the three independent replicates (see Supplemental Dataset S1B). Significance was analyzed using a paired t-test, to focus on the comparison between different time points, and separate this from differences in $A_n$ between replicates The increase was significant ($p = 0.03$, $n = 3$, paired t-test comparing for each replicate average $A_n$ between 267-300 s with average $A_n$ between 1776-1800 s). The x-axis corresponds to time on a linear scale.

(**M**) Each individual leaf measurement of corrected $A_n$ shown separately for replicate LL-ML transition (Figure 1B shows the average and SD of these values). The plot is presented with time on a log scale.

**Supplementary Figure S1. Continued.**

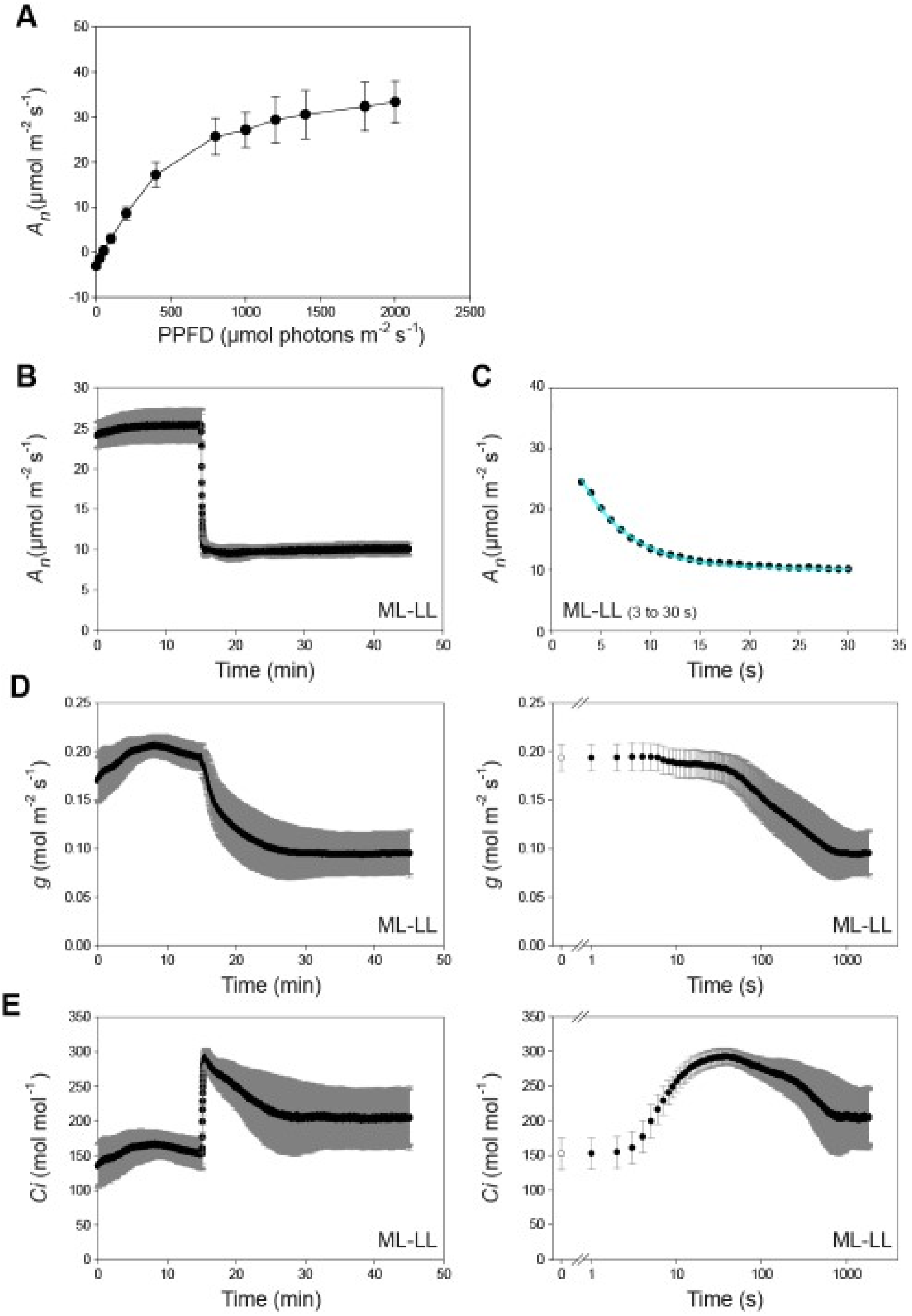

**Supplementary Figure S1. Continued.**

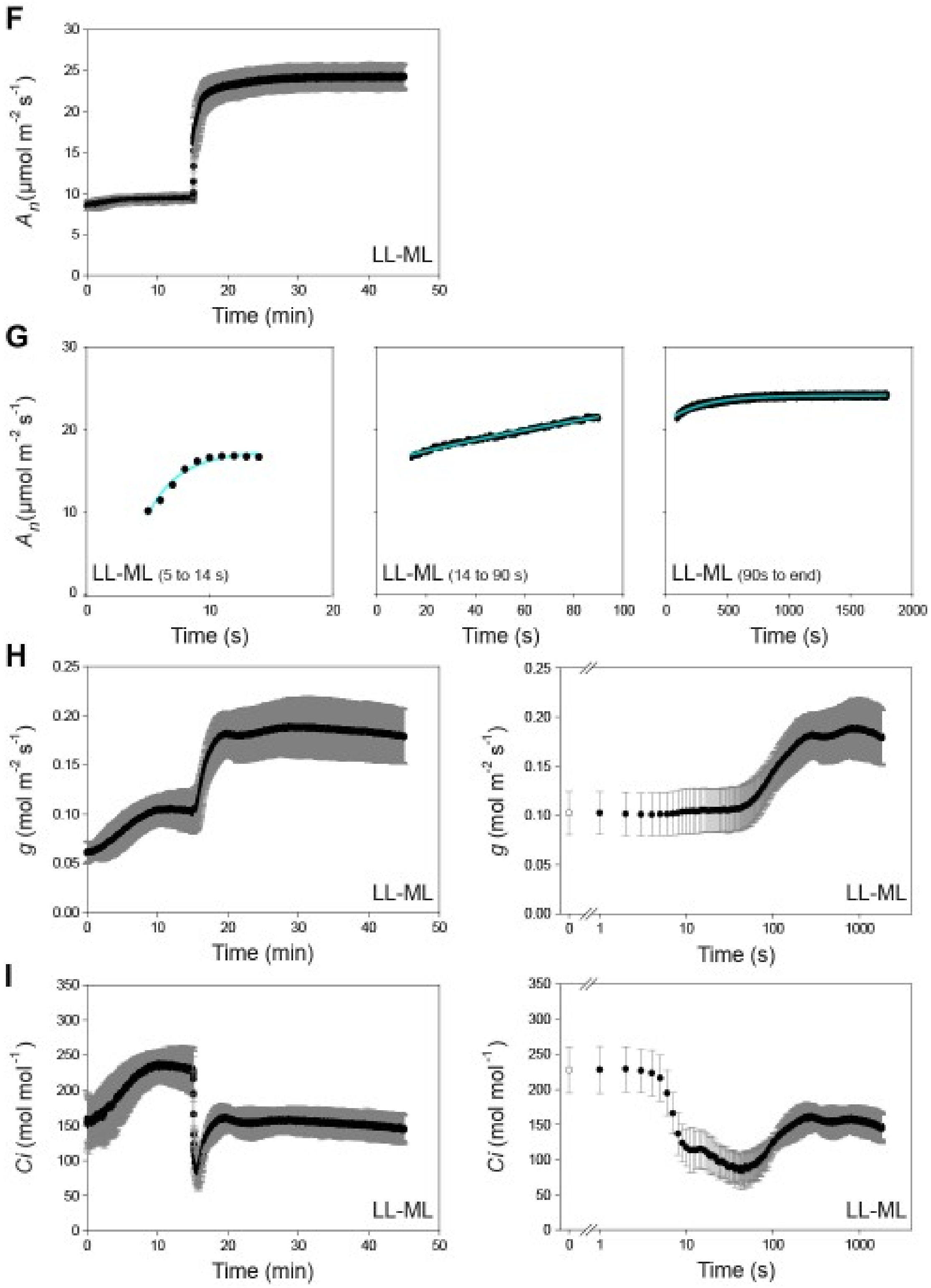

**Supplementary Figure S1. Continued.**

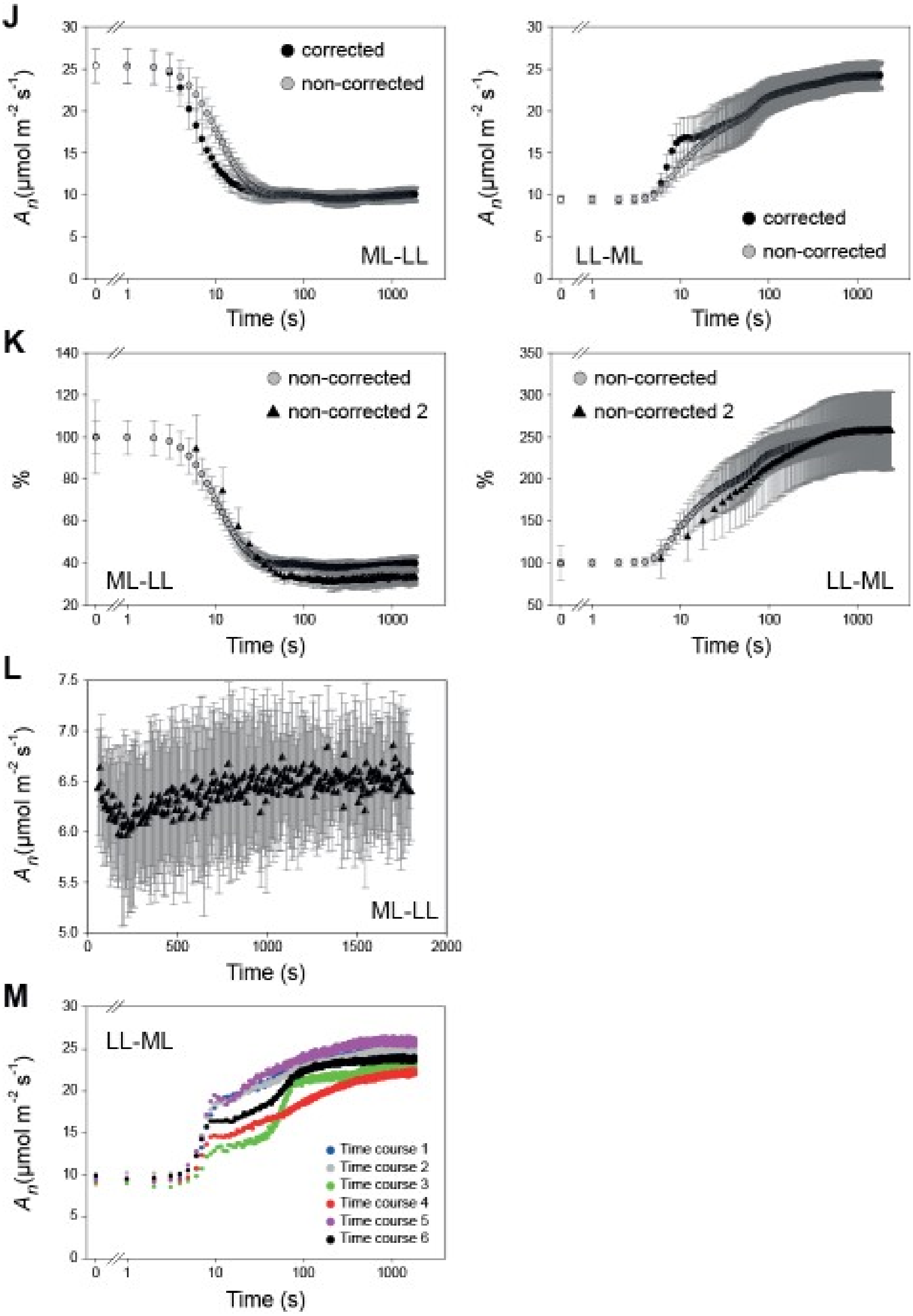

**Supplementary Figure S2. Principal Components Analysis: loadings of metabolites.** (**A**) ML-LL transition and (**B**) LL-ML transition . Means for each time point were used to perform the analyses (*n*=4 to 5 for ML-LL, and *n*=4, except for time 0 s where *n*=10 for LL-ML). In (**A**) and (**B**), both the PC analysis (as in Fig. 2) and the variables are included. $CO_2$ assimilation ($A_n$) is highlighted in green and metabolites from the CBC and CCM are highlighted in dark and light grey, respectively. The time points in the PCA are identified by color (see insert) and the arrows show the time sequence in the transition. This figure is supplementary to Fig. 2.

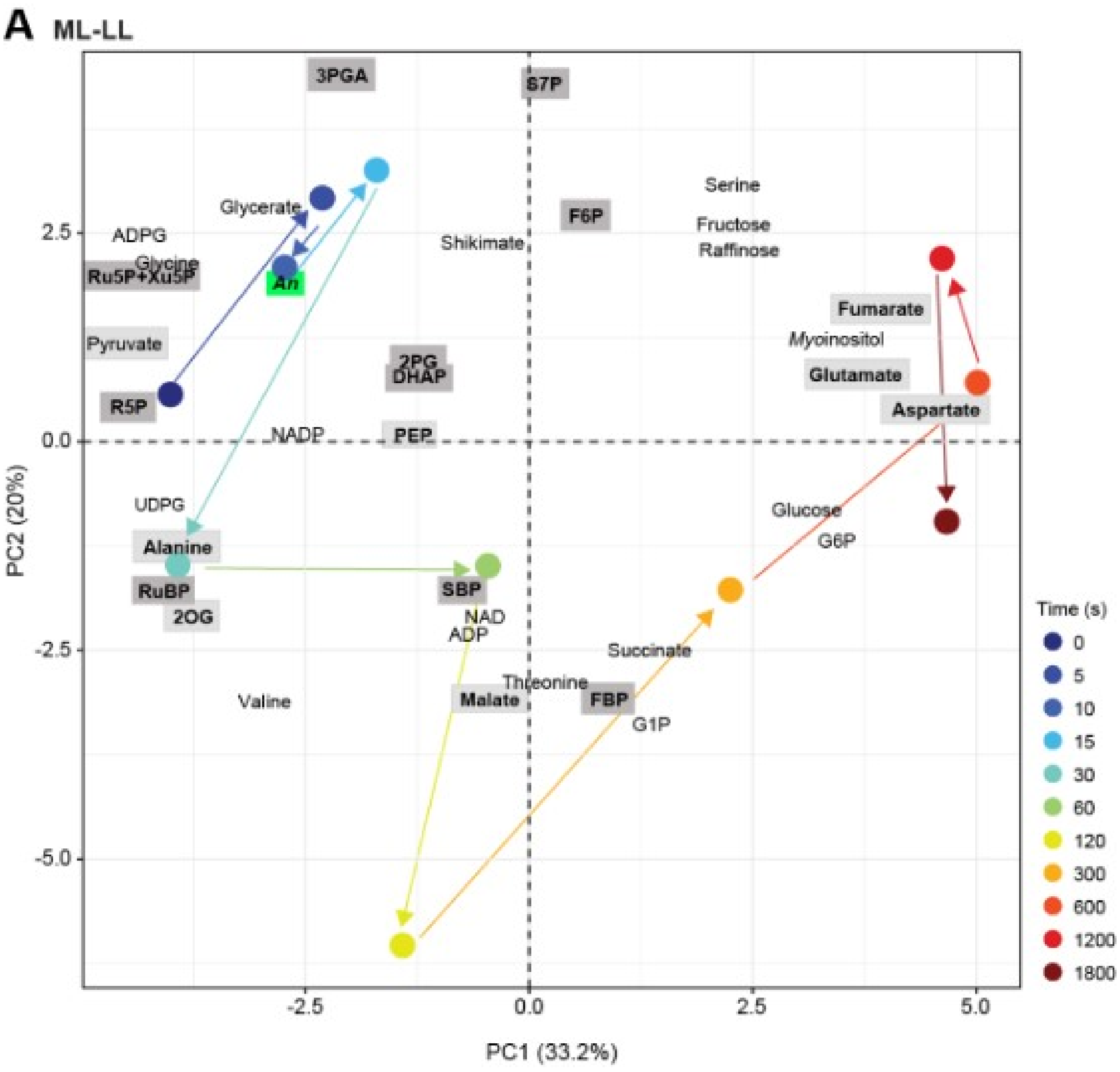

**Supplementary Figure S2. Continued.**

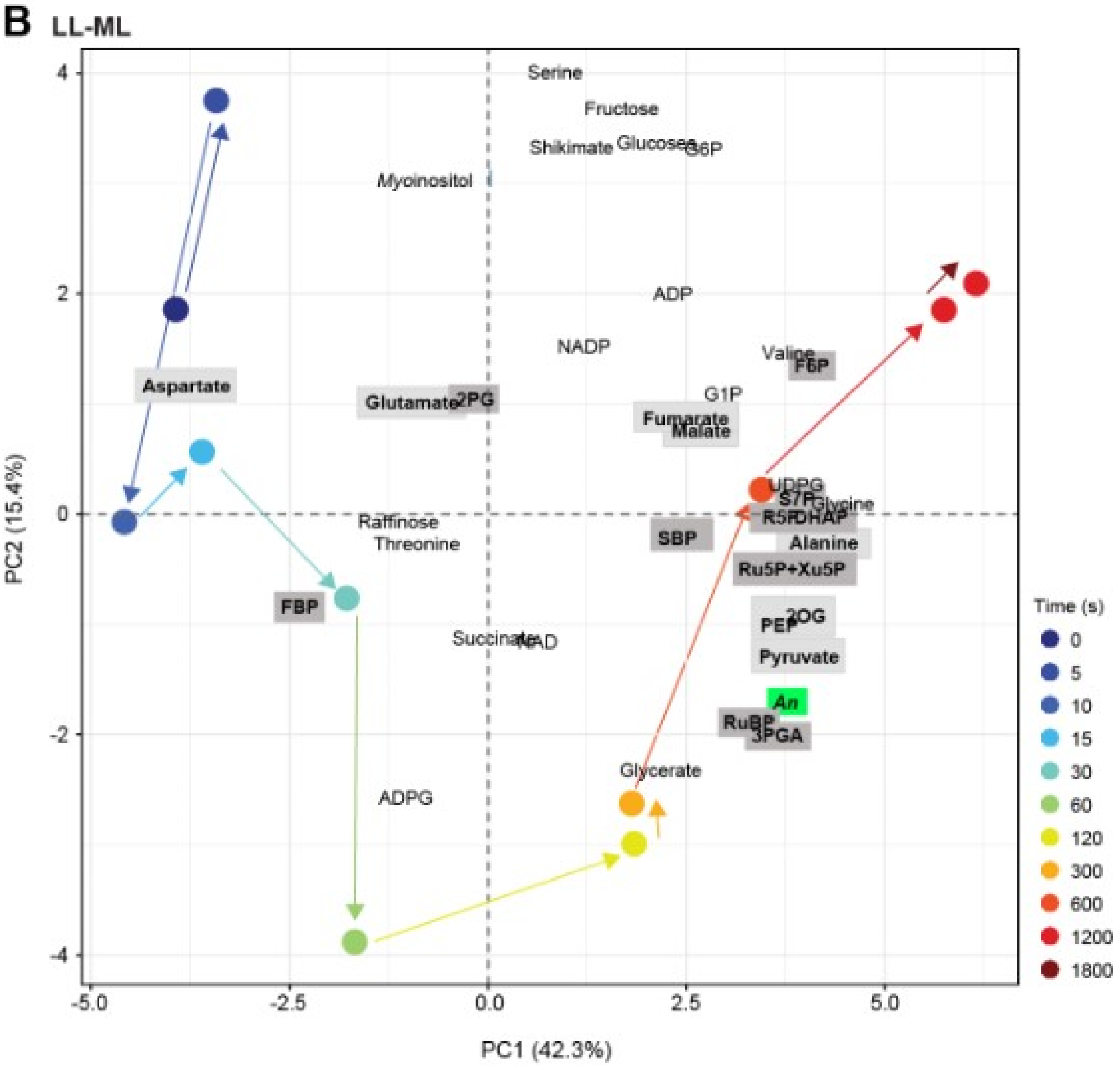

**Supplementary Figure S3. Further metabolites and statistical tests in the moderate to low light transition, or the low to moderate light transition.**

(**A**) Moderate light to low light transition. Amounts of metabolites (nmol $g^{-1}$ FW) are shown as mean ± SD (*n*=4 to 5). Significant differences (T-test) to time zero (i.e., ML) are indicated by stars (* $p < 0.05$; ** $p < 0.01$: *** $p < 0.001$). Time is shown on a log scale. For original data, see Supplementary Dataset S2.

(**B**) Visualization of significant changes. Two approaches were taken. In one approach (left hand side) the entire time sequence was analyzed. T-tests were performed individually between time zero (i.e., ML) and each time in the time sequence. In the second approach (right hand side) the response was divided into time segments to test for changes in different phases of the transient that might be masked in the complete transient. Time segments were defined based on the PCA of the combined $CO_2$ assimilation response and metabolome response (Fig. 2). Time segments are separated by solid vertical lines, and the grey column indicates first time in a given time segment. T-tests were performed between the first time in the time segment and each later time in the time segment. The results are visualized as stars (* $p < 0.05$; ** $p < 0.01$: *** $p < 0.001$; blue denotes an increase and red a decrease (see legend). The time segments are also indicated by colored blocks in **A**.

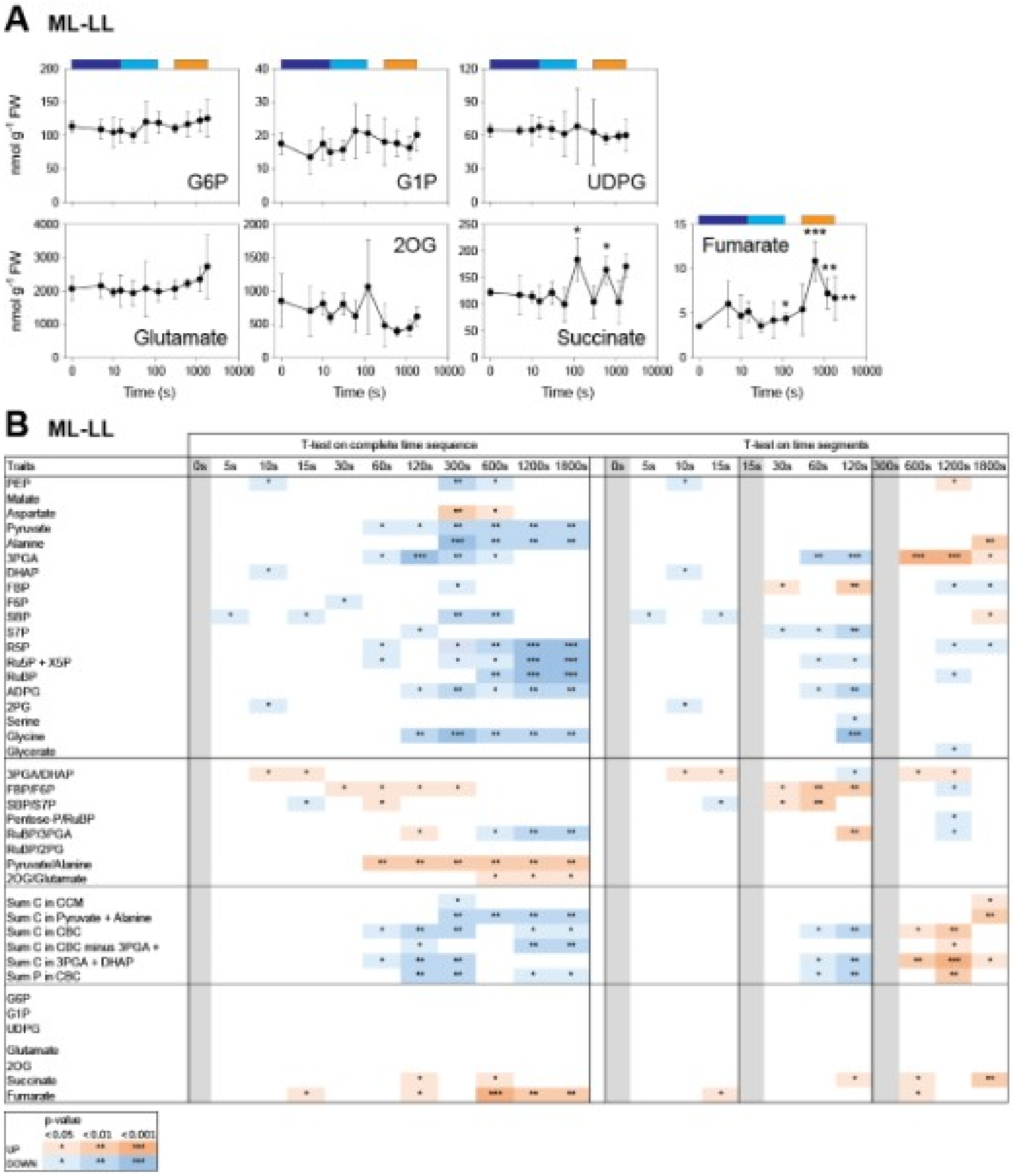

**Supplementary Figure S4. Further metabolites and statistical tests in the low to moderate light transition.**
(**A**) Low light to moderate light transition. Amounts of metabolites (nmol $g^{-1}$ FW) are shown as mean ± SD (*n*=4 except for time zero where *n*=10). Significant differences (T-test) to time zero (i.e., LL) are indicated by stars (* p < 0.05; ** p < 0.01: *** p < 0.001). Time is shown on a log scale. For original data, see Supplementary Dataset S2.
(**B**) Visualization of significant changes. Two approaches were taken. In one approach (left hand side) the entire time sequence was analyzed. T-tests were performed individually between time zero (i.e., ML in the ML-LL transition and LL in the LL-ML transition) and each time in the time sequence. In the second approach (right hand side) the response was divided into time segments to test for changes in different phases of the transient that might be masked in the complete transient. Time segments were defined based on the PCA of the combined $CO_2$ assimilation response and metabolome response (Fig. 2). Time segments are separated by solid vertical lines, and the grey column indicates first time in a given time segment. T-tests were performed between the first time in the time segment and each later time in the time segment. The results are visualized as stars (* p < 0.05; ** p < 0.01: *** p < 0.001; blue denotes an increase and red a decrease (see legend). The time segments are also indicated by colored blocks in **A**.

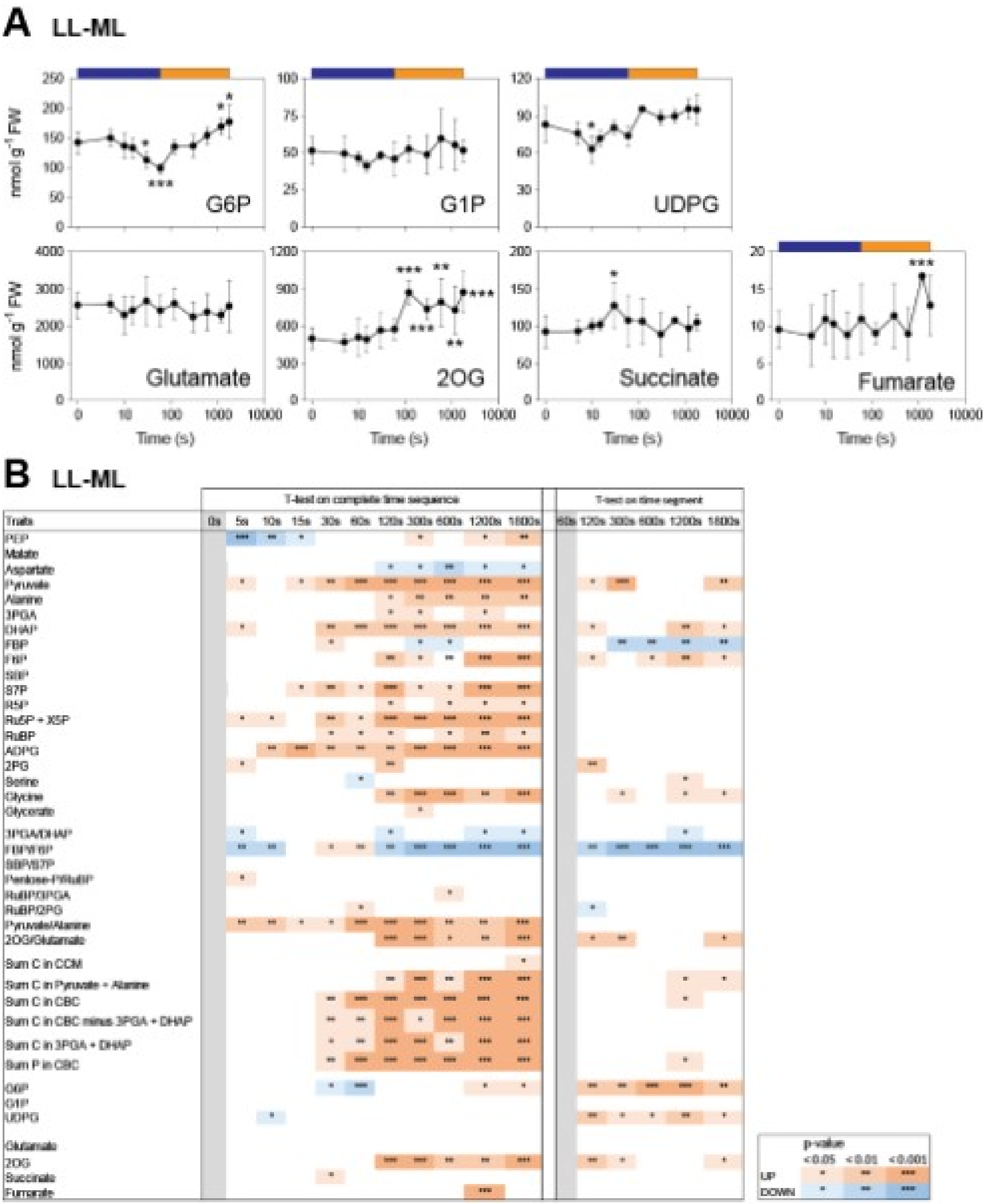

**Supplementary Figure S5. Changes of individual metabolites, metabolite ratios and sets of metabolites in additional light transitions**.

**(A-B)** Levels of metabolites at 10 s and 1200 s,

**(A)** ML-LL transition,

(**B**) ML-LL transition (plot overleaf)**.**

Data shown are means ± SD (*n*=10 for ML-LL, n= 9 to 10 for LL-ML). Statistically significant differences from time zero were determined by t-test (* $p < 0.05$; ** $p < 0.01$: *** $p < 0.001$). All x-axes correspond to time on a log scale. The original data are provided in Supplementary Dataset S3.

**(C-D)** Comparison of changes of individual metabolites, metabolite ratios and sets of metabolites at 10 s and 1200 s in the two experiments (plots overleaf, two and three pages later)

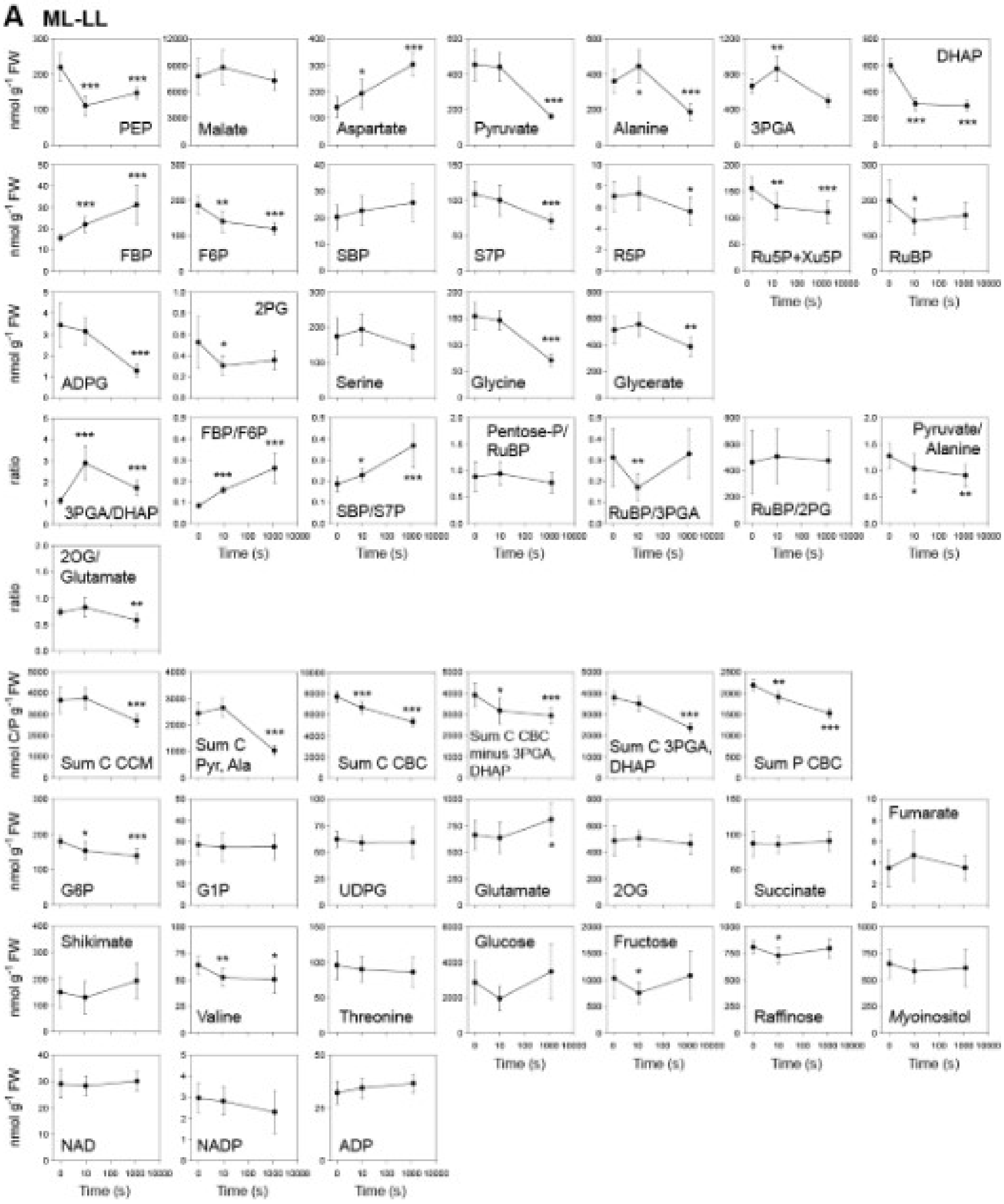

**Supplementary Figure S5. Continued.**

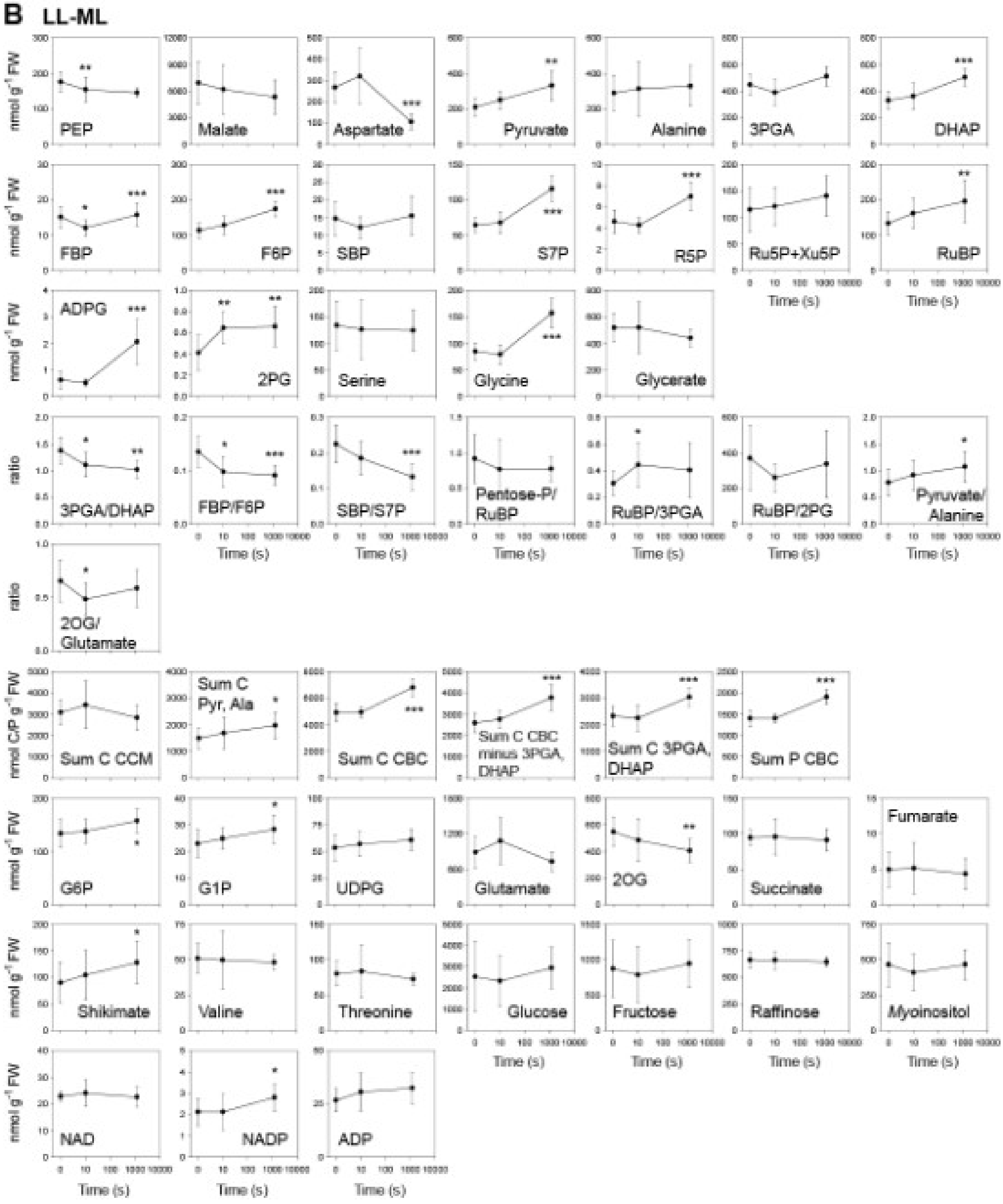

**Supplementary Figure S5. Continued.**

**(C-D)** Comparison of changes of individual metabolites, metabolite ratios and sets of metabolites at 10 s and 1200 s in the two experiments. The plots show the ratios at 10 s and 20 min to the average amounts at time 0 s (before light transition). Time 10 s is presented in light grey while time 20 min is presented in dark grey. The values obtained in the repetition with two time points are indicated with hatching.
**(C)** ML-LL transition
**(D)** LL-ML transition (plot overleaf).

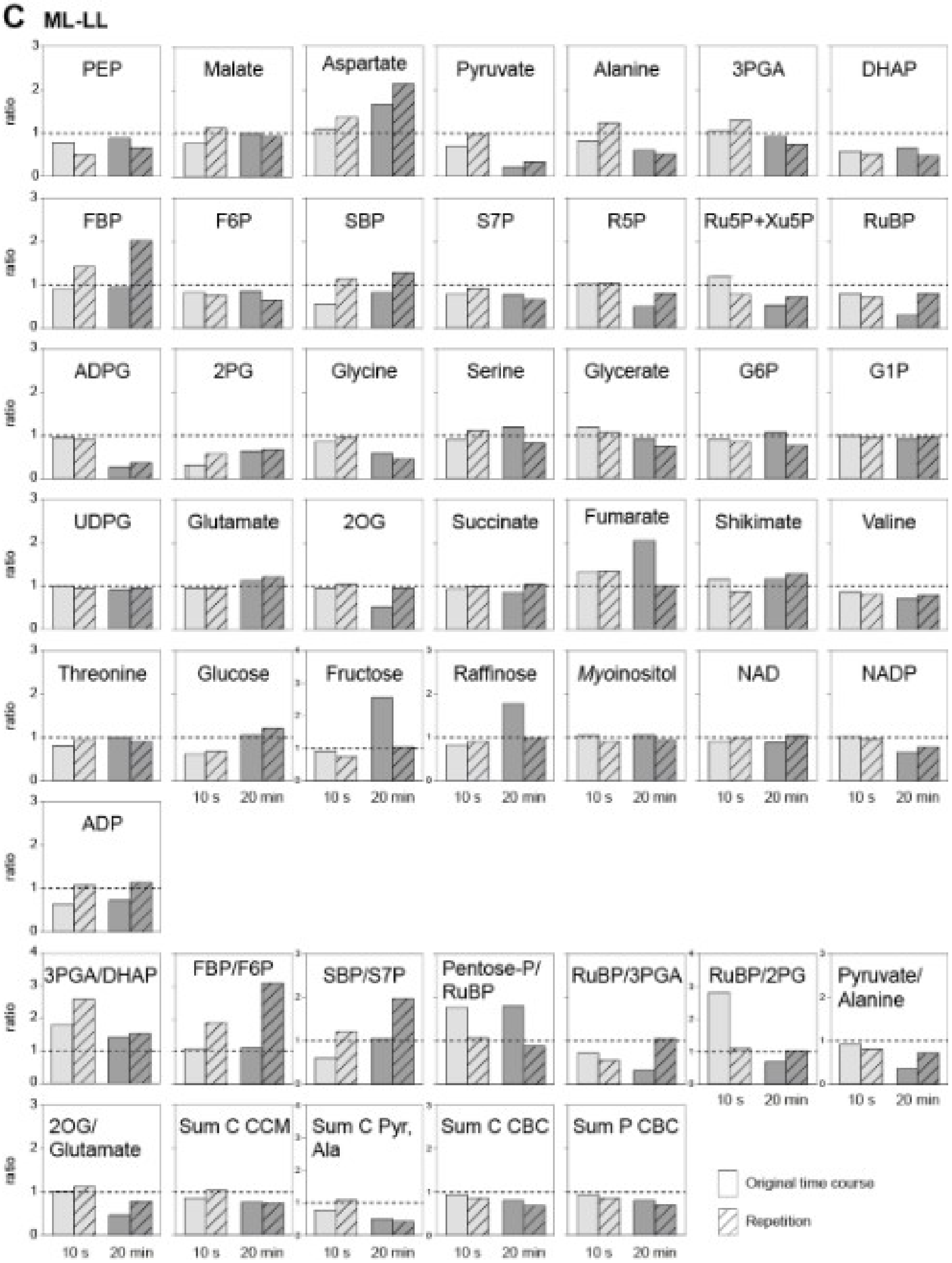

**Supplementary Figure S5. Continued.**

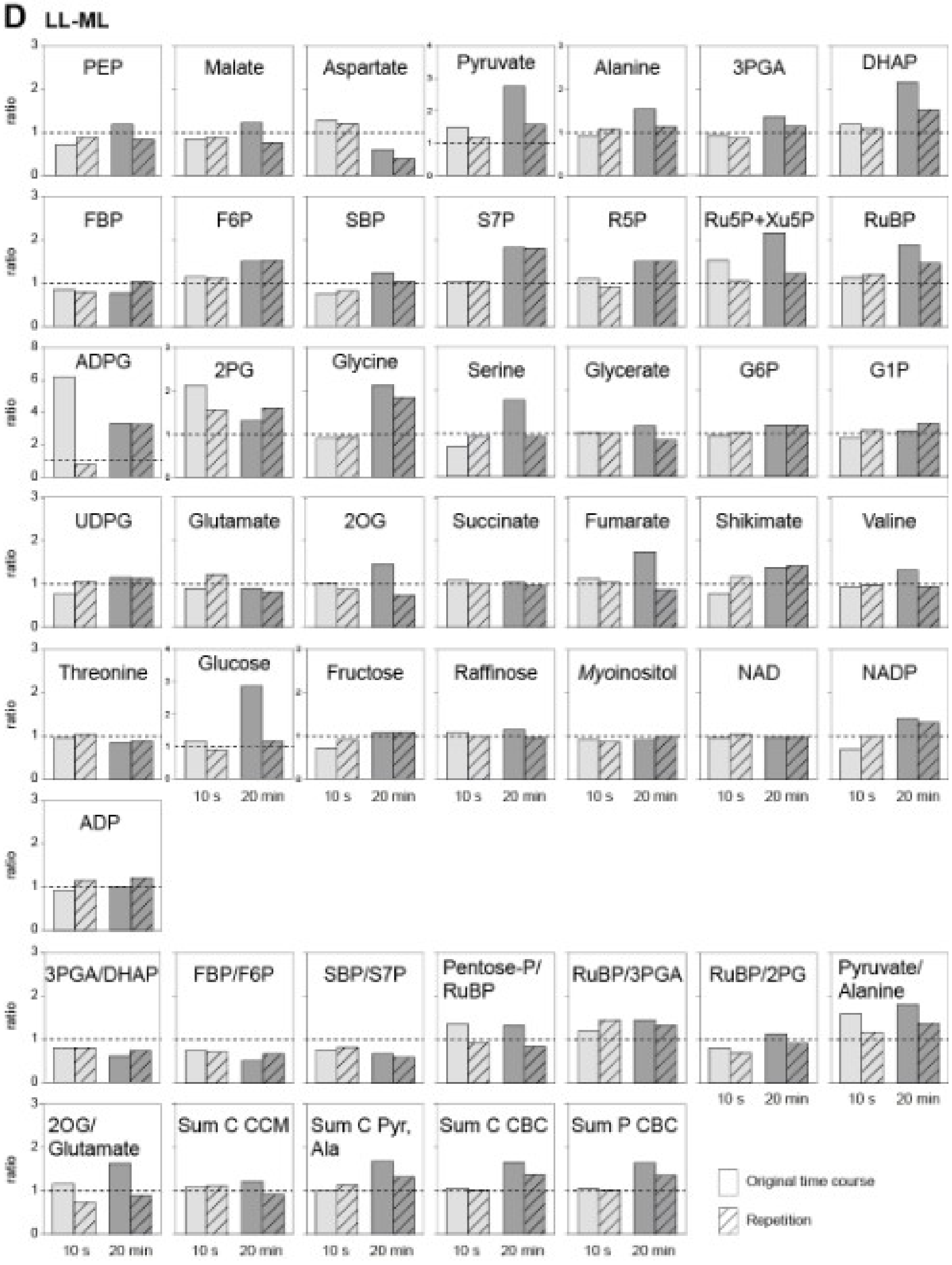

**Supplementary Figure S6. Relationship between $A_n$ and metabolite levels.**
Heatmaps showing Pearson's correlation coefficients between $A_n$ and metabolites in the (**A**) ML-LL transition and (**B**) LL-ML transition (plot overleaf). Pearson's correlation coefficients were calculated using individual samples at all time points after the transition to LL or the transition to ML (time zero excluded). Significant correlations ($p < 0.05$) are coloured red (positive) and blue (negative), while correlations that were not significant are shown in grey. Self-correlations are identified in dark grey. The hierarchical clustering is shown. Panels A and B are expanded versions of Fig. 5A.
**(C)** $R^2$ and $p$ values for a regression between $A_n$ and metabolite levels, ratios or summed pool and **(D)** plots of summed C in the CCM vs CBC (plots overleaf, two pages later).

**A** ML-LL

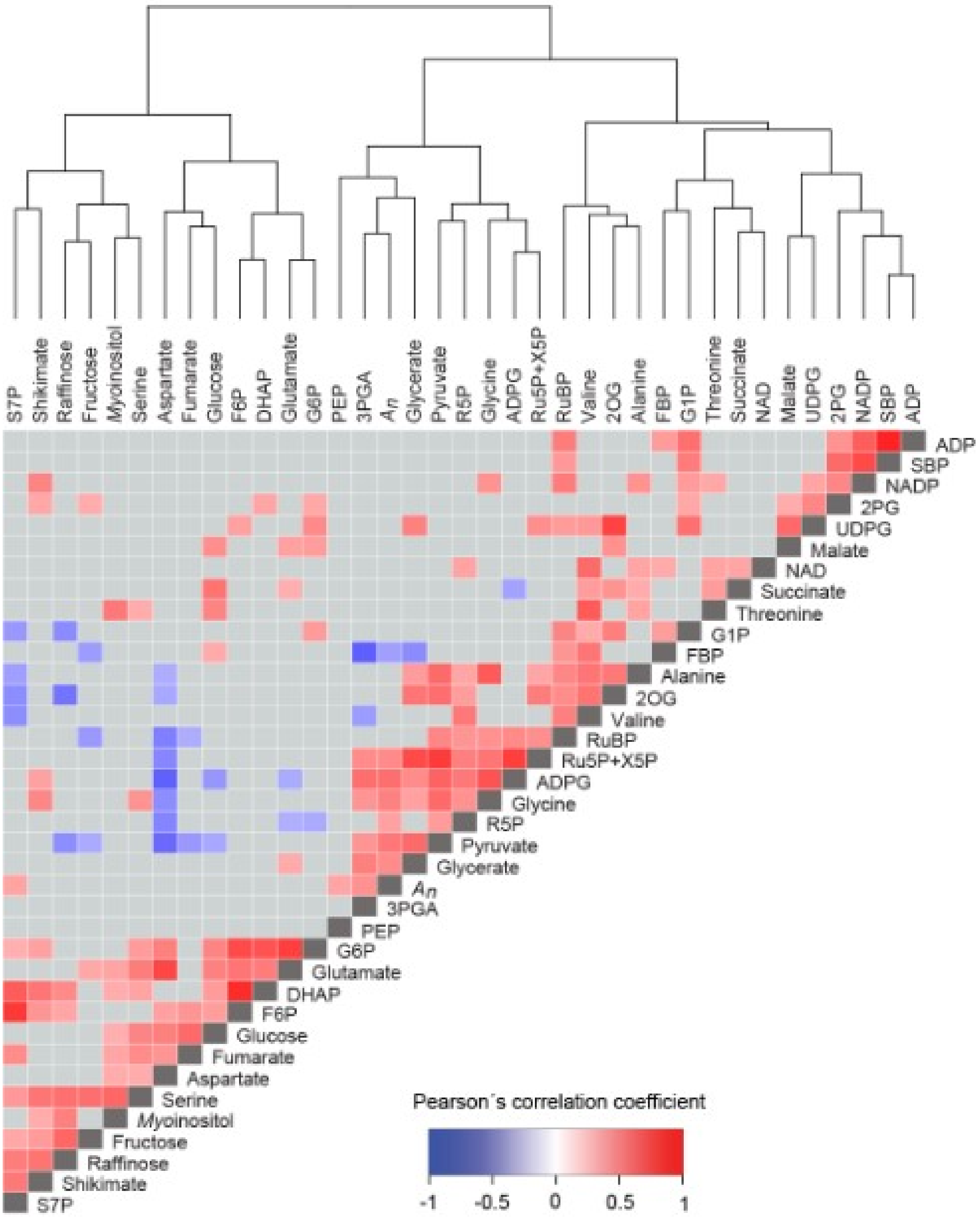

**Supplementary Figure S 6. Continued.**

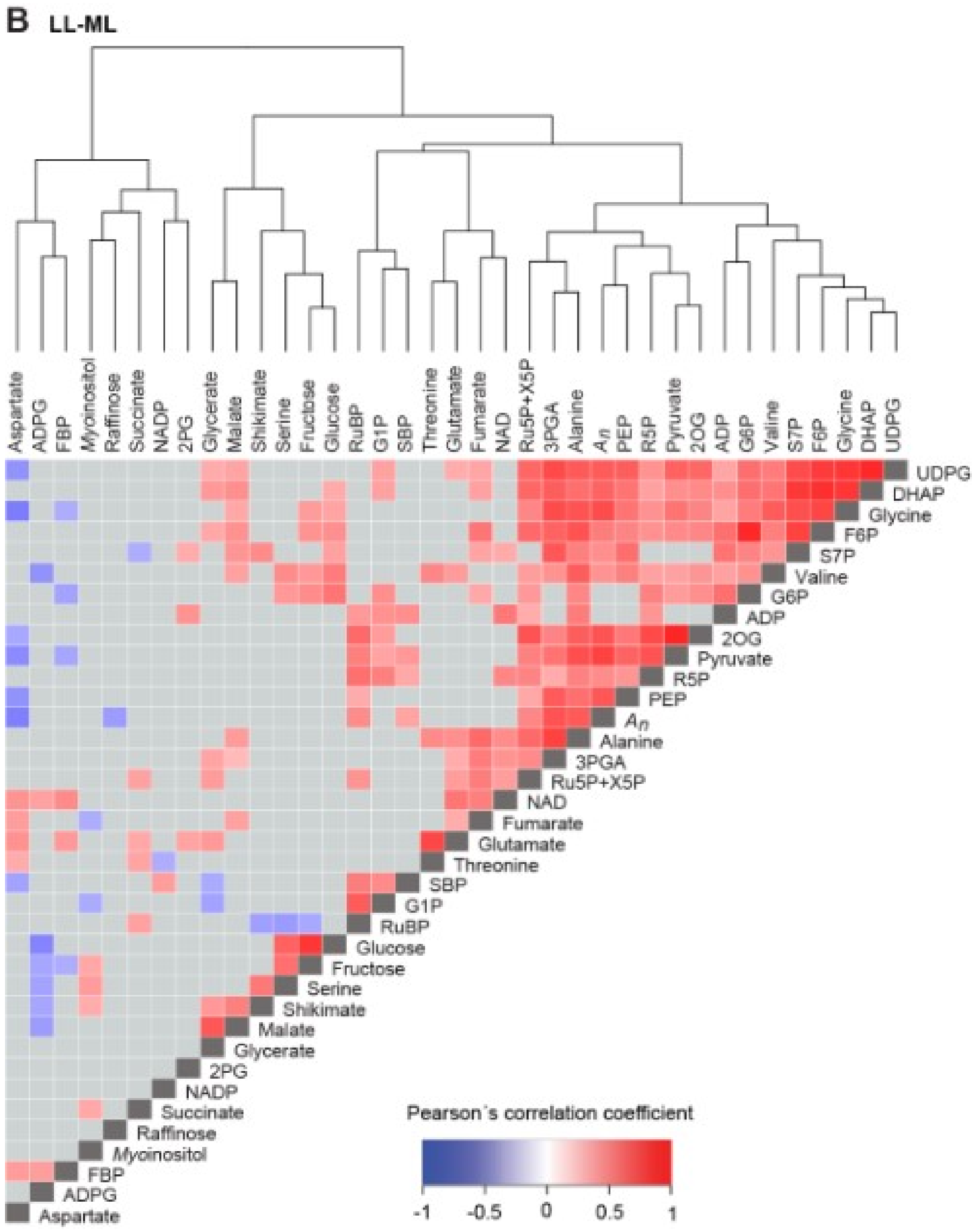

**Supplementary Figure S 6. Continued.**

(**C**) Relation between $A_n$ and metabolite levels or metabolic traits. $A_n$ was plotted against metabolite rates using individual samples ($n$=3 to 5 for ML-LL and $n$=4 for LL-ML) at all points after the transition (time zero excluded). Additional plots are calculated from 300 to 1800 s for the ML-LL transition, corresponding to the slight recovery of $A_n$ in the later part of the transient. The displays summarises $R^2$, the slope direction and p value (regression function with 95% confidence interval). The plots are provided in Supplementary Figs. S7 and S8.

(**D**) Relation between total metabolite pools in the CCM and the CBC. Analogous plots to panel C were made for summed C in metabolites in the CBC and CCM, or in the energy shuttle pools (3PGA and DHAP).

## C

| | ML-LL | | | | | | | LL-ML | | |
|---|---|---|---|---|---|---|---|---|---|---|
| | 5-1800 s | | | 300-1800 s | | | | 5-1800 s | | |
| Traits | $R^2$ | slope | $p$ | $R^2$ | slope | $p$ | | $R^2$ | slope | $p$ |
| PEP | 0.12 | + | 0.03 | 0.33 | + | 0.03 | | 0.44 | + | 0.00 |
| Malate | 0.01 | - | 0.48 | 0.02 | - | 0.61 | | 0.04 | + | 0.23 |
| Aspartate | 0.06 | - | 0.10 | 0.01 | + | 0.68 | | 0.24 | - | 0.00 |
| pyruvate | 0.24 | + | 0.00 | 0.31 | + | 0.02 | | 0.55 | + | 0.00 |
| Alanine | 0.02 | + | 0.43 | 0.30 | + | 0.02 | | 0.42 | + | 0.00 |
| Glutamate | 0.00 | - | 0.73 | 0.14 | + | 0.12 | | 0.01 | - | 0.50 |
| 2OG | 0.01 | + | 0.48 | 0.01 | + | 0.71 | | 0.45 | + | 0.00 |
| 3PGA | 0.18 | + | 0.00 | 0.56 | + | 0.00 | | 0.46 | + | 0.00 |
| DHAP | 0.00 | - | 0.70 | 0.04 | - | 0.41 | | 0.37 | + | 0.00 |
| FBP | 0.13 | - | 0.02 | 0.55 | - | 0.00 | | 0.08 | - | 0.08 |
| F6P | 0.02 | + | 0.35 | 0.03 | - | 0.48 | | 0.25 | + | 0.00 |
| SBP | 0.03 | - | 0.31 | 0.19 | + | 0.07 | | 0.13 | + | 0.02 |
| S7P | 0.12 | + | 0.02 | 0.00 | - | 0.92 | | 0.19 | + | 0.01 |
| R5P | 0.13 | + | 0.02 | 0.53 | - | 0.00 | | 0.20 | + | 0.00 |
| Ru5P + X5P | 0.21 | + | 0.00 | 0.13 | - | 0.14 | | 0.21 | + | 0.00 |
| RuBP | 0.00 | - | 0.70 | 0.31 | - | 0.02 | | 0.20 | + | 0.00 |
| ADPG | 0.32 | + | 0.00 | 0.11 | - | 0.17 | | 0.01 | + | 0.48 |
| 2PG | 0.01 | - | 0.58 | 0.00 | + | 0.97 | | 0.02 | - | 0.38 |
| Serine | 0.01 | + | 0.60 | 0.19 | + | 0.07 | | 0.02 | - | 0.38 |
| Glycine | 0.23 | + | 0.00 | 0.04 | + | 0.43 | | 0.48 | + | 0.00 |
| Glycerate | 0.20 | + | 0.00 | 0.21 | + | 0.05 | | 0.05 | + | 0.16 |
| 3PGA/DHAP | 0.15 | + | 0.01 | 0.34 | + | 0.01 | | 0.01 | - | 0.55 |
| FBP/F6P | 0.15 | - | 0.01 | 0.41 | - | 0.00 | | 0.18 | - | 0.01 |
| SBP/S7P | 0.06 | - | 0.12 | 0.13 | + | 0.14 | | 0.00 | - | 0.94 |
| Pentose-P/RuBP | 0.10 | + | 0.05 | 0.14 | + | 0.13 | | 0.03 | - | 0.29 |
| RuBP/3PGA | 0.05 | - | 0.14 | 0.45 | - | 0.00 | | 0.03 | + | 0.26 |
| RuBP/2PG | 0.00 | - | 0.76 | 0.01 | - | 0.63 | | 0.08 | + | 0.07 |
| Pyruvate/Alanine | 0.30 | + | 0.00 | 0.04 | + | 0.45 | | 0.12 | + | 0.03 |
| 2OG/Glutamate | 0.01 | + | 0.57 | 0.01 | - | 0.67 | | 0.40 | + | 0.00 |
| Sum C in CCM | 0.01 | + | 0.58 | 0.14 | + | 0.18 | | 0.20 | + | 0.00 |
| Sum C in Pyruvate + Alanine | 0.06 | + | 0.12 | 0.33 | + | 0.02 | | 0.50 | + | 0.00 |
| Sum C in CBC | 0.19 | + | 0.00 | 0.11 | + | 0.17 | | 0.55 | + | 0.00 |
| Sum C in CBC minus 3PGA + DHAP | 0.09 | + | 0.05 | 0.23 | - | 0.04 | | 0.39 | + | 0.00 |
| Sum C in 3PGA + DHAP | 0.16 | + | 0.01 | 0.45 | + | 0.00 | | 0.51 | + | 0.00 |
| Sum P in CBC | 0.17 | + | 0.01 | 0.19 | + | 0.07 | | 0.57 | + | 0.00 |

## D

| | | | | | | | | | | |
|---|---|---|---|---|---|---|---|---|---|---|
| C in CBC vs C in CCM | 0.00 | + | 0.90 | 0.03 | - | 0.57 | | 0.37 | + | 0.00 |
| C in CBC vs C in pyruvate + alanine | 0.03 | + | 0.28 | 0.03 | - | 0.53 | | 0.56 | + | 0.00 |
| C in 3PGA + DHAP vs C in CBC minus 3PGA + DHAP | 0.13 | + | 0.02 | 0.00 | - | 0.85 | | 0.45 | + | 0.00 |

| | p-value | | |
|---|---|---|---|
| | < 0.05 | < 0.01 | < 0.001 |
| UP | * | ** | *** |
| DOWN | * | ** | *** |

**Supplementary Figure S7. Additional plots of metabolite levels and metabolic traits against $A_n$ in a transition from moderate to low light.** This figure is Supplementary to Figs. 5-6.

(**A**) Metabolite levels**,**

**(B)** metabolite ratios, and

**(C)** sum of C in CCM and sums of C and P in CBC.

The design of the plots is described in the legend of Fig. 5. Briefly, the insert in shows times between 300-1800 s (dotted grey box in main panel), with an expanded scale for $A_n$ (52 – 56 µmol $CO_2$ $g^{-1}$ FW $s^{-1}$) to visualize relationships during the slight recovery of $A_n$ after 250 s. Arrows denote the time sequence in the main plot and in the insert. Slope directions and *p* values were calculated by linear regressions using individual samples at all time points after the transition (time zero excluded) and also from 300 to 1800 s (in insert). The *p* value of the correlation is given, colored according to the direction of the slope.

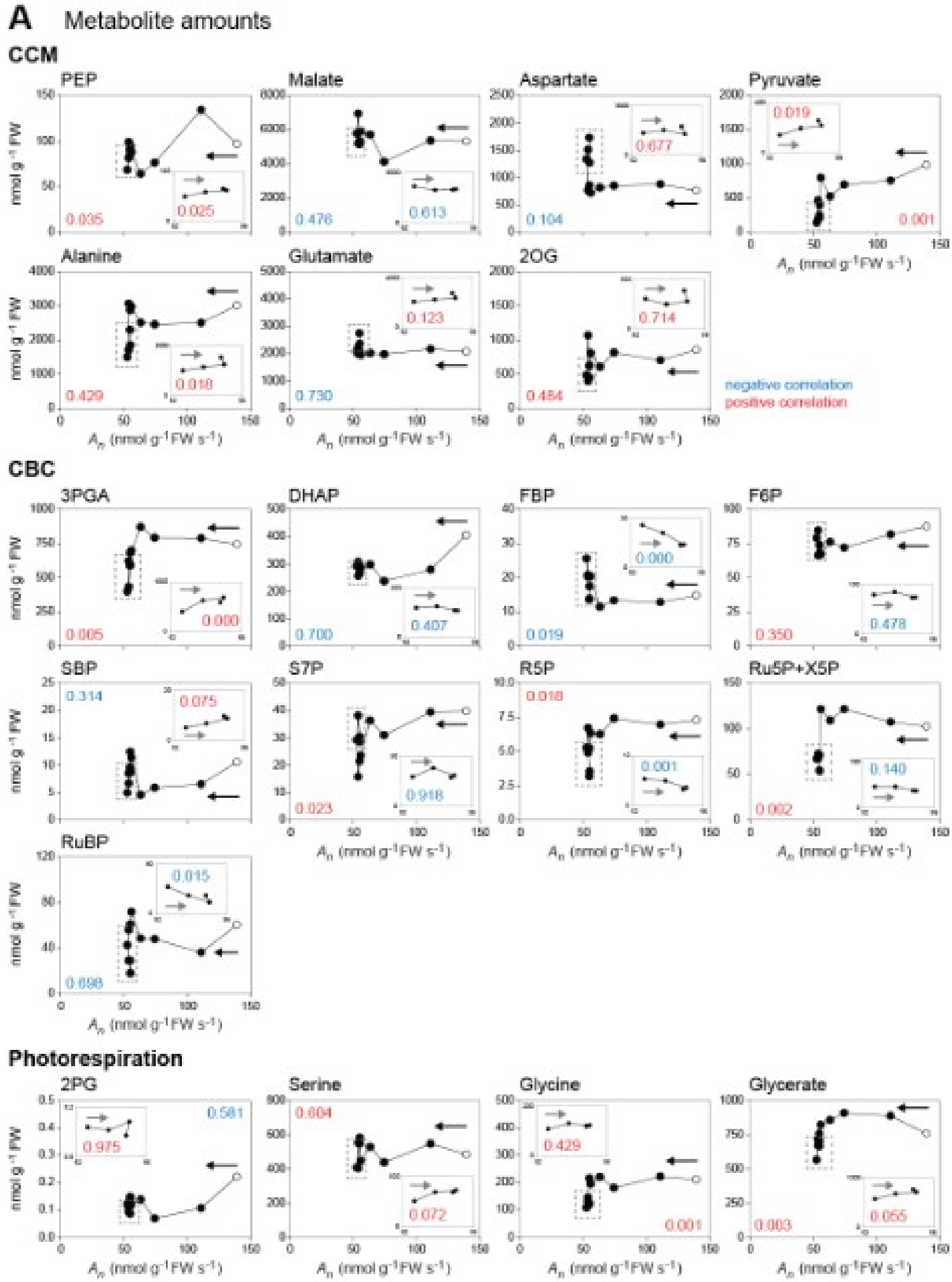

**Supplementary Figure S7. Continued.**

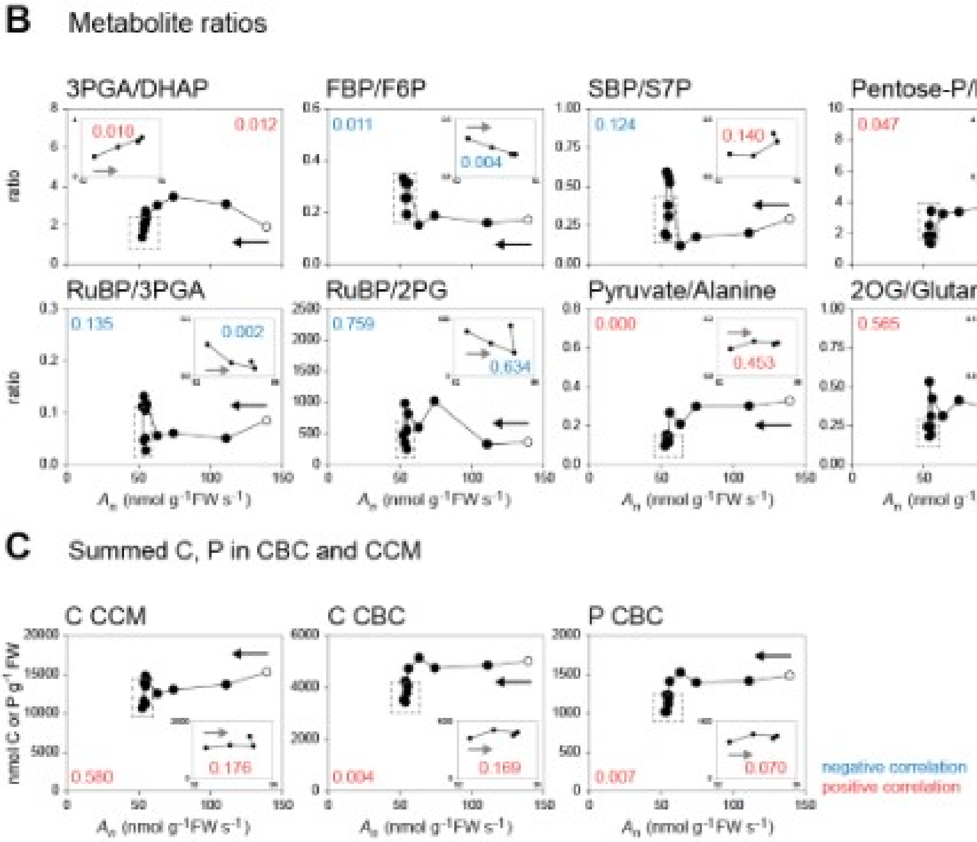

**Supplementary Figure S8. Additional plots of metabolite levels and metabolic traits against $A_n$ in a transition from low to moderate light.** This figure is Supplementary to Figs. 5-6.
(**A**) Metabolite levels,
**(B)** metabolite ratios,
**(C)** sum of C in CCM and sums of C and P in CBC.
The design of the plots is described in the legend of Fig. 5. Arrows denote the time sequence. Slope directions and *p* values were calculated by linear regressions using individual samples at all time points after the transition (time zero excluded). The *p* value is colored according to the direction of the slope.

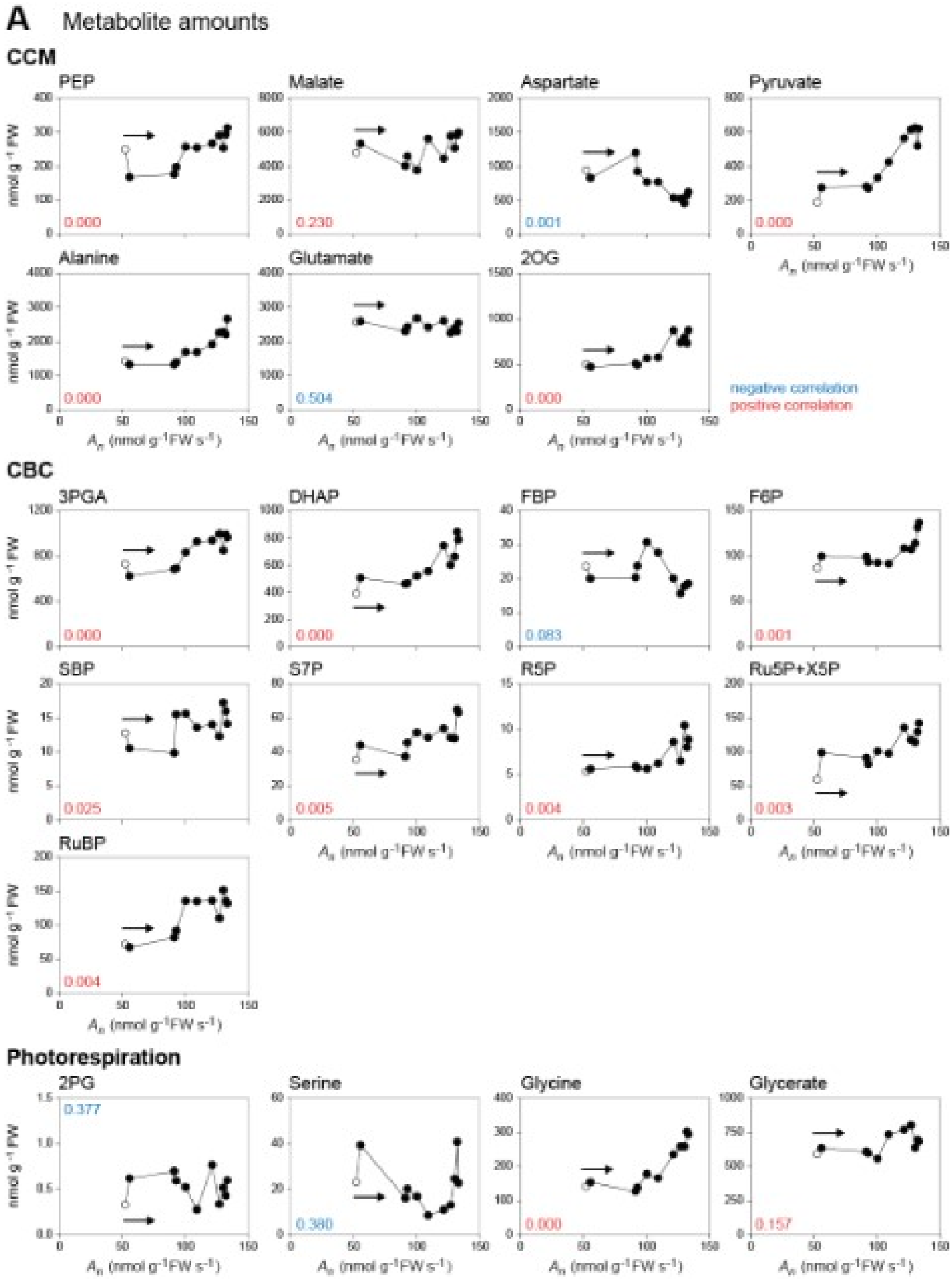

**Supplementary Figure S8.  Continued.**

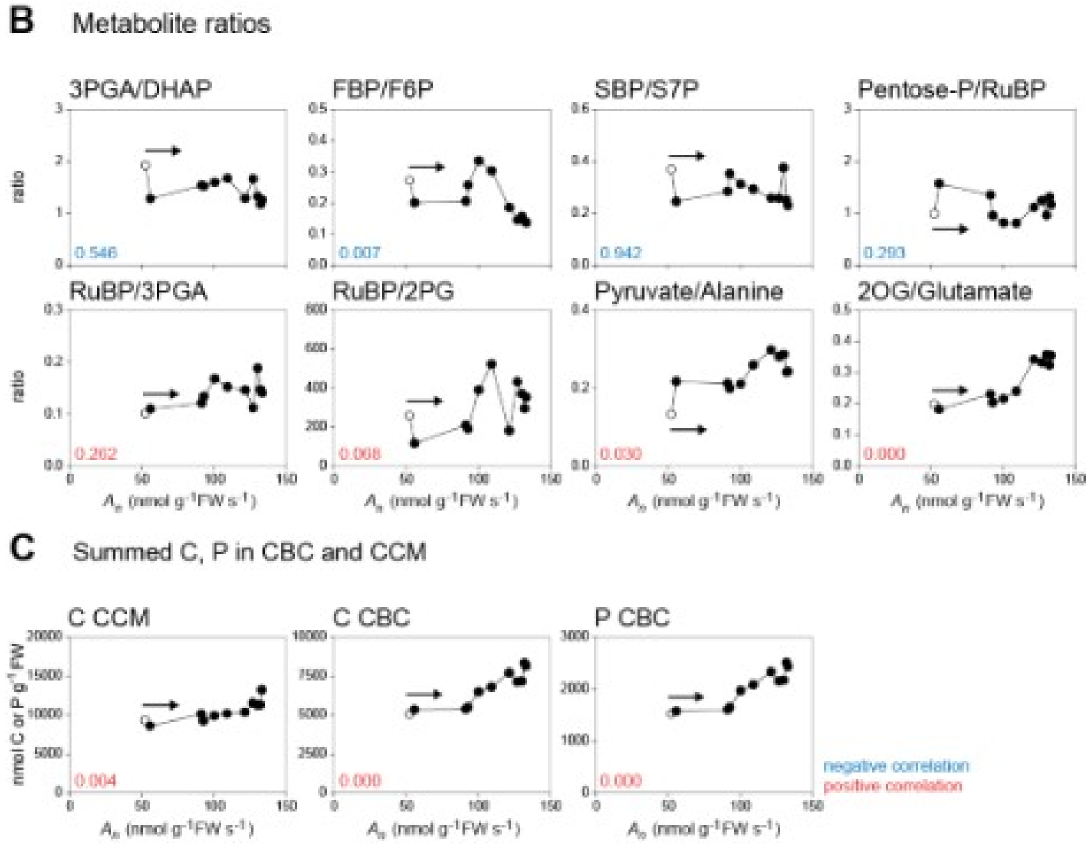

### Supplementary Text

**Contents** **Page**

**1. Gas exchange of plants sampled for metabolite measurements** 22

Supplementary to Results '*Response of $CO_2$ assimilation and stomatal conductance; Global analysis of response of metabolism'*

**2. Detailed account of changes in metabolism during the ML-LL and LL-ML transitions** 2 2

2.1. Supplementary to Discussion section '*Response of the CBC and CCM to a decrease in irradiance* 22

2.2. Supplementary to Discussion section '*Response of the CBC and CCM to an increase in irradiance'* 26

**3. Rate of exchange of C between the CBC (including the energy shuttle metabolites) and the CCM** 30

3.1. Background 30

3.2. Implications for movement of C between CBC and CCM pools in the ML-LL transition. Supplementary to Discussion section '*Response of the CBC and CCM to a decrease in irradiance* 33

3.3*.* Implications for movement of C between CBC and CCM pools in the LL-ML transition. Supplementary to Discussion section '*Response of the CBC and CCM to an increase in irradiance'* 34

**4. Changes in the contribution of different decarboxylation routes** 34

Supplementary to section '*Contribution of different decarboxylation routes'*

**5. Perturbation and adjustment of $C_{BSC}$** 38

Supplementary to the section '*Metabolite analysis point to rapid perturbation and slow adjustment of the $CO_2$ concentration in the bundle sheath'*

**6.Sequestration or recycling of C from metabolite pools in the photorespiratory pathway** 41

Supplementary to section '*Metabolite analysis point to rapid perturbation and slow adjustment of the $CO_2$ concentration in the bundle sheath*

**Additional references** 42

### 1. Gas exchange of plants sampled for metabolite measurements.

(Supplementary to sections '*Response of $CO_2$ assimilation and stomatal conductance', 'Global analysis of response of metabolism'*

The gas exchange data presented in Fig. 1 and Supplemental Dataset S1A were logged every second, and corrected by dynamic equations (Saathoff and Welles, 2021) to account for the lack of steady state. The effects of these corrections are shown in Supplementary Fig. S1J, revealing a clear impact of the correction just after the switch in light intensity for both transitions.

Metabolite measurements were made in samples harvested from separately-grown batches of maize. For technical reasons, gas exchange in this batch of plants was recorded at 6 s intervals and correction was not possible as each time point was a recorded running average (Supplementary Dataset S1B). Comparison of these data with the uncorrected data underlying Fig. 1 recorded every second showed good agreement between the two independent experiments (Supplementary Fig. 1K). In particular, in the ML-LL the initial decay of $A_n$ was delayed, and the trough at ~250 s and subsequent slight recovery of $A_n$ observed in the experiment where data were logged every second (Fig. 1C) was also observed in the experiment where data were recorded every 6 seconds (Supplementary Fig. S1K) with the recovery starting in this case from a slightly earlier trough at ~200 s (Supplementary Fig. S1L, significant at p= 0.3, paired t-test). In the LL-ML transition the slowing down of the rise at about 90 s and subsequent slow further rise until 1000 s was observed in both experiments (Supplementary Fig. S1K). The early plateau observed in the LL-ML transition at 11-15 s with corrected data (Fig. 1B) was visible as a slowing of the rise of $A_n$ in both uncorrected datasets (Supplementary (Fig. S1K).

### 2. Detailed account of changes in metabolism during the ML-LL and LL-ML transitions

#### 2.1 Supplementary to section '*Response of the CBC and CCM to a decrease in irradiance'*

The first phase of the ML-LL transition captured in the PC analysis (Fig. 2A) lasted until about 15 s. This coincides with the time when, instead of falling immediately, there is a progressive decline in $A_n$ (Fig. 1A; Supplementary Fig. 1C). In this time most CBC metabolites remained high and ADPG rose (Fig. 3A), which is consistent with continued operation of the CBC. The response in this phase confirms the prediction of Stitt and Zhu (2014) that after a decrease in irradiance the large pools of metabolites in the energy shuttle and the CCM temporarily buffer $C_4$ photosynthesis against the shortfall of ATP and NADPH from the light reactions.

Relatively high $A_n$ (over 50% of that in ML, or over 30% of the difference between ML and steady state LL, $\Delta A_n$,) was sustained until 10 s, and substantial rates until 15 s after switching to LL ($A_n$ over 40% of that in ML, or over 25% of $\Delta A_n$) (Fig. 1; Supplementary Fig. S1C). The response until 10 s is equivalent to buffering against the decrease in irradiance for about 4.5 s (Fig. 8A). This is a conservative estimate, because it excludes the continued slow decay of $A_n$ after 10 s (at 15 s it would be equivalent to buffering $A_n$ for about 6 s).

Previous studies in $C_4$ plants including maize reported that $CO_2$ assimilation continues for a time after darkening (Laisk and Edwards 1998), after imposing and then removing a short period of high light to imitate a sun fleck (Krall and Pearcy 1993) or in a fluctuating light regime after switching to low light (Lee et al. 2022; Arce-Cubas et al. 2023b).In a detailed study, Laisk and Edwards (1998) employed a custom-built gas exchange set up to investigate the dependence of post-illumination $CO_2$ assimilation on the rate and duration of photosynthesis in the preceding light treatment. They concluded that post-illumination $CO_2$ assimilation was driven by pools of metabolites that were built up in the light, argued that it involved flux at PEPC, and that the amount of $CO_2$ fixed after darkening was limited by the amount of PEP formed from triose-P and 3PGA. Our metabolite data including the maintenance of high CBC intermediate levels and ADPGl in the first 15-30 s after the decrease in light intensity (Fig. 3) show that not only PEPC but also the CBC remains more active than in steady state low light.

Continuation of $A_n$ at rates above those in steady state LL requires additional ATP and NADPH to that provided by the light reactions in LL. The additional cost until 10 s is equivalent to that required to support $CO_2$ fixation for 10 s at a rate of ~36 nmol $g^{-1}$ FW $s^{-1}$ (45% of $\Delta A_n$, where $\Delta A_n$ is 81 nmol $g^{-1}$ FW $s^{-1}$). Taking just costs in the CBC, this represents about 110 nmol ATP $g^{-1}$ FW and 72 NADPH nmol $g^{-1}$ FW. This will be an overestimate if some of the additional $CO_2$ fixed in first seconds in LL remains in $C_4$ acids, rather than being released and reassimilated in the CBC (see below). On the other hand, additional energy would be required if NADP-MDH and/or PPDK continue to operate for a short time at a higher rate than in steady state LL.

DHAP decreased by ~180 nmol $g^{-1}$ FW in the first 10 s after decreasing irradiance (Fig. 3A, see also Supplementary Fig. S3B). Assuming that the decrease is mainly due to oxidation of DHAP to 3PGA, it would generate up to 180 nmol $g^{-1}$ FW of ATP and NADPH. This is a maximum value because some DHAP will be consumed in other reactions, including continued use for RuBP regeneration by chloroplast FBPase, and continued conversion to sucrose by cytosolic FBPase. Nevertheless, the observed decline of DHAP is large enough to cover the estimated shortfall in ATP and NADPH in the first 10 or 15 s after the shift to LL.

Metabolism of malate might provide a further source of reducing equivalents. Measurements of total malate do not provide information about whether there is a decline of the malate pool that is directly involved in the CCM, so the rate of malate metabolism cannot be directly assessed. In principle, one possible route would be conversion of malate back to OAA by NADP-MDH, generating NADPH in the MC chloroplast. Another possibility is that, for a short time after the shift to LL, NADP-ME continues to convert malate to pyruvate and NADPH in the BSC at higher rates than in steady state LL. Laisk and Edwards (1998) argued that decarboxylation of malate by NADP-ME will be restricted after darkening because this reaction would rapidly reduce all the available NADP in the BSC chloroplast. The situation may be different after a switch to LL because NADPH consumption could continue for 3PGA reduction, using ATP produced by the BSC light reactions. That said, the downwards trend of pyruvate in the first 15 s after the decrease in irradiance indicates decreased flux at NADP-ME very soon after the switch to LL. Further, there was a rise of fumarate (Supplementary Figs. S3A-B) which, although not significant, is consistent with a small increase rather than a decrease of the metabolically active pool of malate. Together, these observations indicate that the NADP-ME reaction may not be a major source of NADPH after the transition to LL. Incidentally, the decline of pyruvate is unlikely to be due to increased PPDK activity, because PEP, the product of the PPDK reaction, declined slightly by 10 s (Fig. 3A; Supplementary Fig. S3B) and because lower irradiance will anyway lead to a shortfall of ATP in the MC.

It is possible that some of the $HCO_3$ fixed into OAA by PEPC in the first seconds after the transition to LL accumulates as $C_4$ acids that are not decarboxylated and therefore do not require a large energy input in the CBC. Accumulation as malate would still require one NADPH per C fixed. However, any shortfall of NADPH in the MC might lead to freshly synthesised OAA remaining as OAA or being converted to aspartate (see below for more discussion). There was no significant increase in aspartate in the first 15 s after decreasing irradiance (Fig. 3A; Supplementary Fig. S3B) but as the aspartate pool was large, it is possible that some C does accumulate in aspartate but this is masked by biological noise. If there is uncoupling of $HCO_3$ incorporation by PEPC and $CO_2$ assimilation in the CBC, this will decrease the energy required to support $A_n$ *i*mmediately after the transition to LL.

A second phase defined by the PC analysis started at about 15 s and lasted through to ~120 s (see Fig. 2A), and corresponded to most of the gradual decline of $A_n$ to a minimum, or trough, at about 250 s. This decline was accompanied by an increase of FBP and SBP levels (Fig. 3A; Supplementary Figs. S3B, S7A) The increase of FBP was not accompanied by an increase of DHAP, as might be expected if the aldolase reaction were near to equilibrium. This could reflect the complex intercellular compartmentation of these metabolites in maize leaves. Most of the DHAP is located in the MC and most of the FBP in the CBC in the BSC (Leegood

1985; Stitt and Heldt 1985a; Arrivault et al. 2017). It is possible that the overall DHAP pool mainly reflects the DHAP pool in the MC, the overall FBP pool mainly reflects that in the CBC in the BSC chloroplasts, and that DHAP in the BSC rises in parallel with FBP. Indeed, as $A_n$ falls, the size of the concentration gradient that is required to drive diffusion of DHAP from the MC to the BSC will decline, and this could include not just a decrease of the total DHAP pool (as seen in Fig. 3A), but also a decline of the concentration in the MC and maintenance or even an increase of the concentration in the BSC,

The increase of FBP and SBP levels (see above), maintenance of F6P and decline of S7P (Fig. 3A; Supplementary Figs. S3B, S7A) and the resulting increase of the FBP/F6P and SBP/S7P ratios (Fig. 3B; Supplementary Figs. S3B, S7B) point to inactivation of plastidic FBPase and SBPase. This may reflect falling availability of reducing equivalents. Doncaster et al (1989) reported that NADP-MDH activation, which is proxy for availability of reducing equivalents (Scheibe and Stitt 1988), decreased dramatically in the first minute after decreasing irradiance in maize. There was an increase of RuBP levels (Figs. 3A, 5B; Supplementary Fig. S7A) and of the RuBP/3PGA ratio and, though less consistently, the RuBP/2PG ratio (Fig. 3B; Supplementary Figs. S3B, S7B). These responses are consistent with decreased activation of Rubisco, possibly reflecting restriction of Rubisco activase by a shortfall of ATP (Portis and Parry 2007; Portis et al. 2008). The 3PGA/DHAP ratio, which rose immediately after the drop in irradiance (see above), declined between 15-120 s (Fig. 3B; Supplementary Figs. S3B, S7B), as expected as falling $A_n$ rebalances the relationship between provision of ATP and NADPH by the light reactions and their consumption in the CBC and CCM.

Overall, RuBP and many metabolites involved in its regeneration declined between 15 s and 120 s (Figs. 3A, 3C; Supplementary Figs. S3B, S7A, S7C). In this time span pyruvate decreased, but not PEP, alanine or aspartate (Fig. 3A; Supplementary Figs. S3B, S7B). Summed C in pyruvate and aspartate (Figs. 3C, 6B) and summed C in all CCM intermediates remained high for most of this phase (Fig. 3C; Supplementary Fig. S3B). Overall, the changes of CBC, energy shuttle and CCM metabolites were rather uncoordinated (for reasons see below, section ‘Rate of exchange of C between the CBC (including the energy shuttle metabolites) and the CCM’. The phase ended with a transient trough of $A_n$.

The third phase captured by the PC analysis started by 300 s (Fig. 2A and corresponded to a slow partial recovery of $A_n$ that was equivalent to almost 8 % of the difference in the rate of photosynthesis in ML and LL, and continued until ~1200 s. In experiments where maize was transferred from 1700 to 144 µmol $m^{-2}$ $s^{-1}$, Doncaster et al. (1989) observed a slight recovery of $A_n$ from about 2 min on. A similar trough and slow recovery was observed in many but not all previous studies of fluctuating light (Lee et al. 2022; Arce-Cubas et al. 2023b, see Discussion in main manuscript). In our ML-LL transition, the partial recovery of $A_n$ was accompanied by a decrease in FBP and RuBP levels (Fig. 3A; Supplementary Figs. S3B, S7A) and a decrease in FBP/F6P and RuBP/3PGA ratios (Fig. 3B; Supplementary Figs. S3B, S7B), consistent with activation of FBPase and Rubisco. SBP levels (Fig. 3A; Supplementary Figs. S3B, S7A) and the SBP/S7P ratio (Fig. 3B; Supplementary Figs. S3B, S7B) rose, indicating SBPase is regulated independently to plastidic FBPase. There was a ~20% increase in the total size of CBC pool (Fig. 3C; Supplementary Figs. S3B, S7C) due to a rise in the energy shuttle intermediates, 3PGA and DHAP (Figs. 3C, 6B; Supplementary Fig. S3B). There was also a marked rise in the levels of pyruvate, alanine and also aspartate (Figs. 3A, 6B; Supplementary Fig. S3B) resulting in a ~70% rise in the total C in CCM metabolites (Fig. 3C; Supplementary Figs. S3B, S7C). Overall, the C that accumulated in the CBC, energy and CCM pools was equivalent to about 10% of all the $CO_2$ fixed during the recovery phase, and more in the first part of the recovery.

#### 2.2 Supplementary to section '*Response of the CBC and CCM to an increase in irradiance*'

A sudden increase in irradiance was followed, within 5 s, by a decrease of the 3PGA/DHAP ratio. This immediate response presumably reflects increased availability of ATP and NADPH from the light reactions. It occurred before $A_n$ started to rise, at 5-6 s. Interestingly, the FBP/F6P and SBP/S7P ratios fell in the first 5-10 s after the increase in irradiance (Fig. 4B; Supplementary Fig. 4B; the decrease of the FBP/F6P ratio was significant and the decrease of both ratios was consistent across experiments, see Supplementary Fig. S5D). This points to rapid activation of FBPase and SBPase after an increase in irradiance. As in the ML-LL transient, DHAP changed in an opposite manner to FBP, possibly reflecting the intercellular location of these metabolites (see previous subsection).

The initial decline of 3PGA at 5 s was accompanied by a larger and significant decrease in PEP (Fig. 3A; Supplementary Fig. S3B). This rapid and overproportioned decrease of PEP is unlikely to be solely due to the decrease in 3PGA (see also section 3.1, below). Furthermore, pyruvate showed a small but significant increase at 5 s (Fig 3A; Supplementary Fig. S3B, resulting in a 2-fold increase in the pyruvate/PEP ratio (not plotted). This points to a temporary restriction on flux at PPDK. As shown by Chen et al (2014), PPDK is post-translationally regulated not only in response to dark-light switches, but also in response to changes in irradiance intensity. These observations also rose the question whether the decline in PEP may temporarily restrict flux at PEPC, especially as this enzyme is also not fully activated and the inactive form has lower affinity of PEP, decreased sensitivity to activation by activating metabolites, and higher lower sensitivity to inhibition by inhibitory metabolites (Ashton et al.1990; Doncaster and Leegood 1987; Vidal et al. 2002)

The subsequent increase of $A_n$ was spread over the next 10-15 min, with a rapid rise in the first 90 s (interrupted by a short plateau at ~10-15 s) and then a slower rise (Fig. 1B; Supplementary Figs. 1F-G).

The rise of $A_n$ from 6 s until about 90 s (Fig. 2B; Supplementary Fig. S1G) corresponded broadly to phase 2 of the PC analysis (Fig. 2B). It was associated with a rapid increase in CBC metabolite pools and enzyme regulation. There was a coordinated increase in 3PGA, DHAP, FBP, SBP, S7P, pentose-P and RuBP (Fig. 4A; Supplementary Figs. S4B, S8A). From about 10 s onwards, there was a decrease in the pentose-P/RuBP ratio (Fig. 4B; Supplementary Figs. S4B, S8B), pointing to rapid activation of PRK. After a drop at 5-10 s, the FBP/F6P ratio rose until 30 s and then fell (Fig. 4B; Supplementary Figs. S4B, S8B) pointing to activation of FBPase. The RuBP/3PGA and RuBP/2PG ratios rose (Fig. 4B; Supplementary Figs. S4B, S8B), suggesting that Rubisco is becoming increasingly restrictive for CBC flux.

This rise of $A_n$ was interrupted, slowing down by about 9 s and plateauing between about 11-15 s before starting to rise again (Fig. 2B; Supplementary Fig. S1G). A similar interruption was observed by Lee et al. (2022) after the shift to high light 120 s step fluctuating regime with 2 min for three $C_4$ species but not maize. In our LL-ML shift, the transient interruption was not accompanied by marked changes of metabolite levels, except for the preceding decrease in PEP and increase of the pyruvate/PEP ratio (see above), a decrease of the FBP/F6P ratio, a small increase of DHAP and pentose-P and a marked increase in ADPG (Fig. 4A; Supplementary Fig. S4B). Indeed, the 10 and 15 s time points, close to the start and end of the plateau, lay close to each other in the PC analysis (Fig. 2B). Nevertheless, the metabolite data allow exploration of potential explanations for this transient interruption in the rise of $A_n$.

Potential explanations include the need to increase delivery of $CO_2$ by the CCM, which would require associated changes in CCM intermediate pools and/or post-translational activation of enzyme or the need to increase $CO_2$ assimilation in the CBC by building up CBC pools and/or activating CBC enzymes (see Introduction and above). The decrease of PEP at 5-10 s points to one possible contributory factor being a shortfall in $CO_2$ pumping due to a delay in PPDK activation and restriction of PEPC by low PEP. This is, however, unlikely to be the only factor, as RuBP levels remained unchanged (Fig. 4A; Supplementary Fig. S4B) rather than rising as would be expected if there was a major shortfall of $CO_2$ in the BSC. The implication is that there is also a restriction on RuBP regeneration. The rise in ADPG (Fig. 4A; Supplementary Fig. S4B) might point to a loss of poise between CBC metabolites and Pi due, for example, to slow activation of sucrose synthesis, with falling plastid Pi levels allosterically activating ADPG pyrophosphorylase (Ballicora et al. 2004). However, the level of 3PGA (the allosteric activator of AGPG pyrophosphorylase) showed only a slight rise (at 10-15 s compared to 5 s, and there was, no clear evidence for a general rise in CBC metabolite levels immediately before or during the transient.

Another potential factor contributing to the interruption might be, for a short time, transient back-leakage of $CO_2$ from the BSC to the MC, due to decarboxylation of $C_4$ acids transiently exceeding the rate of $CO_2$ assimilation by the CBC. Bursts of $CO_2$ release have been reported in $C_4$ species including maize following reillumination after a short time in darkness (Krall and Pearcy 1993; Laisk and Edwards 1998). Indeed, Lee et al. (2022) discussed the transient interruption of $CO_2$ uptake in terms of a $CO_2$ burst. Some aspects of our metabolite data are consistent with this idea, especially when interpreted in terms of possible differing responses of metabolites in the BSC and MC. In the first 15 s in ML, 3PGA declined, there was a small non-significant increase in DHAP and an opposing upwards trend for FBP and SBP, as well as the FBP/F6P and SBP/S7P ratios (significant for the FBP/F6P ratio). The opposing changes of DHAP and FBP might be explained as follows: whilst in the MC the increased light intensity may immediately promote reduction of 3PGA to DHAP, this may not be the case for BSC where reduction of 3PGA to DHAP may be limited by availability of NADPH. Incidentally, a transient restriction on 3PGA reduction in the BSC might explain the marked and significant rise of ADPG at 5-15 s (Fig 4A, Supplementary Fig. 4B). At the same time, the rate of carboxylation of RuBP to produce 3PGA may be restricted. On the one hand, low DHAP, FBP, SBP (Fig. 4A) and delayed activation of FBPase (as revealed by the significant increase in the RBP/Fru6P ratio, Fig. 4B, Supplementary Fig. 4B) may restrict RuBP regeneration: in agreement, RuBP levels were initially unaltered and only rose after 15 s (Fig. 4A, Supplementary Fig. 4B). Delayed activation of Rubisco may additionally contribute to a short fall in $CO_2$ assimilation and production of 3PGA in the CBC. Crucially, the marked increase in the overall pools of DHAP and 3PGA does not start until about 30 s after the shift to ML, indeed at 5-15 s the overall 3PGA pool is lower than in LL (Fig. 4A, 4C, Supplementary Fig. 4B). This delayed increase of the 3PGA and DHAP pools will restrict the size of their intercellular concentration gradients and, hence, operation of the energy shuttle that is required to compensate for the NADPH deficit in the BSC. These factors could all contribute to restrict $CO_2$ assimilation in the BSC by the CBC, leading to leakage of $CO_2$ back to the MC and, if it is not all refixed by PEPC, to release of $CO_2$ from the leaf. Our metabolite data are indeed consistent with a restriction on PEPC activity; for example, PEP decreases by up to 40% in the first10 s in ML. The decline of PEP might partly reflect a restriction on flux at PPDK, as indicated by the small but significant rise in pyruvate at 5-15 s. However, it could also reflect conversion of PEP via enolase and phosphoglycerate mutase to 3PGA, which is then reduced to DHAP in the MC chloroplast (see also Sales et al. 2025). One other aspect that might be noted is that the initial significant increase of 2PG at 5 s was reversed and that the RuBP/2PG ration declined by 10 s; these are only trends but are consistent with the idea that an initial shortfall of $CO_2$ in the BSC 5 s after the shift to ML (see Supplementary Test Section 6 below) may be reversed by 10-15 s, possibly due to transient accumulation of $CO_2$ that is delivered by the CCM but is not yet being rapidly assimilated in the CBC

Overall, it is likely that multiple factors contribute to the transient interruption of the rise of $A_n$, which will partly mask the fingerprint of the individual factors. It is also likely that leaf-to-leaf variance complicates the comparison of $A_n$ and metabolite levels in this short transient. There was considerable leaf-to-leaf variation in $A_n$ during the plateau (Fig. 1B) due to different values of $A_n$ in the plateau and to the plateau starting and ending at slightly different times (Supplementary Fig. S1M). This is presumably accompanied by leaf-to-leaf variation in metabolite levels. Further, the half time of photosynthetic intermediates, especially those in the CBC are of the order of one or a few seconds (Stitt et al. 1980; Arrivault et al. 2009). Probably, a better understanding of the reasons for the transient interruption would be aided by denser time sampling of leaf material in which $A_n$ is being simultaneously monitored. That said, the most plausible explanation for the transient interruption in the rise of $A_n$ is that there is a delay in the rise in flux in the CCM and the rise in flux in the CBC, and that fluxes in the two cycles are transient imbalanced after a transition from LL to ML. The extent of the one or the other factor will probably vary, depending on prehistory and perturbation.

From about 90 s onwards, $A_n$ continued to rise, although more slowly (Fig. 2B; Supplementary Fig. S1G). A similar biphasic response was observed in maize and several other NADP-ME subtypes (*Setatia viridis*, *Setaria italica*, *Sorghum bicolor, Andropogon gerardii* Vitma, *Miscanthus* × *giganteus*, *Saccharum spp*., *Purus frumentum*) as well as NAD-ME species (*Amaranthus mangostanus*, *Panicum virgatum* L.) under a fluctuating light regime with a high light phase of 120 s or longer by Li et al. (2022) and Lee et al. (2023). On the other hand, a clear biphasic response was not observed in the PEPCK species *Alloteropsis semialata* (Arce-Cubas et al. 2023b) and *Spartina pectinata* L (Lee et al. 2023). Thus, with a possible exception of PEPCK subtypes, this two-phase response to an increase in irradiance may be a general feature of $C_4$ photosynthesis and occurs both in single transitions and fluctuating light. It represents a second reason for loss of photosynthetic C gain after an increase in light intensity.

In the maize LL-ML transition, the gradual rose of $A_n$ broadly corresponded to the third phase defined in the PC analysis (Fig. 2B). FBP fell and SBP, S7P (except at the very last times), pentose-P and RuBP remained unchanged (Figs. 4A, 5C; Supplementary Figs. S4B, S8A). The FBP/F6P ratio declined, the SBP/pentose-P ratio was fairly constant, the pentose-P/RuBP ratio rose, and the RuBP/3PGA and RuBP/2PG ratios did not show any consistent changes, pointing to small adjustments of poise within the CBC (Fig. 4B; Supplementary Fig. S4B). The most striking changes in this phase were a continuation of the rise of 3PGA and DHAP levels (Fig. 4A), and an increase in the levels of PEP, pyruvate and alanine (Fig. 4A; Supplementary Figs. S4B, S8A). This resulted in a marked increase in the summed pools involved in the energy shuttle and in the CCM (Figs. 4C, 6C; Supplementary Figs. S4B, S8C).

The gradual rise in $A_n$ correlated strongly with the increase in the pool size of the CBC, energy shuttle and CCM (Figs. 6A, 6C; Supplementary Figs. S4B, S8C). The CBC contributed in the first part of the transient mostly as $A_n$ rose from 50 to about 90 nmol $CO_2$ $g^{-1}$ FW $s^{-1}$, and the energy shuttle and CCM metabolites in the second part as $A_n$ rose from 90 to 130 nmol $CO_2$ $g^{-1}$ FW $s^{-1}$. This provides strong correlative evidence that the second and slower part of the rise in $A_n$ closely linked with the gradual build-up of the large metabolites pools that drive these intercellular shuttles. Overall, the rise in $A_n$ was accompanied by an increase of summed C in the CBC, energy shuttle and CCM of about 7000 nmol C $g^{-1}$ FW, which is equivalent to about 55 s of $A_n$ in ML, and considerably more of the lower $A_n$ that prevailed for the first part of the ML-LL transition.

### 3. Rate of exchange of C between the CBC (including the energy shuttle metabolites) and the CCM (Supplementary to both of the above sections)

#### *3.1 Background*

The CBC (including the energy shuttle intermediates 3PGA and DHAP) and the CCM are nominally independent pathways with no shared metabolites. However, exchange of C between the CBC and the CCM metabolites can occur via equilibration of 3PGA and PEP. These two metabolites are interconverted via near-equilibrium reactions catalysed by phosphoglycerate mutase and enolase. When operating in steady state, the CBC and CCM can form closed cycles. In the CBC, 3PGA is formed by Rubisco and either reduced in the BSC or moves to the MC where it is reduced. In the CCM, PEP is the substrate for PEPC, and is regenerated from pyruvate in the PPDK reaction. However, the near-equilibrium reactions catalysed by phosphoglycerate mutase and enolase provide the potential to move C between the CBC (and energy shuttle) and the CCM (see diagram overpage, the steady state corresponds to the scenario in the left-hand panel). This potential to interconvert 3PGA and PEP is underlined by the higher activity of phosphoglycerate mutase and enolase in the MC compared to the BSC reported in Furbank and Leegood (1984).

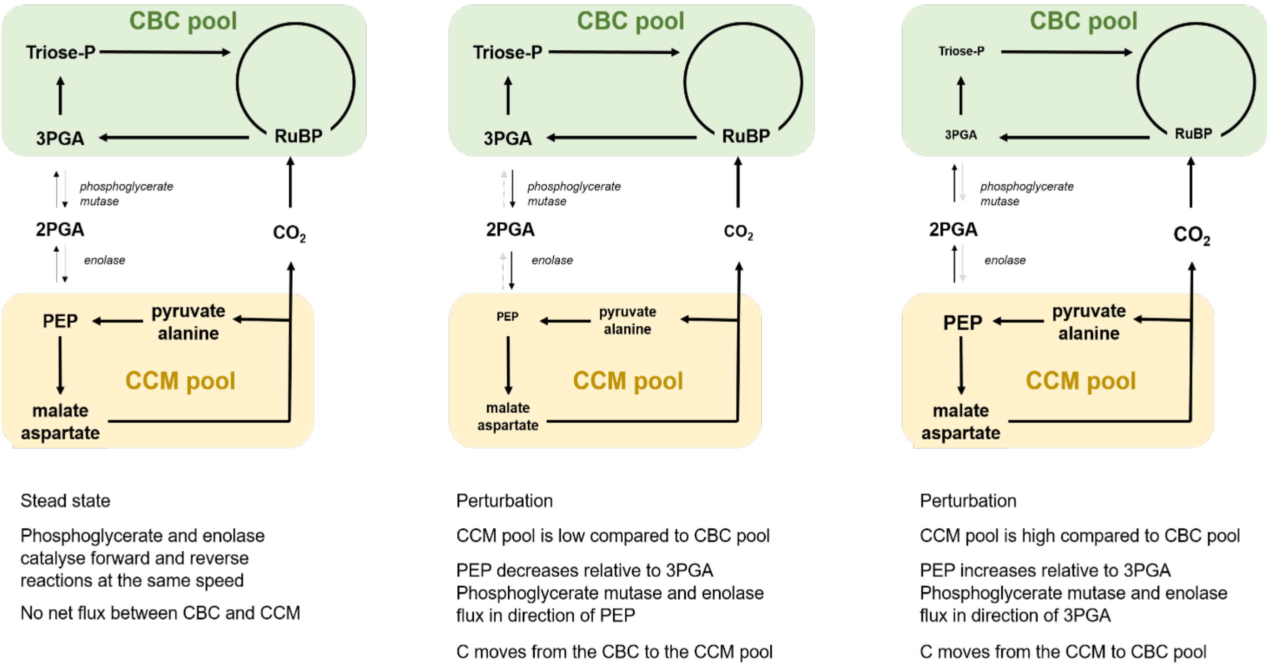


Based on $^{13}C$ labelling kinetics, Medeiros et al. (2022) recently estimated that during steady state photosynthesis phosphoglycerate mutase and enolase exchange $^{13}C$ at a rate equivalent to 18 and 22% of the net rate of $CO_2$ fixation at an irradiance of 550 and 160 µmol$^{-2}$ m$^{-2}$ s$^{-1}$, respectively. Exchange of label does not, however, mean that there is a net flux of C. At thermodynamic equilibrium, when no net flux occurs, the 3PGA:PEP ratio would be about 3:1 (based on the theoretical equilibrium constants of phosphoglycerate mutase and enolase of 0.17 and 2.07, Stryer 1990). This may be modified by the conditions in the cellular milieu like pH and $Mg^{2+}$ levels, as these differentially affect ionization and ion binding and thus the concentration of the actual substrate. Further, due to compartmentation, the overall level may not precisely reflect the cytoplasmic level in the MC. The first reason could be differing distribution of 3PGA and PEP between the BSC and MC. The second and probably more important reason is subcellular distribution between the chloroplast stroma and cytosol. Due to the absence of phosphoglycerate mutase and enolase from the plastid stroma of chloroplasts in mature leaves (Stitt and ap Rees 1980; Fukuyama et al. 2015), there are substantial amounts of 3PGA but only very low PEP in the stroma (Szecowka et al. 2013). As a result, the overall 3PGA/PEP ratio in leaf material will overestimate the ratio in the MC cytosol, where interconversion of 3PGA and PEP will directly impact on the C flow towards PEPC. Detailed studies of metabolite levels across a range of conditions in maize and Amaranthus in which 3PGA and PEP levels varied over a <10-fold range revealed that the relationship is maintained very constant, with a 3PGA:PEP ratio of about 4.6 (Leegood and von Caemmerer 1989). In our current studies, the measured levels 3PGA and PEP in maize leaves in steady state condition were close to or above the theoretical 3:1 ratio (Figs. 3A, 4A). Thus, whilst labeling studies demonstrate there is rapid movement of C from 3PGA into PEP

(Medeiros et al. 2022), in steady state conditions this will be largely balanced by flow of C from PEP back to 3PGA. There will be only slow net conversion of 3PGA to PEP to replace any PEP that is consumed by PEPC in excess of the PEP that is recycled in the CCM by PPDK (for example, to cover any flux out of the TCA for net amino acid synthesis).

Net flux of C will occur after perturbations that alter the rate of photosynthesis and/or the balance between CBC and CCM pool size. After perturbations that increase the level of 3PGA relative to PEP (or more precisely, when the 3PGA/PEP level in the MC cytosol rises above the ratio found at thermodynamic equilibrium) there will be an increase in next flux from 3PGA to PEP (middle scenario in above sketch). After perturbations that decrease in the level of 3PGA relative to PEP (or more precisely, when the 3PGA/PEP level in the MC cytosol falls below the ratio found at thermodynamic equilibrium) there will be net flux from PEP to 3PGA (right hand scenario in above sketch).

Based on the rate of label exchange in steady state photosynthesis in Medeiros et al. (2022), and estimated for the most extreme scenario:

i) when the 3PGA/PEP ratio is very high and phosphoglycerate mutase and enolase catalyze almost exclusively reactions in the direction that converts 3PGA to 2PGA to PEP, this flux could generate a ~10% decrease in the total CBC pool (including 3PGA and triose-P) in about 25 s under an irradiance of 550 µmol $m^{-2}$ $s^{-1}$, with slightly longer being required under an irradiance 160 $µmol^{-}$ $m^{-2}$ $s^{-1}$.

ii) when the 3PGA/PEP ratio is very low and enolase and phosphoglycerate mutase catalyze almost only reactions in the direction that converts 3PEP to 2PGA to 3PGA, the flux could generate a ~10% increase in the total pool of CCM metabolites in about 36 s under an irradiance of 550 $µmol^{-2}$ $m^{-2}$ $s^{-1}$, with slightly longer being required under an irradiance 160 $µmol^{-2}$ $m^{-2}$ $s^{-1}$.

These calculations are based on measured values of $A_n$ in ML (~ 135 nmol C $g^{-1}$ FW $s^{-1}$) and the summed pool of CCM metabolites and summed pool of CBC plus energy metabolites (8200 and 6250 nmol C $g^{-1}$ FW, respectively).

It should be noted that the summed pool for CCM metabolites is underestimated because it does not include malate (the active pool of malate cannot be well determined due to the presence of two or more pools of malate in maize leaves, with the pool that is directly involved in the CCM being only a small component, probably less than 20% (see Arrivault et al. 2017; Medeiros et al. 2022). Consequently, the estimated time for a ~10% increase in the total pool of CCM metabolites of about 36 s is a minimum value.

#### 3.2 Implications for movement of C between CBC and CCM pools in the ML-LL transition

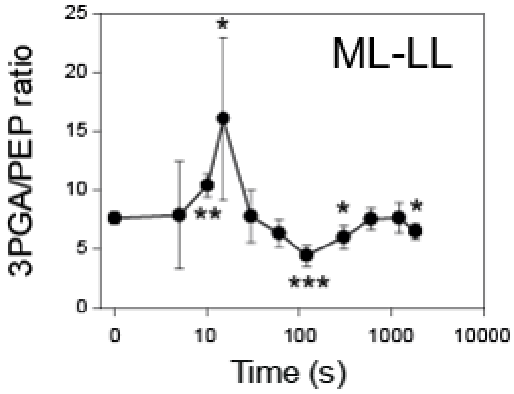


During a ML-LL transition, the summed C in the pool of CBC plus energy metabolites fell in the first 120 s (Fig. 3C), followed slightly later (120-300 s) by a fall in the summed C in the estimated pool of CCM metabolites, especially pyruvate and alanine (Figs. 3A, 3C). Interestingly, this slightly delayed drop of the CCM metabolites started after a delay. This corresponded with changes of 3PGA and the 3PGA/PEP ratio. 3PGA remained high at 10-15 s and even rose relative to PEP (compare plots in Fig. 3A, also plot of the 3PGA/PEP ratio alongside, T-test indicated as in Fig. 3). From 30 s onwards, 3PGA decreased both in absolute terms (Fig. 3) and relative to the PEP (Fig. 3 and plot alongside). The delayed and slow decay of the CCM pool was probably due at least in part to this delay until 3PGA decreased, and to the 3PGA/PEP ratio only decreasing about two-fold compared to steady state ML or LL (Fig. 3A, see plot alongside) as well as the rather restricted capacity for flux over phosphoglycerate mutase and enolase (see previous subsection). This may explain why the responses of the CBC pool, the energy metabolite pool and the CCM pool were rather uncoordinated during the ML-LL transition (Fig. 7B). Nevertheless, by 300 s there was a substantial decrease in both the CBC and the CCM pools (by 1500 and ~5000 nmol C $g^{-1}$ FW, respectively, accounting for ~30 and ~33%, respectively, of the initial pool size in ML; see Fig. 1C and Supplementary Dataset S2).

As described in the main text and Supplementary Text Section 2.1, the initial decline of CBC and CCM metabolite pools is partly reversed later in the ML-LL transition. The slight (7%) but highly significant rise in $A_n$ (Fig. 1C) was accompanied by a ~20% increase in the total size of CBC pool, and a ~70% rise in the C in CCM metabolites (Fig. 3C; Supplementary Fig. S3B). As also mentioned in Supplementary Text Section 2.1, the partial recovery of the CBC and CCM pools together accounted for about 10% of the $CO_2$ fixed during the recovery phase. This increase was mostly due to the rise in the CCM pool (from 10,339 to 14,828 nmol C $g^{-1}$ FW) compared to a smaller and possibly earlier rise of the CBC metabolite pool (from 3,443 nmol C $g^{-1}$ FW at 120 s to a peak of 4,252 nmol C $g^{-1}$ FW at 600 s, that was followed by a plateau or slight decline) (Fig. 1C, for details see Supplementary Dataset S2). Thus, during this recovery, over 8% of the fixed C is moved from the CBC pool via phosphoglycerate mutase and enolase into the CCM pool. It is noteworthy that during this slight recovery, the 3PGA level rises more markedly than the PEP level (see Fig. 3A and insert above), which would favour increased net flux from the CBC pool to the CCM pool. The relatively small rise in PEP between 300-1800 s may also imply that PEPC activity has been upregulated.

Thus, CCM and CBC metabolite pools change in an uncoordinated manner in the ML-LL transition, both i) early in the transition as $A_n$ is rapidly decreasing, when the CCM pool declines later than the CBC pool and ii) later in the transition as $A_n$ recovers slightly, when the CCM pool rises later but more markedly than the CBC pool. In both cases, the differing response can be at least partly explained by changes in the 3PGA/PEP ratio that, on the one hand are indicative of an imbalance between the CCM and CBC and, on the other hand, determine the net direction and rate of C exchange between the CBC and CCM.

**3.3 *Implications for movement of C between CBC and CCM pools in the LL-ML transition.***

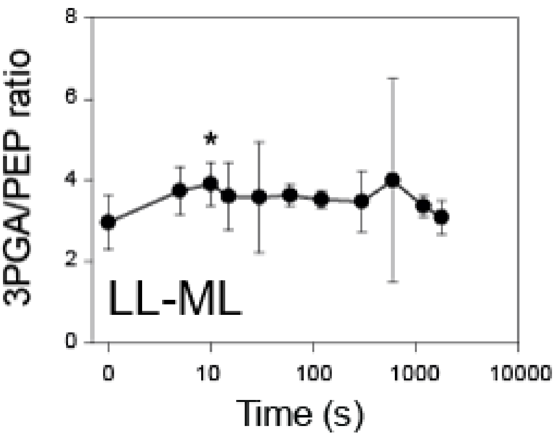


During an LL-ML transition, the summed C in the pool of CCM metabolites and the summed C in the pool of CBC plus energy metabolites rise gradually (Fig. 4C), with the CBC pool (excluding 3PGA and DHAP) rising in the first part of the response (until about 100 s) and the energy shuttle and CCM metabolites increasing in the later part of the response from about 100 s onwards (Fig. 4,6C; Supplementary Fig. S4B; Supplementary Text Section 2.2). Thus, in contrast to the ML-LL transition, these pools change in a rather coordinated manner (see Fig. 7). After an initial rapid decrease, the 3PGA and PEP levels rise gradually, and roughly in parallel with  small (~30%) significant increase at 10 s (Fig. 4A, see also plot of 3PGA/PEP ratio alongside, note the y-axis scale is smaller than for the plot of the ML-LL transition). This rather stable 3PGA/PEP ratio is consistent with the rather coordinated rise in the CBC and CCM pools during the LL-ML transition.

### 4. Contribution of different decarboxylation routes

Although $C_4$ plants are often subdivided into NADP-ME, NAD-ME and PEPCK sub-types based on their main route for decarboxylation, in many species including maize different decarboxylases operate concomitantly (Furbank 2011; Bräutigam et al. 2014; Wang et al. 2014a). Maize has substantial expression (Furomoto et al. 1999; Pick et al, 2011; Wang et al. 2014c) and activity of PEPCK in the BSC (Walker et al. 1997). In this, maize resembles other NADP-ME subtypes. In maize bundle sheath preparations, whilst decarboxylation of malate occurred via NADP-ME leading to formation of pyruvate, decarboxylation of aspartate occurred via PEPCK leading to formation of alanine (Wingler et al. 1999). The proportion of $CO_2$ carried by aspartate-based routes like PEPCK in moderate light was estimated from analysis of labelling kinetics in leaves as ~10% wild-type maize (Arrivault et al., 2017) or as high as 25% in a comparison of wild-type maize and the *dct2* mutant (Weissmann et al. 2016).

The contribution of different decarboxylation routes may depend on irradiance. Analyses of steady state metabolite levels in maize in steady state low irradiance and high irradiance revealed with higher levels of aspartate and alanine relative to pyruvate under low irradiance, pointing to an increased contribution of PEPCK at low irradiance (Usuda 1987; Leegood and von Caemmerer 1989). A similar conclusion was reached from $^{13}CO_2$ labelling studies in maize at low and moderate irradiance, with substantial labelling of aspartate at low irradiance and lower labelling at moderate irradiance, whilst malate showed an opposite pattern (Medeiros et al. (2022). Further, aspartate increased in maize leaves of maize and other $C_4$ species during adjustment to a large sudden decrease in irradiance (Doncaster et al. 1989).

In our current study of irradiance transitions, aspartate showed a strikingly different pattern to most other metabolites (Fig. 5; Supplementary Figs. S6A-B). Aspartate levels rose significantly during the adjustment to low irradiance (ML-LL, Fig. 3A; Supplementary Figs. S3B, Fig. S7A), whereas most other metabolites declined. Aspartate levels decreased significantly during the adjustment to higher irradiance (LL-ML, Fig. 4A; Supplementary Figs. S4B, S8A), whereas most other metabolites increased. In both cases, the change in aspartate levels occurred in the later part of the response, from ~120 s onwards. These observations indicate that the contribution of decarboxylation routes other than NADP-ME may increase in the recovery phase after a transition to lower irradiance, and may decrease in the later part of the adjustment to higher irradiance.

One possible explanation for the increase of aspartate in low light is that any shortfall of NADPH in the MC would, via mass action, restrict conversion of OAA to malate, and favor conversion of OAA via aminotransferase to aspartate. A reciprocal scenario might occur after an increase in irradiance. Such effects would, however, be expected to occur immediately after the change in irradiance. The changes of aspartate level occurred relatively slowly during the transitions (Figs. 3A, 4A). The slow response indicates either that redox-driven changes in metabolite levels do not play a large role, or that conversion of OAA to aspartate also depends on other factors, which change more slowly. One might be the availability of amino donors. This explanation is supported by the observation that the 2OG/glutamate ratio and the pyruvate/alanine ratio were high in moderate irradiance and remained high in the first part of the adjustment to low irradiance and that both ratios were high in low irradiance and declined gradually after transfer to higher irradiance (Figs. 3B, 4B). Another contributory factor may be post-translational regulation of NADP-MDH. Activation and inactivation of NADP-MDH requires up to 10 min after illuminating or darkening maize chloroplasts (Rebeille et al. 1986;

Ashton et al. 1990). If the changes in activation and deactivation are similarly slow in maize leaves after changes in irradiance, this might contribute to the delayed response in the levels of aspartate in the ML-LL and LL-ML transitions (see also below for further discussion) although probably being less relevant in fluctuating light

Movement of aspartate from the MC to the BSC will need to be coupled to movement of an amino acid back from the BSC to the MC to maintain nitrogen stoichiometry. In the most parsimonious pathway, this would involve movement of alanine (Weissmann et al. 2016; Bräutigam et al. 2018). In the ML-LL transition the rise of aspartate was accompanied by a significant decrease of the pyruvate/alanine ratio (Fig. 3B; Supplementary Figs. S3B, S7B) but this was due mainly to the decrease of pyruvate being larger than the decrease of alanine (Fig. 3A). In the LL-ML transition, the decline of aspartate was accompanied by a significant increase in the pyruvate/alanine ratio (Fig. 4B; Supplementary Figs. S4B, S8B) but this was due mainly to the increase of pyruvate being larger than the increase of alanine (Fig. 4A). In both irradiance transitions, for most of the time the levels of aspartate and alanine were correlated negatively rather than positively (see Fig. 3A, 4A, 5A; Supplementary Figs. S6A, S6B). Whilst there was a positive correlation between aspartate and alanine in the last part of the transition to low light, this was also accompanied by a rise in pyruvate and might just reflect a general increase in the pool of 3C metabolites (Fig. 3A, see also Discussion section 'Response of the CBC and CCM to a decrease in irradiance' and Supplementary Text Section 2.1). In studies of light-dark transitions in maize leaves, Furbank and Leegood (1984) reported that aspartate and alanine levels decline in the first 5 min after illumination, but Usuda (1985) reported that aspartate transiently declined and recovered whereas alanine transiently rose and then decreased. Taken together, these observations argue against operation of a tight obligatory intercellular shuttle between aspartate and alanine, as this would require parallel changes of both metabolite pools to drive a coordinated change in the intercellular movement of both metabolites.

An alternative scenario would be that some of the amino groups from aspartate return from the BSC to the MC as another amino acid. One possibility is that glutamate and 2OG provide a secondary shuttle to maintain nitrogen stoichiometry (Mallmann et al. 2024; Medereiros et al. 2022). The levels of both were high and changed markedly during our sudden transients. During adjustment to low light, aspartate correlated positively with glutamate (Supplementary Fig. S6, compare Fig. 3A and Supplementary Fig. 4B) and negatively with the 2OG/glutamate ratio (compare Fig 3A and Fig. 3B). During adjustment to an increase in irradiance, the aspartate level declined (Fig. 4A; Supplementary Fig. 4B) and the 2OG/glutamate ratio rose (compare Fig 4B, Supplementary Fig. 4B). Interestingly, one of the metabolic traits that changed markedly during the optimisation of $C_4$ photosynthesis between $C_4$-like and true $C_4$

species in the *Flaveria* genus was that the levels of 2OG and glutamate increased (Borghi et al. 2019). A secondary nitrogen shuttle might provide added flexibility to $C_4$ photosynthesis, both in avoiding restrictions that a shortfall of one single metabolite might place on the rate of intercellular movement and thence the rate of photosynthesis, as well as by decreasing the required pool size for any individual metabolite because intercellular movement is distributed across several metabolites (see also Furbank 2011; Wang et al. 2014a; Bellasio and Griffiths 2014a).

Several factors might contribute to the increased contribution of aspartate-based routes like PEPCK to decarboxylation in low irradiance. On the one hand, operation of the NADP-ME route may be constrained in low light (see Medeiros et al. 2022). Several factors might potentially restrict operation of this decarboxylation route in low irradiance. They may include, in the MC, any restriction on conversion of OAA to malate due to low availability of NADPH or incomplete activation of NADP-MDH. Post-translational activation of NADP-MDH by thioredoxin is favored by a rising NADPH/NADP ratio as the light intensity rises in $C_3$ plants (Scheibe and Stitt 1988; Scheibe 1990; Knuesting and Scheibe 2018) and $C_4$ plants (Ashton and Hatch 1983; Rebeille et al. 1986; Ashton et al. 1990). As a result, high rates of OAA reduction in $C_4$ plants can only occur at high NADPH/HADP ratios (Ashton and Hatch 1983; Rebeille and Hatch 1986; Ashton et al. 1990). Incidentally, it was shown recently using Arabidopsis lines expressing constitutively active NADP-MDH that inactivation of NADP-MDH in low irradiance is important for photosynthetic performance of $C_3$ plants under fluctuating light (Yokochi et al. 2021). By extrapolation, inactivation and activation of NADP-MDH may also be important during light transitions in $C_4$ photosynthesis. Irrespective of the reasons, any restriction on synthesis of malate in low irradiance will presumably hinder the generation of a high concentration of malate in the MC and slow down movement of malate by diffusion to the BSC. In the BSC, several factors may restrict decarboxylation by NADP-ME in low irradiance, including a lack of demand for the products of NADP-ME (Hatch and Kagawa 1976; Bräutigam et al. 2018), incomplete light-activation of NADP-ME (Bovdilova et al. 2019) and/or poor transport of pyruvate back to the MC (see Medeiros et al. 2022 for discussion). On the other hand, operation of the PEPCK route may be favored at low irradiance compared to high irradiance. Maize PEPCK is not subject to post-translational light activation and is active in the dark (Wingler et al. 2019; Walker et al. 2002) and is presumably as active under low irradiance as in high irradiance. Indeed, moderate to high light may even lead to phosphorylation and inactivation of maize PEPCK (Chao et al. 2014).

A further factor that may allow a larger contribution of PEPCK under low irradiance relates to movement of PEP from the BSC back to the MC. It is an open question how this occurs in PEPCK subtypes (Braütigam et al. 2018) and analogous issues arise when PEPCK is operating in parallel with for example, NADP-ME. The rate of movement of PEP will be restricted by the relatively small size of the PEP pool (see Figs 3A, 4A), which restricts the size of the intercellular concentration gradient that can be generated to drive diffusion of PEP from the BSC to the MC. Further, movement by this route will also be constrained by the need to maintain high enough PEP levels in the MC to support rapid flux at PEPC. The relatively low concentration of PEP reflects the thermodynamics of the enolase and phosphoglycerate mutase reactions, which favor formation of 3PGA and result, at equilibrium, in an approximately 3-fold excess of 3PGA over PEP (Supplementary Text, Section 3 above). In principle, a thermodynamically more favorable route to return PEP to the MC would be to convert PEP to 3PGA in the BSC, followed by diffusion of 3PGA down its larger concentration gradient to the MC and conversion of 3PGA back to PEP in the MC. However, this route would require activities of enolase and phosphoglycerate mutase that are well in excess of flux at PEPCK in order to maintain near-equilibrium concentrations of PEP and 3PGA in the BSC and in the MC. As discussed in above (Supplementary Text, Section 3), flux between 3PGA and PEP in maize is high enough to allow rebalancing of pool sizes in the CBC and CCM, with about 10% of the C in these pools being able to move in about half a minute. It is however, not commensurate with carrying a net flux that exceeds flux at PEPCK, or even a fraction of this in the scenario where PEPCK makes a small contribution to the CCM. Further this route would require enolase and enolase and phosphoglycerate mutase in both the BSC and the MC, rather than an asymmetric distribution with the majority in the MC as is the case in maize (Furbank and Leegood 1984). Incidentally, in a study of the PEPCK subtype *Spartina anglica,* Hubb, Smith and Woolhouse (1983) concluded that the activities or enolase and phosphoglycerate mutase were insufficient to support interconversion of PEP and 3PGA at the required rate in either the BSC or the MC. Thus, a relatively low concentration of PEP and a low capacity for interconversion of PEP and 3PGA probably place an upper limit on flux that can be supported by a PEPCK carboxylation cycle in high light (see also Medeiros et al. 2022 for discussion). However, under low irradiation, they could support a considerably larger contribution of PEPCK to the CCM.

### 5. Perturbation and adjustment of $C_{BSC}$

As discussed in the main text, changes in irradiance might result in a temporary imbalance between the concentration of $CO_2$ by the CCM and its utilization by Rubisco (Furbank et al. 1990; von Caemmerer 2000; Kromdijk et al. 2014). An excess of $CO_2$ influx over $CO_2$ consumption will lead to an increase in $C_{BSC}$ and might increase back-leakage of $CO_2$ to the MC whilst an excess of $CO_2$ consumption over $CO_2$ influx will lead to a decrease in $C_{BSC}$ and an increase in the rate of RuBP oxygenation relative to RuBP carboxylation. Both increased photorespiration and increased $CO_2$ back-leakage will decrease photosynthetic efficiency and could contribute to loss of photosynthetic efficiency after a change in irradiance. Measurements of the rate of photorespiration and of $C_{BSC}$ are complicated and challenging (Ubierna et al. 2011; Sage 2014; Kromdijk et al. 2014), especially in non-steady state conditions. Kubacek et al. (2013) that growth in fluctuating light led to increased back-leakage but could not determine in which part(s) of the fluctuating regime this occurred. Recently Wang et al. (2022) used a tunable diode laser absorption spectroscope to monitor delta$^{13}$C in combination with gas exchange to track back-leakage during a dark-light transition in maize and sorghum. For technical reasons, the measurements were performed about 800 ppm $CO_2$ After illuminating maize, estimated leakiness increased progressively to a maximum at about 150 s before declining 2-fold to a steady state value over the next 1200 s. A slower increase in leakiness followed by a slow decline were reported for maize. This was qualitatively consistent with earlier modelling of leakiness during induction (Wang et al. 2021). However, whilst the time resolution was about 10 s, the error associated with determination of delta$^{13}$C was >50% in the first 100 s after illumination indicating that the measurements in this time may he been masked by instrument error.

We asked whether our metabolite data might allow detection of changes in $C_{BSC}$ with shorter time resolution. Measurements of 2PG provide a qualitative proxy for the rate of RuBP oxygenation, and the relationship between the RuBP/2PG ratio and the RuBP/3PGA ratio provides a qualitative proxy for the relative rates of RuBP oxygenation and RuBP carboxylation. Thus, these metabolic traits provide proxy information about $C_{BSC}$. However, it should be stressed that they do not provide unambiguous information because the levels of these metabolites are also affected by other events. The level of 2PG will also depend on how quickly it is degraded by 2-phosphoglycolate phosphatase, and the level of 3PGA (and hence the RuBP/3PGA ratio) will depend on the rate of 3PGA reduction, which in turn depends on the level of triose-P and on the availability of ATP and NADPH.

Immediately following a decrease in irradiance, there was a rapid and significant >2-fold decrease in the level of 2PG (Fig. 3A; Supplementary Fig. S3B), whilst the RuBP/3PGA ratio declined and the RuBP/2PG ratio rose. Conversely, following an increase in irradiance, there was a significant increase of 2PG (Fig. 4A; Supplementary Fig. S4B), and non-significant increase of the RuBP/3PGA ratio and decline of the RuBP/2PG ratio (Fig. 4B; Supplementary Fig. S4B). In their study with fluctuating light, Sales et al. (2025) observed that 2PG increased 30 s after shifting to high light, and decreased 10 and 30 s after shifting to low light.

These early responses point to an increase in carboxylation relative to oxygenation immediately after a decrease in irradiance, and a transient decrease in carboxylation relative to oxygenation immediately after an increase in irradiance. The former may transiently enhance and the latter transiently decrease photosynthetic efficiency, although the impact of the latter may be small due to the relatively low rate of oxygenation relative to carboxylation in $C_4$ photosynthesis. The observed significant changes in 2PG and opposed trends of the RuBP and RuBP/2PG ratios are also indicative of an increase of $C_{BSC}$ after a decrease in irradiance, and an increase of $C_{BSC}$ after a decrease in irradiance. This would result in transiently increased and decreased back-leakage of $CO_2$, respectively.

Back-leakage cannot be directly measured but is instead modelled in various ways, each involving several assumptions (Ubierna et al. 2011; Sage 2014; Kromdijk et al. 2014). Estimated values of back-leakage (expressed as phi) are usually in the range of 0.2-0.3 of the C fixed by PEPC (Hatch et al.1995; Kromdijk et al. 2014; Wang et al. 2024) but depend on the conditions, for example, may increase in low irradiance (Evans et al. 1986; Kromdijk et al. 2014), and can be as high as 0.6-0.9 after a sudden decrease in irradiance (Cousins et al. 2006, 2008; Tazoe et al. 2006, 2008; Kromdijk et al. 2008, 2010, 2014; Pengelly et al. 2010). In a recent analysis of $^{13}CO_2$ labelling patterns in stable low irradiance, Medeiros et al. (2022) estimated that flux at PEPC exceeded that at Rubisco, providing independent support for substantial back-leakage in low irradiance. Our metabolite analyses are consistent with the idea that back-leakage increases transiently after a sudden decrease in irradiance and decreases transiently after a sudden increase in irradiance

We also asked if our metabolite analyses provide information about how quickly these initial changes of $C_{BSC}$ are reversed. Later in the ML-LL transition, the 2PG level did not show consistent changes, the RuBP/3PGA ratio rose and then declined and the RuBP/2PG ratio tended to decline. In the LL-ML transition, 2PG level, and the RuBP/3PGA and RuBP/2PG ratios did not show strong or consistent changes. This unclear picture in both transitions may reflect the likelihood that these metabolic traits are influenced by other factors during the gradual adjustment to a change in irradiance (see above).

Another approach to obtain information about $C_{BSC}$ is to compare $A_n$ with RuBP levels, under the assumption that they should correlate unless further factors are influencing Rubisco activity. Such factors would include Rubisco activation (see above) but also $C_{BSC}$. After a shift to lower irradiance, the 7% recovery of $A_n$ in the recovery phase from about 250 s on was accompanied by a large and significant decline in RuBP levels (Figs. 3A, 5B; Supplementary Figs. S3B, S7A). This recovery of $A_n$ and decline of RuBP was accompanied by a significant increase in the level of CCM metabolites like pyruvate and alanine as well as summed CCM metabolites (Figs. 3A, 3C, 5B; Supplementary Figs. S3B, S7C see also Discussion section of the manuscript). These observations are consistent with a decline of $C_{BSC}$ at the trough of $A_n$

at about 250 s, followed by a rise of $C_{BSC}$ as CCM operation is optimized. After a shift to higher irradiance, from about 120 s onwards $A_n$ rose (from about 86 to 140 nmol $CO_2$ $g^{-1}$ FW $s^{-1}$) without any further increase in RuBP levels (Figs. 4A, 5C; Supplementary Figs. S4B, S8A), consistent with a gradual increase in $C_{BSC}$. This explanation is supported by the observation that the increase of $A_n$ was accompanied by a rise in the levels of individual CCM metabolites including PEP, pyruvate, alanine and aspartate and the summed pool size of CCM metabolites (Figs. 4A, 4C, 6C; Supplementary Figs. S4B, S8C). Incidentally, this predicted slow increase in $C_{BSC}$ from about 120 s on contrasts with the decline in back-leakage (and by inference $C_{BSC}$) after 150 s reported by Wang et al. (2024) after a dark-light transition (see above). This may reflect differences in the speed of activation of the CCM and CBC in these different transitions, or the use of elevated $CO_2$ in the study of Wang et al. (2024)

### 6. Sequestration or recycling of C from metabolite pools in the photorespiratory pathway does not make a large contribution in light transients

Further downstream in the photorespiration pathway, the glycine pool showed marked changes in transitions to lower or to higher irradiance, but there were no consistent changes in serine or glycerate (Figs. 3A, 4A). In the ML-LL transition, glycine decreased by 125 nmol $g^{-1}$ FW between 60 and 600 s (Fig. 3A), equivalent to additional release of $CO_2$ of about 62 nmol $g^{-1}$ FW, or about 1.5 s of photosynthesis at the rate sustained in LL. In the LL-ML transition, glycine increased by about 150 nmol $g^{-1}$ FW between 60 and 1200 s (Fig. 4A). equivalent to a reduction in $CO_2$ release in the photorespiratory pathway of about 75 nmol $g^{-1}$ FW, or only 0.5 s of photosynthesis in ML. Thus, in both cases, changes in the glycine pool can only make a very minor contribution to the delay in reaching the final $A_n$. The same holds for NADH released during glycine decarboxylation as a potential source of energy. Incidentally, in $C_3$ plants some C may exit the photorespiration pathway as glycine or, especially, serine (Fu et al. 2023). If this occurs in $C_4$ plants, this might further dampen any impact of photorespiratory metabolism during light transitions. However, any impact of photorespiration on transition in $C_4$ plants is small. Arce-Cubas et al. (2023a) measured response in phylogenetically coupled pairs of $C_3$ and $C_4$ species to step-changes from darkness to ML or HL in 21% and 2% $O_2$, and Arce Cubas et al (2023b) measured responses to fluctuations in 21% and 2% $O_2$. In all cases, low $O_2$ had a large impact on the response in $C_3$ species, in particular by decreasing photorespiration and the associated post-illumination burst of $CO_2$, but only a small impact in $C_4$. species.

### Additional references


Ballicora, M.A., Iglesias, A.A., Preiss, J. (2004) ADP-glucose pyrophosphorylase; a regulatory enzyme for plant starch synthesis. Photosynth Res 79, 1-24. https://doi.org/10.1023/B:PRES.0000011916.67519.58

Braütigam, A., Schlüter, U., Lundgren, M.R., et al. (2018) Biochemical mechanisms driving rapid fluxes in $C_4$ photosynthesis. bioRxiv 387431; doi: https://doi.org/10.1101/387431

Chao, Q., Liu, X.Y., Mei, Y.C., Gao, Z.-F., Chen, Y.B., Qian, C.R., Hao, Y.B., Wang, B.C. (2014) Light-regulated phosphorylation of maize phosphoenolpyruvate carboxykinase plays a vital role in its activity. Plant Mol Biol 85, 95–105 https://doi.org/10.1007/s11103-014-0171-3

Cousins, A.B., Badger, M.R., von Caemmerer, S. (2006) Carbonic anhydrase and its influence on carbon isotope discrimination during $C_4$ photosynthesis. Insights from antisense RNA in *Flaveria bidentis*. Plant Physiol 141, 232-242. https://doi.org/10.1104/pp.106.077776

Cousins, A.B., Badger, M.R., von Caemmerer, S. (2008) $C_4$ photosynthetic isotope exchange in NAD-ME- and NADP-ME-type grasses. J Exp Bot 59, 1695-1703. https://doi.org/10.1093/jxb/ern001

Evans, J.R., Sharkey, T.D., Berry, J.A., Farquhar, G.D. (1986) Carbon isotope discrimination measured concurrently with gas exchange to investigate $CO_2$ diffusion in leaves of higher plants. Aust J Plant Physiol 13, 281-292. https://doi.org/10.1071/PP9860281

Fukuyama, H., Masumoto, C., Taniguchi, Y., Baba-Kasai, A., Katoh, Y., Ohkawa, H., Mitsue Miyao,M. (2015) Characterization and expression analyses of two plastidic enolase genes in rice. Biosci Biotechnol Bioche. 79, 402-409. https://doi.org/10.1080/09168451.2014.980219

Furumoto T, Hata S, Izui K (1999) cDNA cloning and characterization of maize phosphoenolpyruvate carboxykinase, a bundle sheath cell-specific enzyme. Plant Mol Biol 41(3):301–311. doi.org/10.1023/A:1006317120460

Hatch, M.D., Kagawa, T. (1976) Photosynthetic activities of isolated bundle sheath cells in relation to differing mechanisms of C4 pathway photosynthesis. Arch Biochem Biophys 175: 39–53. DOI: 10.1016/0003-9861(76)90483-5

Kromdijk, J., Schepers, H.E., Albanito, F., et al. (2008) Bundle sheath leakiness and light limitation during $C_4$ leaf and canopy $CO_2$ uptake. Plant Physiol 148, 2144-2155. https://doi.org/10.1104/pp.108.129890

Kromdijk, J., Griffiths, H., Schepers, H.E. (2010) Can the progressive increase of $C_4$ bundle sheath leakiness at low PFD be explained by incomplete suppression of photorespiration? Plant Cell Environ 33, 1935-1948. https://doi.org/10.1111/j.1365-3040.2010.02196.x

Pick, T.R., Brautigam, A., Schluter, U., Denton, A.K., Colmsee, C., Scholz, U., Fahnenstich, H., Pieruschka, R., Rascher, U., Sonnewald, U., and Weber, A.P. (2011). Systems analysis of a maize leaf developmental gradient redefines the current $C_4$ model and provides candidates for regulation. Plant Cell 23, 4208-4220. DOI: 10.1105/tpc.111.090324

Portis, A.R. Jr., Li, C., Wang, D., Salvucci, M.E. (2008) Regulation of Rubisco activase and its interaction with Rubisco. J Exp Bot 59, 1597-1604. https://doi.org/10.1093/jxb/erm240

Portis, A.R. Jr., Parry, M.A. (2007) Discoveries in Rubisco (ribulose 1,5-bisphosphate carboxylase/oxygenase): a historical perspective. Photosynth Res 94, 121-143. https://doi.org/10.1007/s11120-007-9225-6

Sage, R.F. (2014) Stopping the leaks: new insights into $C_4$ photosynthesis at low light. Plant Cell Environ 37, 1037–1041. https://doi.org/10.1111/pce.12246

Scheibe, R., aStitt, M. (1988) Comparison of NADP-malate dehydrogenase activation, QA reduction and $O_2$ evolution in spinach leaves. Plant Physiol Biochem 26, 473-481. https://api.semanticscholar.org/CorpusID:82024560

Smith, A., Woolhouse, H. W. (1983) Metabolism of phosphoenolpyruvate in the C4 cycle during photosynthesis in the phosphoenolpyruvate-carboxykinase C4 grass Spartina anglica Hubb. Planta 159, 570.578.DOI: 10.1007/BF00409147.

Stitt, M., ap Rees, T.A. (1980) Carbohydrate breakdown by chloroplasts of *Pisum sativum*. Biochim. Biophys. Acta 627, 131-143. https://doi.org/10.1016/0304-4165(80)90315-3

Stryer, L. (1990) Biochemistry. Springer, Heidelberg

Szecowka, M., Heise, R., Tohge, T., et al. (2013) Metabolic fluxes of an illuminated *Arabidopsis thaliana* rosette. Plant Cell 25, 694-714. https://doi.org/10.1105/tpc.112.106989

Tazoe, Y., Noguchi, K., Terashima, I. (2006) Effects of growth light and nitrogen nutrition on the organization of the photosynthetic apparatus in leaves of a $C_4$ plant, *Amaranthus cruentus*. Plant Cell Environ 29, 691-700. https://doi.org/10.1111/j.1365-3040.2005.01453.x

Ubierna, N., Sun, W., Cousins, A.B. (2011) The efficiency of $C_4$ photosynthesis under low light conditions: assumptions and calculations with $CO_2$ isotope discrimination. J Exp Bot 62, 3119-3134. https://doi.org/10.1093/jxb/err073

Wang, L., Czedik-Eysenberg, A., Mertz, R.A., et al. (2014c). Comparative analyses of $C_4$ and $C_3$ photosynthesis in developing leaves of maize and rice. Nature Biotechnology 32, 1158-1165. doi.org/10.1038/nbt.3019

Yokochi, Y., Yoshida, K., Hahn, F., Miyagi, A., Wakabayashi, K.I., Kawai-Yamada, M., Weber, A.P.M., Hisabori, T. (2021) Redox regulation of NADP-malate dehydrogenase is vital for land plants under fluctuating light environment. Proc Natl Acad Sci U S A. 118: e2016903118. doi: 10.1073/pnas.2016903118
